# Development of pyrazolo[1,5-a]pyrimidine based macrocyclic kinase inhibitors targeting AAK1

**DOI:** 10.1101/2025.04.02.646796

**Authors:** Theresa E. Mensing, Christian G. Kurz, Jennifer A. Amrhein, Theresa A. L. Ehret, Franziska Preuss, Sebastian Mathea, Marwah Karim, Do Hoang Nhu Tran, Zuzana Kadlecova, Tuomas Tolvanen, Daniel Martinez-Molina, Susanne Müller, Shirit Einav, Stefan Knapp, Thomas Hanke

**Author notes:** Correspondence (S.K.); (T.H.).

## Abstract

Since the outbreak of SARS-CoV-2 in recent years, our society has become more aware that zoonotic diseases pose a real threat. Therefore, the demand for small molecules that target host proteins, essential for viral entry and replication, has increased as an interesting strategy for the development of antiviral agents, as these agents may be effective against several different pathogens. NAK kinases is one such potential target family because they are involved in a variety of cellular functions, hijacked by viruses to invade host cells, such as clathrin-mediated endocytosis. A large number of different inhibitors have already been reported targeting NAK kinases, but there are still no compounds that selectively target AAK1 over other NAK family members, in particular the closely related family member BIKE. Here, we developed a series of pyrazolo[1,5-a]pyrimidine-based macrocyclic NAK inhibitors, starting from the acyclic AAK1 inhibitor LP-935509. Through a structure-guided activity relationship study within the NAK family, we identified potent AAK1 inhibitors **16**, **18** and **27**, which show promising selectivity within the NAK family. The inhibitors showed a potent inhibition of the phosphorylation of the AP-2 complex and the antiviral activity of the compounds was evaluated against various RNA viruses.

---

Zoonotic diseases have accompanied humans since time immemorial, with rabies being the best-known example. More recently, coronaviruses have come to the fore. Currently more than 200 different viruses are known to cause diseases in humans. The antiviral agents currently available mostly target enzymes essential viral proteins and these agents are therefore referred to as direct-acting antivirals (DAAs). One of the problems with this approach to antiviral therapy is that the development of such individual drugs is time-consuming and cost-intensive. In addition, these antiviral agents offer little flexibility for the treatment of variants of these viruses that may arise quickly or novel zoonotic diseases. Host-targeted antivirals (HATs) are therefore playing an increasingly important role in the development of new antiviral agents.(1)(2, 3) One of the most common host mechanisms used by viruses to enter host cells is the process of clathrin-mediated endocytosis (CME) (**Figure 1A**).(4) Small to medium-sized viruses, such as the dengue virus (DENV) or the Venezuelan equine encephalitis virus (VEEV), have been described to highjack this mechanism.(5–7) CME is one of the most important endocytic signalling pathways in mammalian cells and regulates processes such as signal transduction at the cell surface, the uptake of transmembrane receptors and the remodelling of the plasma membrane.(8) The CME mechanism is controlled by a complex network of protein interactions that facilitates vesicular transport in clathrin-coated-vesicles (CCV) from the cell membrane to the endosome. One part of this protein complex network is the adaptor protein-2-complex (AP2), which plays a central role in all phases of CME. By recruiting clathrin to form CCVs, the active AP2 adaptor complex regulates CME initiation. Various kinases, such as AAK1 (adaptor associated kinase 1), BIKE (BMP-2-inducible kinase) and GAK (cyclin G-associated kinase), phosphorylate the µ2 subunit of the AP2 adaptor complex, which leads to an activation.(9–14) This phosphorylation process begins with the formation of the clathrin-coated pit (CCP) and intensifies as the CCP matures. It triggers a conformational change in the AP2 adaptor complex that enables it to bind cargo proteins and to form a platform for the attachment of endocytic NECAP proteins. Recruitment of NECAP proteins is enhanced by phosphorylation of the µ2 subunit. As a result, NECAP contributes to the recruitment of proteins that facilitate the transition of CCPs to CCVs, such as SNX9 (**Figure *6*A**).(15–17)

**Figure 1:**
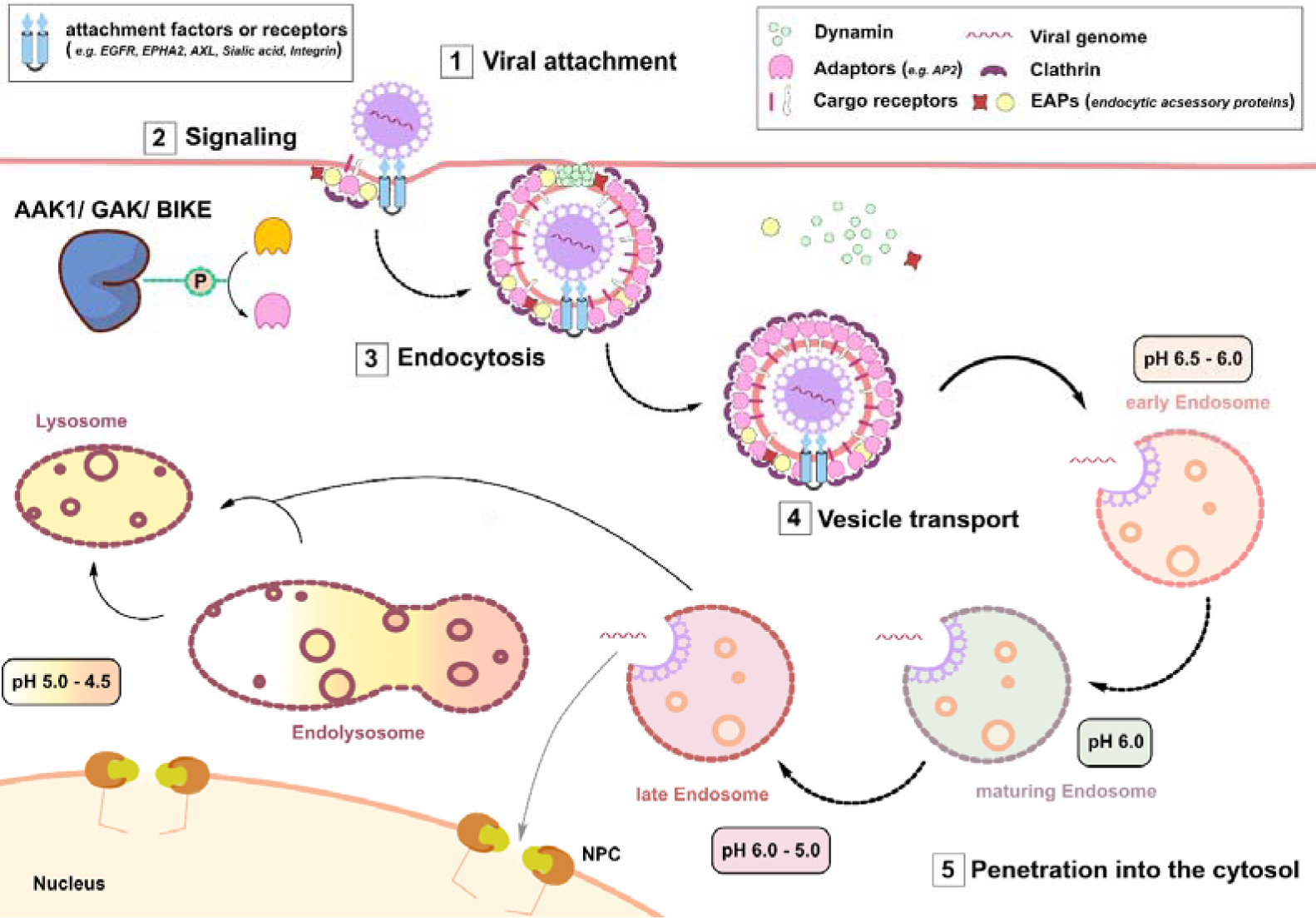
Mechanism of viral entry via clathrin-mediated-endocytosis (adapted from Robinson et al. and Yamauchi et al.).(18, 19) Viral attachment occurs through the binding to cell surface structures e.g. proteins, lipids or receptors, able to actively promote endocytosis of the virus. Following attachment, viruses commonly activate signalling systems of the cell for viral entry. This internalization process is dependent on clathrin and clathrin adaptors (e.g. AP2). AP2 activation is mediated through phosphorylation of its µ-subunit by kinases such as AAK1, GAK and BIKE. Viral particle-bearing clathrin-coated-vesicles (CCV) are transported to and through endosomal compartments, releasing the viral genome, which then penetrates the cytoplasm.

AAK1, BIKE and GAK are serine/threonine kinases and belong to the family of Numb-associated protein kinases (NAK). The fourth and last member of this family is the myristoylated and palmitoylated serine/threonine kinase 1 (MPSK1), also known as STK16. The members of the NAK family share about 30% of sequence identity in their kinase domain, while the rest of the protein has less similarity. The most closely related members of the NAK family are AAK1 and BIKE with a total sequence identity of 50%. In their kinase domains, this identity increases to 74%. With a sequence identity of 39% and 30% respectively, GAK and MPSK1 are the least related to AAK1.(20) This indicates that it is easier to achieve selectivity for AAK1 over GAK and MPSK1, but not over BIKE. To date, several potent NAK inhibitors have been developed. The chemical probe **SGC-GAK-1** is a 4-anilinoquinoline-based GAK inhibitor with a 16LJ000-fold selectivity compared to the other three members of the NAK family (**Figure 2**).(21) In contrast, attempts to develop inhibitors with a selectivity for AAK1 or BIKE proved to be more difficult. Most AAK1 inhibitors are in fact dual AAK1 and BIKE inhibitors, such as the chemical probe **SGC-AAK1-1** or the arylamide-based compound **BMS-911172** (**Figure 2**).(21, 22) Due to the highly conserved structure within the kinase domain of both kinases, selective inhibition of one kinase without simultaneously targeting the other is extremely challenging.(23, 24, 22, 21)

**Figure 2:**
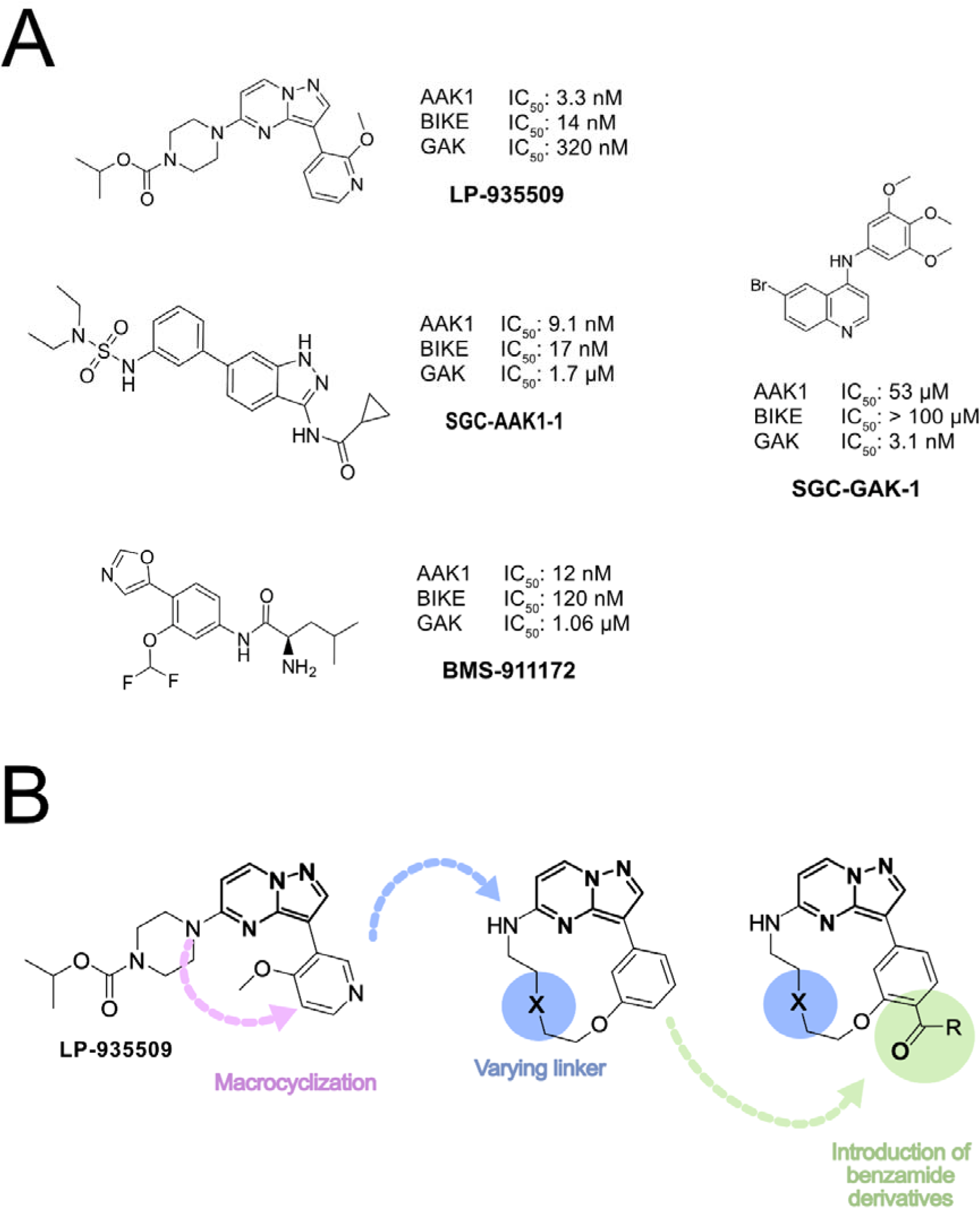
Small Molecule Kinase Inhibitors targeting AAK1 and BIKE. **A:** Chemical structures of NAK kinase inhibitors LP-935509, SGC-AAK1-1, BMS-911172, SGC-GAK-1 and their corresponding IC_50_ values on AAK1, BIKE and GAK**. B:** Design and synthetic strategy of macrocyclic NAK-inhibitors, based on the acyclic NAK inhibitor LP-935509.

AAK1, BIKE and GAK have been shown to be important regulators of CME by initiating the formation of CCVs, a cellular mechanism that is often hijacked by various RNA viruses.(9–14) In addition, AAK1 and GAK were found to regulate the spread and intracellular trafficking of hepatitis C virus (HCV) from cell to cell by controlling clathrin-associated adaptor proteins (APs).(25, 2, 26) Therefore, the interest in their potential to achieve a broad antiviral effect has increased, especially during the SARS-CoV-2 pandemic.(27, 26, 14) The approved kinase inhibitors erlotinib and sunitinib showed strong antiviral activity against members of the Flaviviridae family (e.g. DENV, Hepatitis C virus) as well as unrelated families of RNA viruses through the potent NAK inhibition.(2, 28–32, 26, 33–36) In addition, *Pu et al.* showed that AAK1 is overexpressed in DENV-infected cells, while non-infected cells from the same culture have physiologically lower protein levels.(37) A correlation was observed between expression levels and cellular virus abundance, highlighting AAK1 as an even more interesting target for viral diseases.(37)

As all members of the NAK family are involved in the regulation of CME, the development of selective inhibitors targeting only one family member would provide an opportunity to further investigate their specific roles in virus entry. To date, potent and selective GAK and MPSK1 inhibitors are available, while few inhibitors are available for AAK1 and BIKE. Reported inhibitors show usually dual activity for AAK1 and BIKE or have also GAK activity (**Figure *2***).(38, 21) The obstacles to the development of a selective AAK1 or BIKE inhibitor are mainly due to the sequence homology in the kinase domain of AAK1 and BIKE.(20),(23) Although selective GAK inhibitors have already been developed, it is a common off-target for many kinase inhibitors. *Chaikuad et al.* reported using structural analysis of GAK in its active and inactive states, that the kinase domain of GAK exhibits high degree of plasticity, suggesting that many different kinase inhibitors can bind to GAK recognizing a variety of conformations.(39)

Macrocyclization of linear compounds has proven to be a versatile approach to improve the selectivity of an inhibitor while maintaining or even increasing its potency.(40, 41) The conformational flexibility of linear molecules can have a negative impact on their selectivity profile, as the binding mode can vary for different kinases. Macrocyclization results in a pre-organized structure with a more restricted conformation. This minimizes entropic costs and ideally forces the pharmacophore into an active and selective conformation.(42) Using this macrocyclization strategy, we have recently identified selective ligands for e.g. MST3 or EPHA2.(43),(44)

**LP-935509** (**Figure 2A**) is a potent, acyclic and ATP-competitive inhibitor of AAK1 (K_D_: 3 nM) and BIKE (K_D_: 14 nM), while it only moderately inhibits GAK (IC_50_: 320 nM).(45) The compound has been tested in several preclinical animal models where it showed efficacy reducing neuropathic pain.(45) In SH-SY5Y cells, **LP-935509** significantly inhibited the AP2M1 (AP2-complex µ-subunit) phosphorylation at Thr156, while it showed a dose-dependent inhibition of SARS-CoV-2 S-RBD (receptor binding domain of the viral spike protein) internalization in host cells.(46, 47) Based on its potent inhibition of NAK kinases, we hypothesized that **LP-935509** represents a promising lead structure for the development of compounds with an improved selectivity profile for AAK1 over BIKE. We aimed to achieve this goal by macrocyclization and derivatization of the linear inhibitor **LP-935509** (**Figure 2B**). We established a SAR based on a macrocyclic pyrazolo[1,5-a]pyrimidine scaffold (**Figure 2B**), focusing on the back pocket where we introduced different benzylamide derivatives as well as different linkers. The selectivity profile of the synthesized inhibitors was tested against a representative panel of 102 human kinases. Inhibitors with the most promising potency and selectivity were tested in Western blots for cellular activity on AP-2 complex phosphorylation and on viral infection of different cell lines.

## Results and Discussion

### Benzylamide Derivatization leads to Selectivity between AAK1 and BIKE

The macrocyclic compounds **14**–**30** were synthesized according to **Scheme 1**. Starting from chlorinated pyrazolo[1,5-a]pyrimidine **1**, bromination with NBS led to compound **2** with a yield of 96%. Nucleophilic substitution with 6-aminohexan-1-ol or 2-(2-aminoethoxy)ethan-1-ol, gave the corresponding intermediates **3a** and **3b** with yields of 63% to 81%, respectively. TBDMS-protection of the primary alcohol (75% for **4a**, 99% for **4b**) and subsequent Boc-protection (95% for **5a** and 90% for **5b**) provided the starting material for Suzuki cross-coupling. To generate the pinacol boronic ester, required for the Suzuki cross-coupling, 4-bromo-2-hydroxybenzoic acid **6** was esterified in methanol, under acidic conditions (yield: 46%) and then borylated by Miyaura borylation, with a yield of 81%, to **8**. Suzuki cross-coupling of **8** with compounds **5a** and **5b**, yielded the intermediates **9a** (91%) and **9b** (83%), respectively. Subsequent desilylation with TBAF produced the Mitsunobu reaction precursors **10a** and **10b** in yields between 85–91%. An intramolecular Mitsunobu reaction was performed at high temperatures and high dilution to obtain the macrocyclic compounds **11a** (yield: 87%) and **11b** (yield: 90%).

**Scheme 1:**
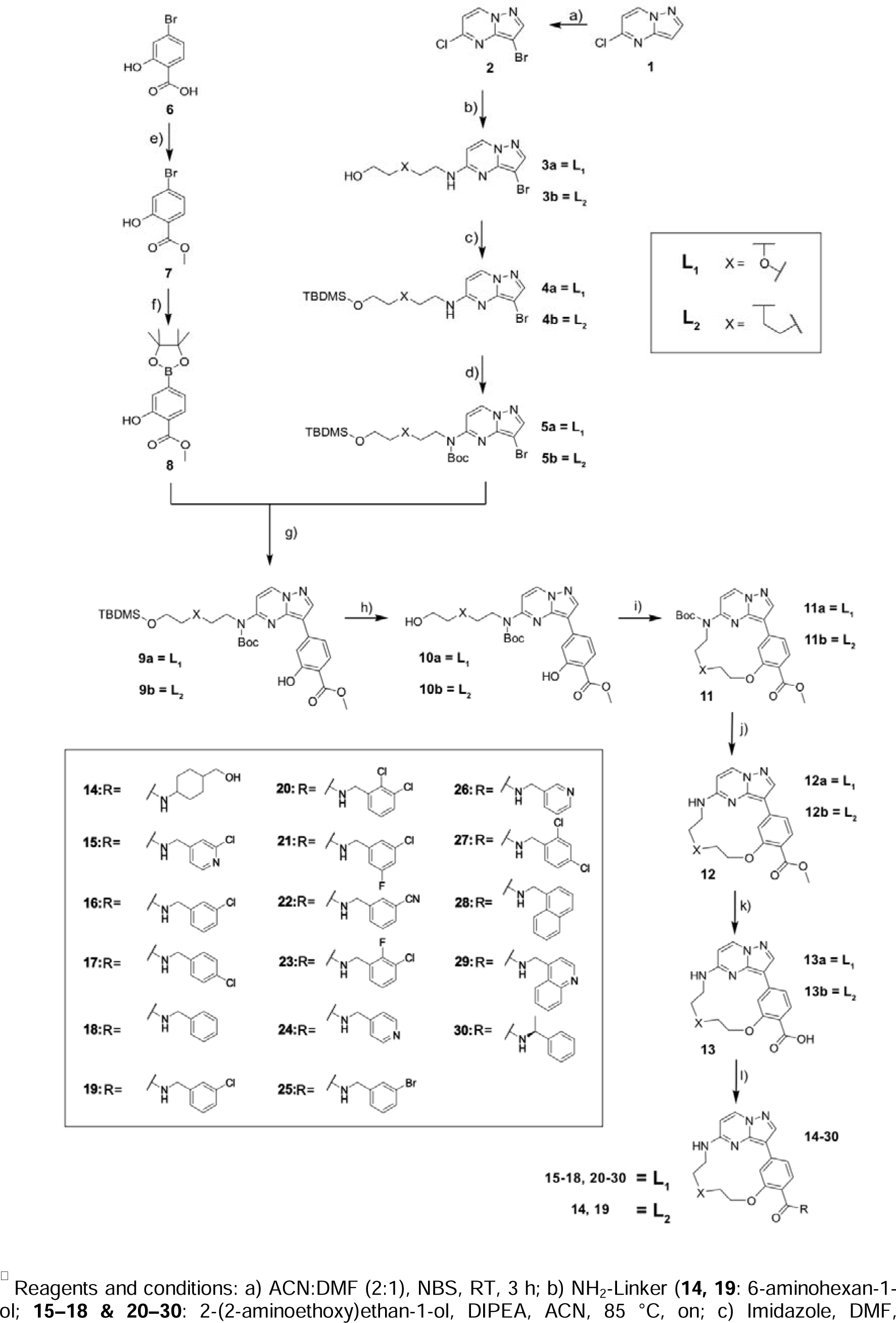
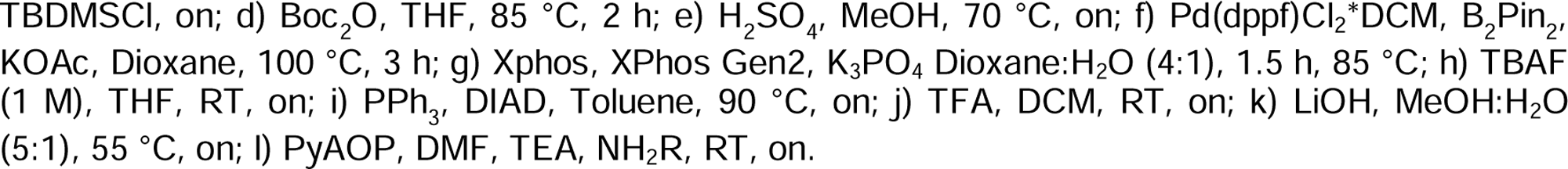
Synthesis of macrocyclic pyrazolo[1,5-α]pyrimidines **14**-**31**

Subsequent Boc-deprotection with TFA and ester hydrolysis with LiOH yielded the basic macrocyclic structures **12a** and **12b** bearing the carboxylic function required for derivatization. The introduction of various substituents into the macrocyclic scaffold was carried out via amide coupling after activation of the carboxylic acid with PyAOP and led to compounds **14–30**.

To investigate the ability of the compounds to bind to kinases of the NAK-family, a differential scanning fluorimetry (DSF) assay was performed with AAK1, BIKE and GAK (**Table 1**). In this assay, the specific thermal shift of a protein (ΔT_m_) is compared with the thermal shift exhibited by this protein after incubation with the test compounds. An increased ΔT_m_-shift leads to stabilization of the protein and indicates binding of the test compound to the protein.(48) The promiscuous kinase inhibitor staurosporine, **LP-935509**, the chemical probes **SGC-AAK1-1**, **SGC-GAK-1** and the corresponding negative controls (**SGC-AAK1-1N**, **SGC-GAK-1N**) were used as control compounds.(49, 21, 45) Since the absolute values of the thermal shift are different for the individual kinases, the ΔT_m_-value for each compound is shown as a percentage shift compared to the reference compound staurosporine (**Table 1**). Staurosporine is a potent binder with comparable K_d_ values for all three members of the NAK family, but it has different ΔT_m_ shifts for each of these kinase (ΔT_m_-shifts^1^: AAK1: 14 K, K_d_: 1.2 nM(50), BIKE: 18 K, K_d_:3.7 nM(51); GAK: 7.7 K, K_d_: 17 nM).(21, 49)

**Table 1:**
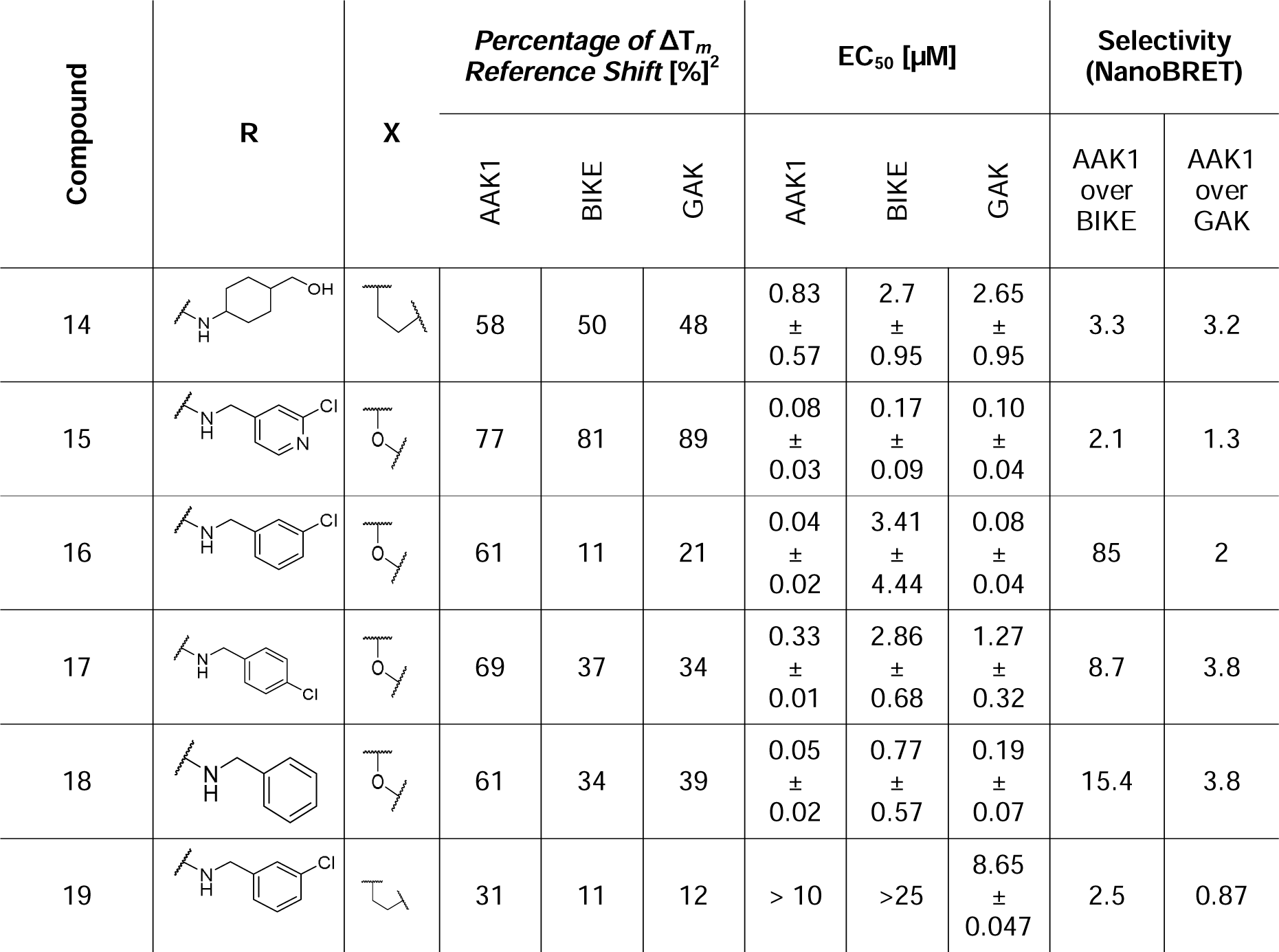
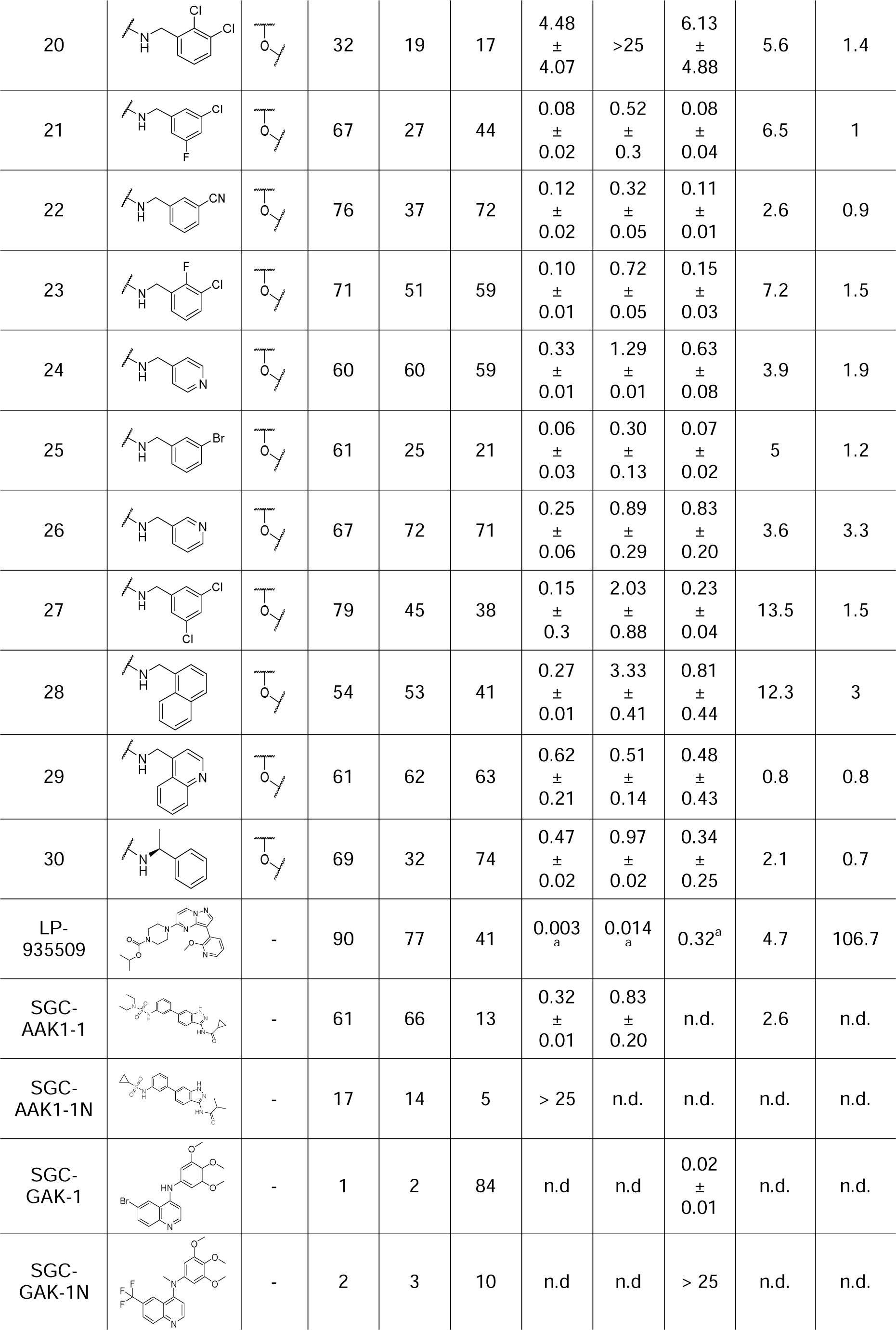
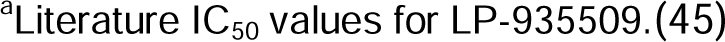
Profiling of the respective derivatives and the control compound LP-935509(45) within members of the NAK family. Determination of EC_50_ values in a cellular NanoBRET target engagement assay, as well as protein stabilization and selectivity scores via thermal shift assay.

| Compound | R | X | Percentage of $\Delta T_m$<br>Reference Shift [%]<sup>2</sup> | | | EC<sub>50</sub> [μM] | | | Selectivity<br>(NanoBRET) | |
| --- | --- | --- | --- | --- | --- | --- | --- | --- | --- | --- |
|  |  |  | AAK1 | BIKE | GAK | AAK1 | BIKE | GAK | AAK1<br>over<br>BIKE | AAK1<br>over<br>GAK |
| 14 |  |  | 58 | 50 | 48 | 0.83<br>±<br>0.57 | 2.7<br>±<br>0.95 | 2.65<br>±<br>0.95 | 3.3 | 3.2 |
| 15 |  |  | 77 | 81 | 89 | 0.08<br>±<br>0.03 | 0.17<br>±<br>0.09 | 0.10<br>±<br>0.04 | 2.1 | 1.3 |
| 16 |  |  | 61 | 11 | 21 | 0.04<br>±<br>0.02 | 3.41<br>±<br>4.44 | 0.08<br>±<br>0.04 | 85 | 2 |
| 17 |  |  | 69 | 37 | 34 | 0.33<br>±<br>0.01 | 2.86<br>±<br>0.68 | 1.27<br>±<br>0.32 | 8.7 | 3.8 |
| 18 |  |  | 61 | 34 | 39 | 0.05<br>±<br>0.02 | 0.77<br>±<br>0.57 | 0.19<br>±<br>0.07 | 15.4 | 3.8 |
| 19 |  |  | 31 | 11 | 12 | > 10 | >25 | 8.65<br>±<br>0.047 | 2.5 | 0.87 |

<sup>2</sup> Staurosporine was used as Reference Compound to determine the respective percentage Shifts for each kinase

|  |  |  |  |  |  |  |  |  |  |  |
| --- | --- | --- | --- | --- | --- | --- | --- | --- | --- | --- |
| 20 |  |  | 32 | 19 | 17 | 4.48<br>±<br>4.07 | >25 | 6.13<br>±<br>4.88 | 5.6 | 1.4 |
| 21 |  |  | 67 | 27 | 44 | 0.08<br>±<br>0.02 | 0.52<br>±<br>0.3 | 0.08<br>±<br>0.04 | 6.5 | 1 |
| 22 |  |  | 76 | 37 | 72 | 0.12<br>±<br>0.02 | 0.32<br>±<br>0.05 | 0.11<br>±<br>0.01 | 2.6 | 0.9 |
| 23 |  |  | 71 | 51 | 59 | 0.10<br>±<br>0.01 | 0.72<br>±<br>0.05 | 0.15<br>±<br>0.03 | 7.2 | 1.5 |
| 24 |  |  | 60 | 60 | 59 | 0.33<br>±<br>0.01 | 1.29<br>±<br>0.01 | 0.63<br>±<br>0.08 | 3.9 | 1.9 |
| 25 |  |  | 61 | 25 | 21 | 0.06<br>±<br>0.03 | 0.30<br>±<br>0.13 | 0.07<br>±<br>0.02 | 5 | 1.2 |
| 26 |  |  | 67 | 72 | 71 | 0.25<br>±<br>0.06 | 0.89<br>±<br>0.29 | 0.83<br>±<br>0.20 | 3.6 | 3.3 |
| 27 |  |  | 79 | 45 | 38 | 0.15<br>±<br>0.3 | 2.03<br>±<br>0.88 | 0.23<br>±<br>0.04 | 13.5 | 1.5 |
| 28 |  |  | 54 | 53 | 41 | 0.27<br>±<br>0.01 | 3.33<br>±<br>0.41 | 0.81<br>±<br>0.44 | 12.3 | 3 |
| 29 |  |  | 61 | 62 | 63 | 0.62<br>±<br>0.21 | 0.51<br>±<br>0.14 | 0.48<br>±<br>0.43 | 0.8 | 0.8 |
| 30 |  |  | 69 | 32 | 74 | 0.47<br>±<br>0.02 | 0.97<br>±<br>0.02 | 0.34<br>±<br>0.25 | 2.1 | 0.7 |
| LP-935509 |  | - | 90 | 77 | 41 | 0.003<sub>a</sub> | 0.014<sub>a</sub> | 0.32<sup>a</sup> | 4.7 | 106.7 |
| SGC-AAK1-1 |  | - | 61 | 66 | 13 | 0.32<br>±<br>0.01 | 0.83<br>±<br>0.20 | n.d. | 2.6 | n.d. |
| SGC-AAK1-1N |  | - | 17 | 14 | 5 | > 25 | n.d. | n.d. | n.d. | n.d. |
| SGC-GAK-1 |  | - | 1 | 2 | 84 | n.d. | n.d. | 0.02<br>±<br>0.01 | n.d. | n.d. |
| SGC-GAK-1N |  | - | 2 | 3 | 10 | n.d. | n.d. | > 25 | n.d. | n.d. |

<sup>a</sup>Literature IC<sub>50</sub> values for LP-935509.(45)

The thermal shift assay confirmed that the two reference compounds **LP-935509** and **SGC-AAK1-1** showed no selectivity within the NAK family and that AAK1 and BIKE were stabilized about equally well, which was in agreement with literature values. Both inhibitors exhibited a relatively low stabilization of GAK (41%, 13% resp.), while they displayed strong stabilization of AAK1 and BIKE (AAK1: 90% and 77%, BIKE: 60% and 66% respectively). Of all macrocyclic compounds, compound **27** showed the strongest stabilization of AAK1, reaching 79% of the shift of the reference (staurosporine). Compounds **16**, **21** and **22** showed smaller ΔT_m_-shifts for AAK1 compared to **27**, but the stabilization of AAK1 was significantly larger compared to BIKE (**16**: 61% vs 11%; **21**: 67% vs 27%; **22**: 76% vs 37%; **Table 1**). The SAR revealed that compounds with a substituent in the meta-position on the benzylamide led to increased selectivity for AAK1 over BIKE. However, other substitution patterns, such as compound **24** or **29**, were non-selective and showed approximately the same stabilization for all three kinases (**24**: 60/60/59%; **29**: 61/62/63%; AAK1/BIKE/GAK; **Table 1**). It was also interesting to note that compound **19**, in which only the macrocyclic linker was exchanged compared to compound **16**, led to the weakest stabilization of AAK1 (**19**: 31%; **16**: 61% for AAK1). This indicated that the back-pocket motif was probably no longer able to adapt the bioactive conformation most likely due to the longer linker in compound **19**.

To evaluate the cellular potency of the synthesized compounds, all macrocycles were tested in NanoBRET assays (Promega, Madison, WI, USA). Compounds were profiled against all members of the NAK family (**Table 1**, **Figure 4A**, MPSK1: Supplementary data **Table S3**). Compound **16** revealed to be a potent inhibitor of AAK1, with a cellular EC_50_ of 40 nM. It however bound GAK with comparable potency (80 nM) but only showed negligible activity on BIKE (3.41 µM). In contrast, compound **27,** which showed the highest stabilization of AAK1 in the DSF assay, had only an EC_50_ of 150 nM for AAK1. Compound **18** was a potent binder of AAK1 (EC_50_ of 50 nM) but had only 15.4-fold selectivity over BIKE. In addition, compound **18** had an EC_50_ of 190 nM for GAK. Compound **25** exhibited approximately equal activity on all three NAK members in the NanoBRET assay, with EC_50’s_ of 60 nM, 70 nM and 300 nM for AAK1, GAK and BIKE, respectively. In agreement with the low activity of all three NAK members in the DSF assay, the EC_50_ values of compound **19** in the NanoBRET assay were also all in the high micromolar range. Within this SAR series, macrocycles **15**, **22** and **25** showed the highest potency for BIKE. All three compounds contained a *meta* substituent at the aromatic benzylamide moiety. Surprisingly, the *meta* chlorinated **16** showed a 20-fold lower potency on BIKE compared to **15**. The meta chlorinated pyridine of **15** was tolerated by all three NAK kinases, whereas the chlorinated benzene derivative (**16**) was only tolerated for AAK1 and GAK. Larger substituents in the *meta* position such as bromine and nitrile (**22**, **25**) increased the potency for BIKE compared to the chlorinated compound **16**. Small substituents in the ortho position, such as a fluorine in **23**, were still tolerated by all three kinases (AAK1: 100 nM, BIKE: 720 nM and GAK: 150 nM). In contrast, larger substituents in the ortho position, such as the *ortho*- and *meta*-chlorinated compound **20**, had EC_50_ values in the high micromolar range. Replacement of the linker in compound **19** also resulted in loss of activity for all three kinases compared to compound **16**.

To further evaluate the selectivity profile outside the NAK family, all compounds were screened against a panel of 88 kinases using a DSF assay (Supplementary data **Table S2**). Compounds **16**, **18** and **27** showed the most favorable potency and selectivity profiles of all compounds tested (**Figure *3***). While **16** showed only weak activity on the kinases (FGFR1 and DAPK3), compound **18** stabilized 14 additional kinases with a percentage shift more than 35% compared to the reference compound. Macrocycle **27** showed significant Tm shifts for five additional kinases (FGFR2, GAK, BIKE, MERTK, DAPK3), each with a percentage shift of approximately 35%.

**Figure 3:**
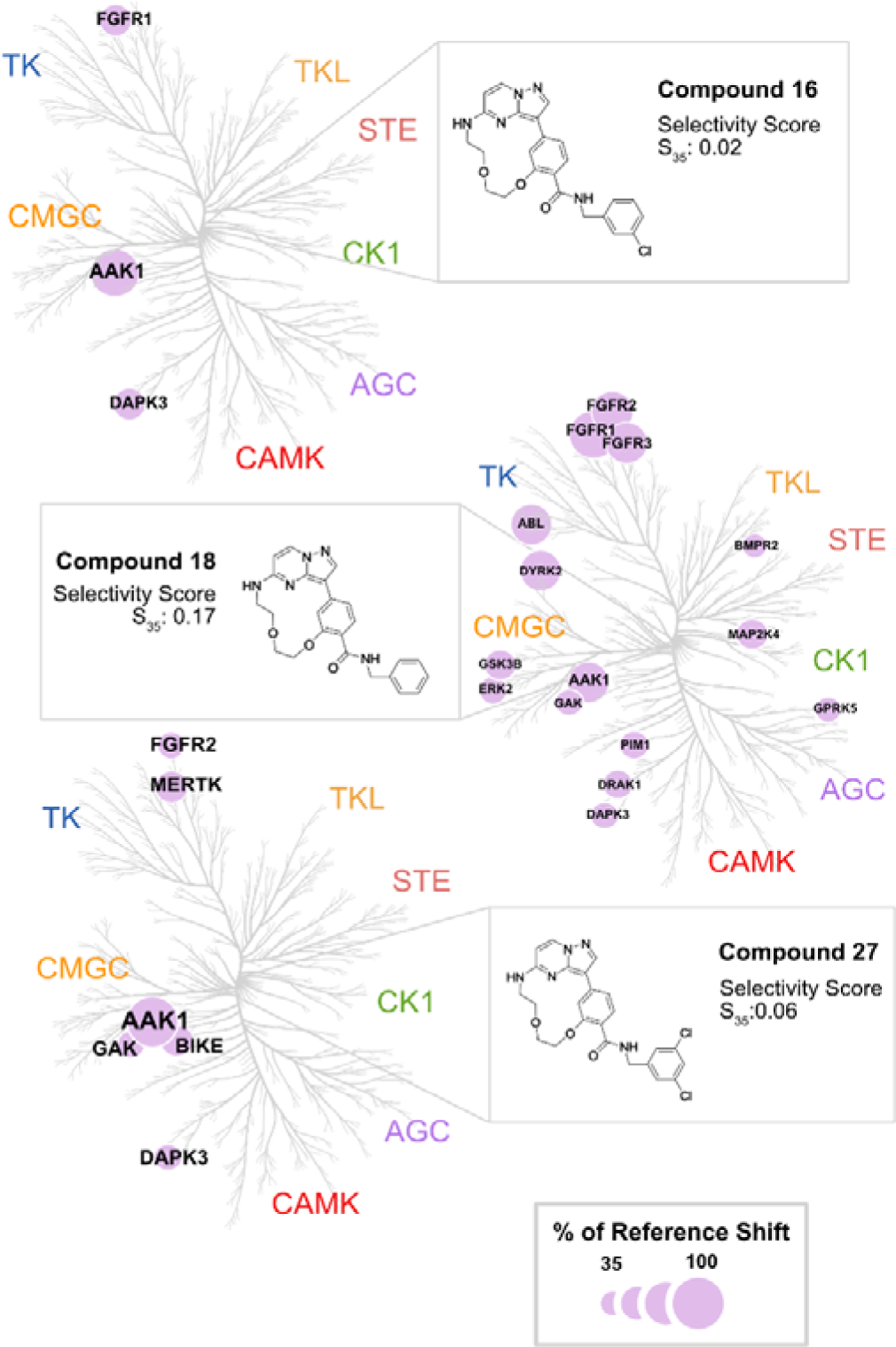
Phylogenetic kinome tree depicting the screened kinases for **16**, **18** and **27** as dots of different size, relative to the thermal shift of a control compound in K set as 100%. Hits above 35% of the Reference shift depicted as violet dots, their size corresponding to the percentage of the measured staurosporine shift. AGC (protein kinase A, G, C), CAMK (calcium/calmodulin-dependent kinases), CK1 (casein kinase 1), CMGC (cyclin-dependent kinases, MAP kinases, glycogen synthase kinases, casein kinase 2), STE (homologues of yeast sterile 7, 11, 20), TK (tyrosine kinases) and TKL (tyrosine kinase-like)-family.

To verify the cellular activity of the compounds, several compounds were tested in a cellular thermal shift assay (CETSA), which also provided information on the cellular selectivity beyond the kinome. Similar to our in vitro DSF assay, CETSA measures the stabilization of a protein in the presence and absence of a ligand in cellular systems and thus evaluates the binding of a compound to all expressed proteins. Macrocycle **16** showed excellent selectivity in cellular CETSA with significant stabilization of AAK1, while no significant stabilization of GAK and BIKE was observed (**Figure 4B**). On the other hand, we observed stabilization of AAK1/BIKE and GAK for macrocycle **24**, which agreed well with our NanoBRET data, where **24** was also found to be non-selective within the NAK family.

**Figure 4:**
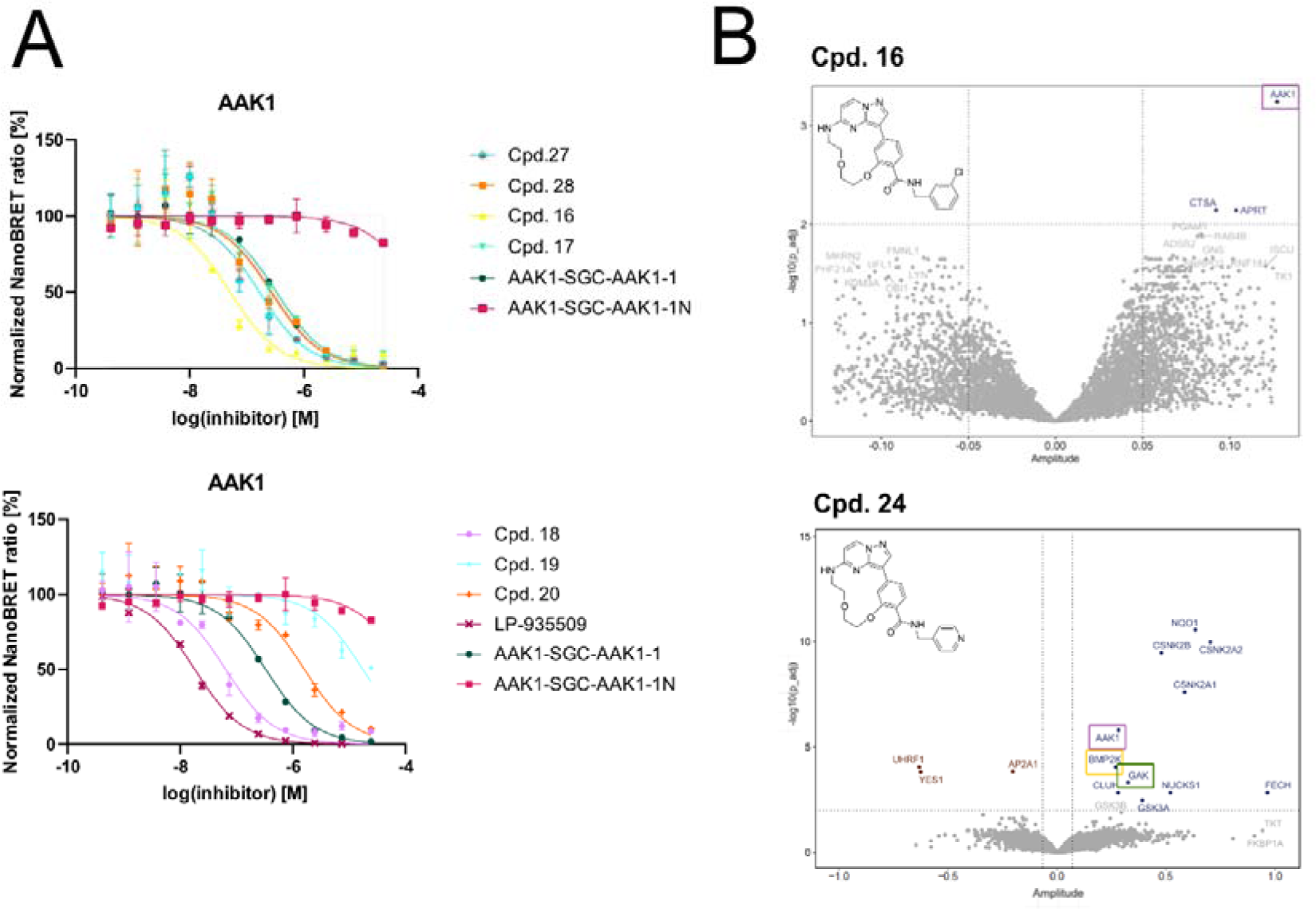
Compound target engagement in cells. **A:** Curves generated via NanoBRET assay for **16, 17, 18, 19, 20, 27, 28, LP-935509, SGC-AAK1-1** and **SGC-AAK1-1N**. The normalized NanoBRET ratio [%] was plotted against the logarithmic function of the increasing inhibitor concentration [M]. The experiments were performed in biological and technical duplicates (N=2). Data is displayed as mean ± SD. **B:** Volcano plots of stabilized (positive amplitude) and destabilized (negative amplitude) proteins, after incubation with compounds **16** and **24.** The Y-axis shows statistical significance, while the X-axis gives the amplitude of the change in melting temperature. Stabilized proteins are depicted in blue, destabilized proteins in red and non-significant proteins are marked grey. Dotted lines represent the cut-off values.

### Structural Insights into the binding mode of Compound **18** with AAK1

The co-crystal structure of **18** with AAK1 (PDB: 9QB5) revealed a similar binding mode as observed for related macrocyclic inhibitors of the core scaffold (**Figure 5A**)(52, 53). The aromatic pyrazolo[1,5-a]pyrimidine nitrogen was responsible for the hinge interaction, while the benzyl moiety of the primary amide attached to the macrocyclic core structure extended towards the glycine rich loop (**Figure 5B**). To identify potential structural reasons for the recurring selectivity for AAK1 and GAK over BIKE, the co-crystal structure of AAK1 with **18** (PDB: 9QB5) was superimposed with structures solved for BIKE and GAK (PDB-ID:4W9W, and 5Y7Z) (**Fig. 4B**). The ATP binding pockets of AAK1 and BIKE differ in only a few amino acids, indicating that it is challenging to achieve a selectivity between these two closely related kinases. Two amino acid substitutions are present in the hinge region, where D127 in AAK1 is replaced by E131 in BIKE and F127 (AAK1) is replaced by Y132 (BIKE). While these minor changes at the hinge region cannot explain the preferential affinity of most inhibitors for AAK1, the third divergent residue may provide a clue. Despite the atypical consensus sequence of the glycine rich loop, a double glycine motif ensures the high flexibility of the loop (**Figure 5A**). In the AAK1 glycine rich loop, A58 is replaced by S63 in BIKE (**Figure 5B**). The bulkier polar side chain possibly clashes with certain hydrophobic substitution patterns on the benzylamides. Since the glycine rich loop residues of GAK have the same composition as those of AAK1, the benzylamide substitution patterns cannot confer comparable selectivity for AAK1 over GAK. However, structural analysis provided a possible explanation of how selectivity between the closely related kinases AAK1 and BIKE can be achieved.

**Figure 5:**
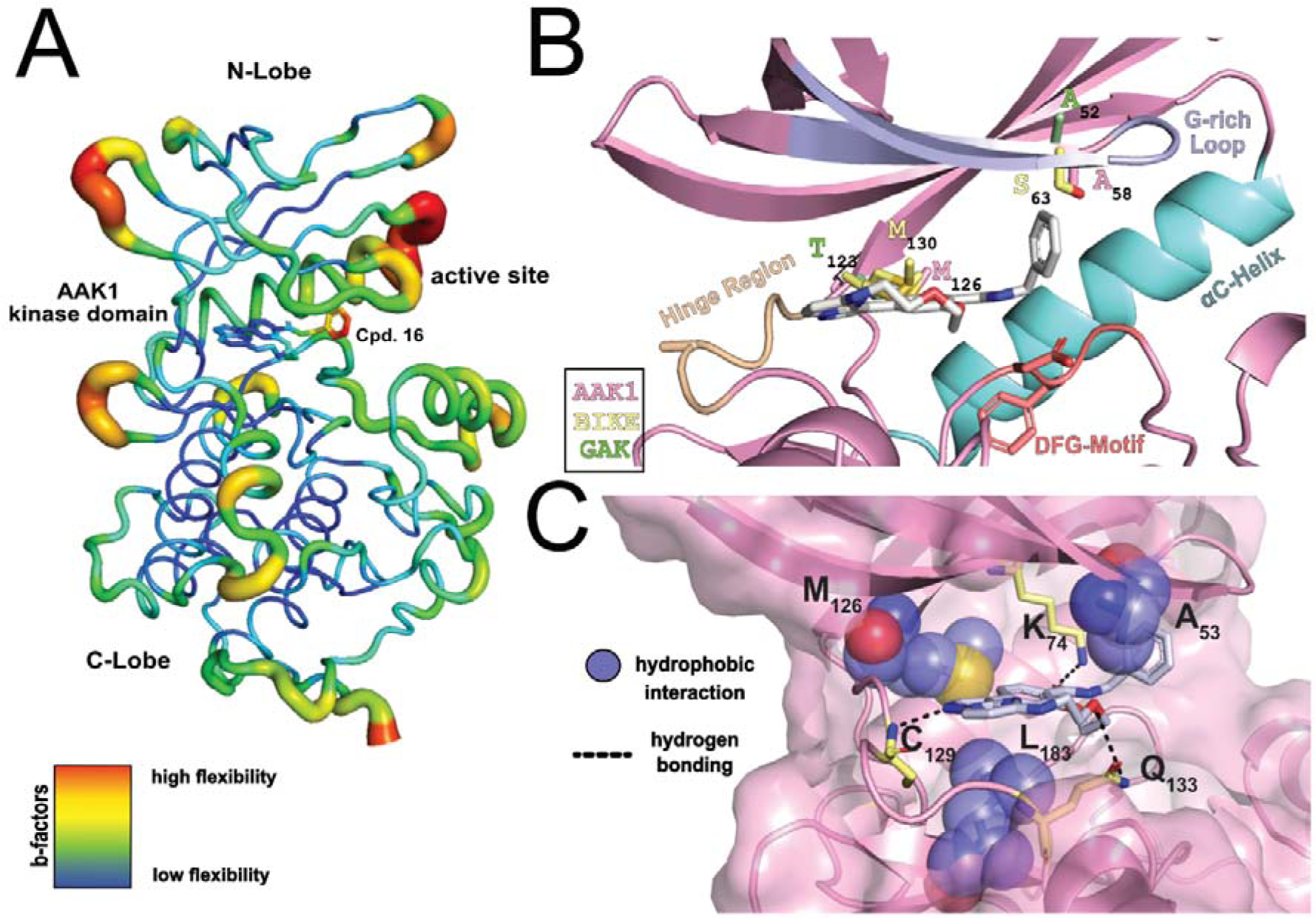
Crystal structure of AAK1 with Compound 18. **A:** Co-crystal structure of AAK1 with the macrocyclic inhibitor compound **18** (AAK1, PDB: 9QB5). Highlighting the flexible structures within the kinase through a b-factor colour code. **B:** Active site of AAK1 bound to compound **18**. Structural motifs are accentuated in different colours. Distant residues in AAK1 (pink), GAK (green) and BIKE (yellow), emphasize the highly conserved active site within the NAK kinases. **C:** Binding mode of compound **18**. Spheric displayed amino acids emphasize hydrophobic interactions, while hydrogen bonding’s are depicted as dotted lines between 18 and the respective amino acids (yellow sticks).

### Macrocyclic pyrazolo[1,5-a]pyrimidine inhibits AP-2 complex phosphorylation and possess antiviral activity

It is known that NAK kinases mediate AP-2 complex activation (**Figure 6A**) by phosphorylation at T156. We therefore investigated the ability of certain macrocycles to inhibit AP-2 complex phosphorylation. (**Figure 6B-C**).(17, 12, 54, 14, 11) We selected compounds **18** and **LP-935509** as compounds targeting all AP2-associated kinases (AAK1, BIKE, GAK) and compound **16** as the most potent macrocycle with selectivity for AAK1 over BIKE. hTERT RPE-1 cells were selected as a non-cancerous cell line, as all three kinases are equally expressed in these cells. To determine the influence of the compounds on the AP-2-complex, hTERT RPE-1cells were incubated with **16**, **18** and **LP-935509** at various concentrations for 2 hours. **LP-935509** was used as a positive control compound at a high concentration, while the macrocyclic compounds **16** and **18** were tested up to 2 µM to distinguish between effects driven by AAK1 and/or BIKE. The analysis was performed by Western Blot and showed inhibition of phosphorylation for all compounds at their respective concentrations (**Figure 6C**). While incubation with **LP-935509** at 10 µM resulted in an inhibition of approximately 60%, compound **18** achieved a comparable inhibition at the lower concentration of 2 µM. Compound **16** caused a slightly lower inhibition of 40% at the same concentration (2 µM).

**Figure 6:**
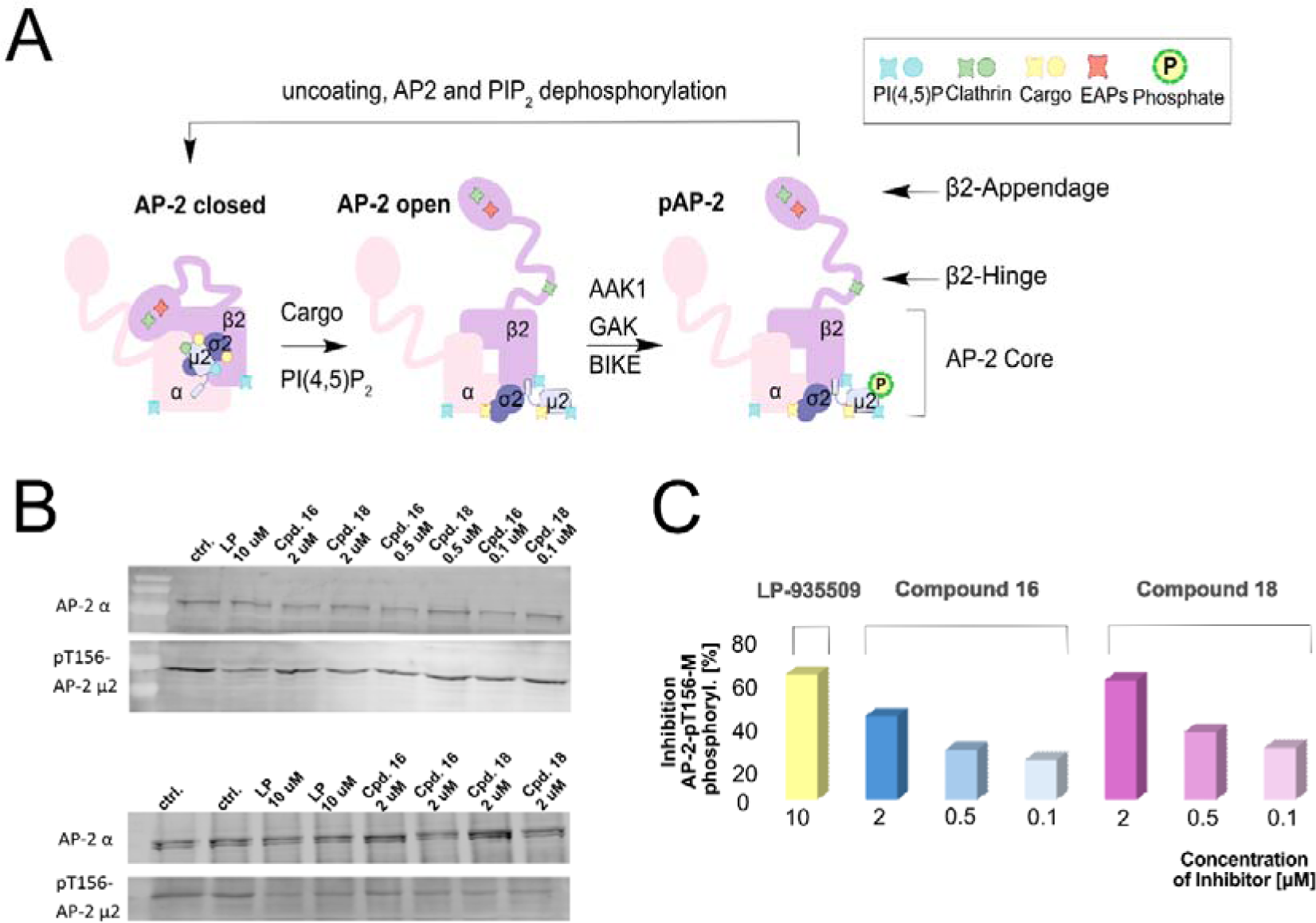
Regulation of the AP2-complex through phosphorylation and inhibition of said complex with small molecule kinase inhibitors (SMKIs) **A:** AP2 conformational changes and phosphorylation regulate AP2-complex interaction (adapted from Redlingshöfer et al.). Inactive lipid and protein binding sites, as well as a covered hinge-clathrin binding site are key features of the ‘closed’ AP2 structure.(55) Its recruitment to the plasma membrane is followed by Phospholipid (PI(4,5)P_2_) binding, which enables cargo binding. Thereupon, the ‘open’ AP2 conformation is triggered, now able to bind clathrin. Phosphorylation of the µ2-subunit occurs by kinases like AAK1, BIKE and GAK. Thereby, a new binding site is formed, which serves as an interaction site for EAPs (appendage-endocytic accessory proteins), whose binding enable CCV maturation. This cycle is restored by CCV uncoating, dephosphorylation of phospholipids and AP2 and dissociation of AP2 from the membrane to the cytosol.(56) **B:** Western Blot analysis of AP2-µ2-subunit phosphorylation at T156 after 2 hour incubation of hTERT RPE-1 cells with **16, 18** and **LP-935509** (**LP**) at various concentrations. Compounds **16** and **18** were used at concentrations of 2, 0.5 and 0.1 µM, while the concentration was 10 µM for **LP-935509**. **C:** Percentage Inhibition of AP2-µ2-subunit phosphorylation at T156 by **16, 18** and **LP-935509** at various concentrations. The percentage inhibition is depicted in bars, coloured according to each compound (**LP-935509**: yellow, **16**: blue, **18**: pink) and dependent on the concentration in different colour shades (lower concentration in lighter shade).

Encouraged by the results that the selective inhibition of AAK1 and GAK, without affecting BIKE, already led to an inhibition of the phosphorylation of the AP-2 complex, we investigated selected compounds from this series for their antiviral activity (**Figure 7**). We chose compounds **16** and **27**, which showed a potent binding for AAK1 and selectivity over BIKE. **SGC-GAK-1** was used as a positive control for selective GAK inhibition against SARS-CoV-2.(27) The Huh7 (DENV-2), U-87 MG (VEEV(-TC83)) and Vero E6/TMPRSS2 (SARS-CoV-2, WA1) cells were pre-treated with compounds for 1h, followed by infection with DENV-2 (48 h incubation), VEEV(-TC83) (18-20 h incubation) and SARS-CoV-2 (24 h incubation), respectively. Viral infection was measured by luciferase assays (infection with wild-type virus, carrying a Renilla or Nano luciferase reporter genes), while cell viability was measured using AlamarBlue assays. Compared to the potent GAK inhibitor **SGC**-**GAK**-**1**, which had an EC_50_ value of 0.75 µM against SARS-CoV-2 in Vero E6/TMPRSS2 cells and other cells(27), compounds **16** and **27** showed no significant antiviral activity against SARS-CoV-2. In the DENV-2 infection assay with Huh7 liver cancer cells, compound **27** showed weak antiviral activity with an EC_50_ of 3.5 µM, while compound **16** showed no significant antiviral activity up to a concentration of 10 µM. No activity was also observed for compound **16** in the VEEV(TC-83) infection assay, while compound **27** showed weak antiviral activity against the VEEV(TC-83) infection of U-87 MG cells, with an EC_50_ of 4.8 µM (**Figure 7**). Despite the potency of compound **16** for AAK1 (40 nM) and GAK (80 nM), it showed no significant antiviral activity, compared to compound **27**, which was a less active on both kinases (AAK1: 150 nM, GAK: 230 nM). However, while both compounds inhibited AAK1 and GAK, they lacked BIKE activity. Compound **27** showed only a 13.5-fold selectivity for AAK1, while compound **16** had an 85-fold selectivity for AAK1 and thus showed a slightly higher EC_50_ value in the NanoBRET assay on BIKE (see Table 1). These results were consistent with the work of Pu et al. who showed that DENV-2 infection is also regulated by BIKE.(14) Thus, the antiviral activity of **27** against infection of U-87 MG cells with VEEV(TC-83) may also be due to synergistic effects of pan NAK inhibition.

**Figure 7:**
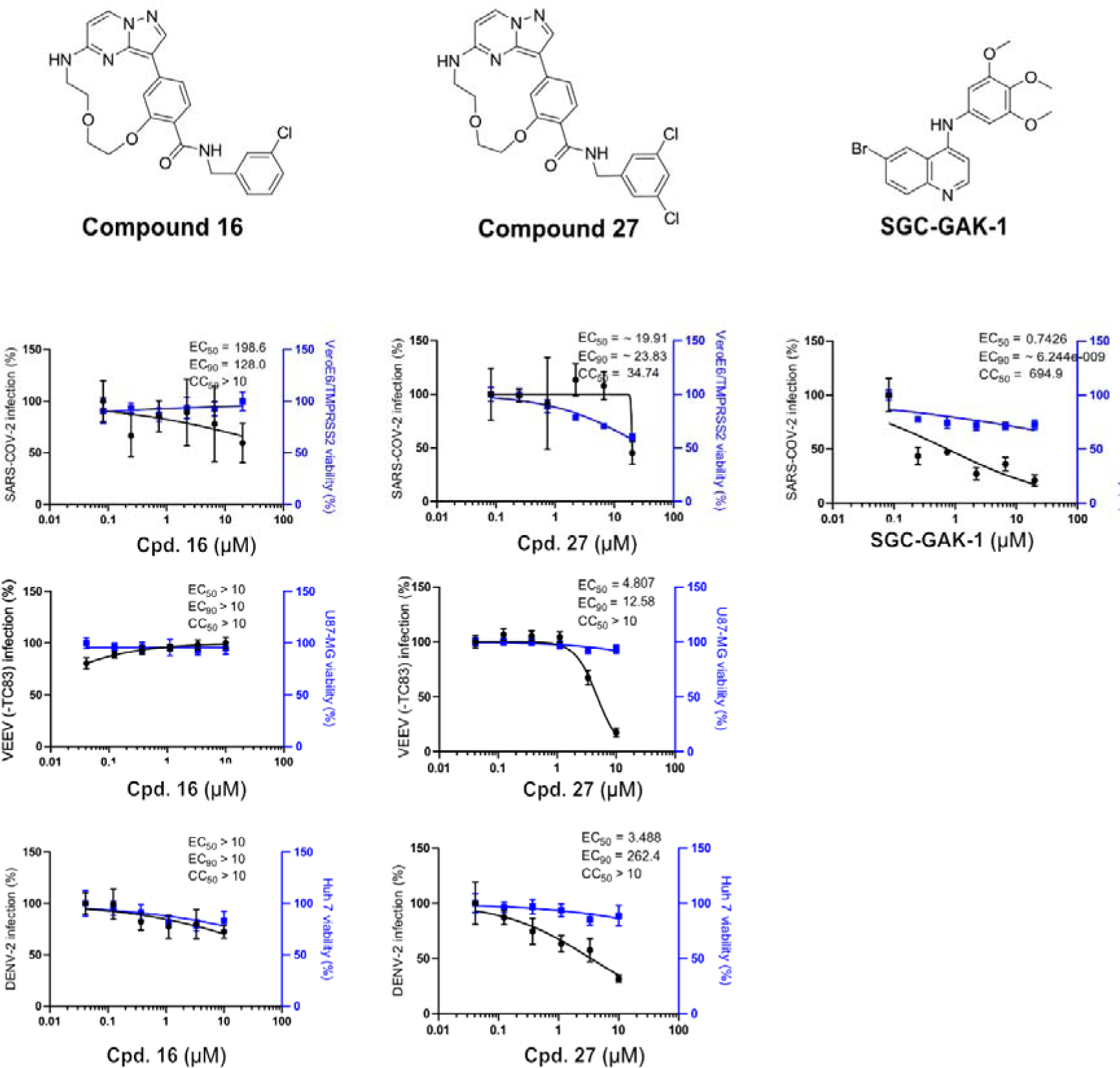
Evaluation of viral infection after incubation with **Compounds 16** and **27** in vitro. Antiviral activity of **16** and **27** was tested in a SARS-CoV-2, VEEV(-TC83) and DENV**-**2 infection assay. SGC-GAK-1 was used as positive control in the SARS-CoV-2 infection assay. The antiviral activity against SARS-CoV-2 was tested with Vero E6/TMPRSS2. For the VEEV(-TC83) infection, U87-MG cells were used and the antiviral activity in the DENV-2 infection assay was tested with Huh7 liver cancer cells. The compounds structures, their respective EC50- and CC50-values are displayed.

## Conclusion

The AAK1 inhibitors known to date are generally dual AAK1/BIKE inhibitors, as BIKE is the closest related kinase to AAK1. The aim of this study was to develop an inhibitor based on the pyrazolo[1,5-a]pyrimidine scaffold with an improved selectivity profile between the two closely related kinases AAK1 and BIKE. To this end, we started from the literature inhibitor **LP-935509** and combined it with our macrocyclic pyrazolo[1,5-a]pyrimidine inhibitors, previously described in the literature, to restrict the free rotatability within the ATP binding pocket. The focus of this SAR was mainly on the back-pocket, which we optimized to represent multiple compounds with different activity profiles within the NAK family. The most promising compound was **16**, which proved to be the most potent AAK1 inhibitor (40 nM) with the best selectivity window to BIKE (85-fold selectivity of AAK1 compared to BIKE), but still with slightly lower activity on GAK (80 nM). In addition, we have identified several compounds with pan-NAK activity and evaluated a number of compounds for their ability to reduce phosphorylation of the AP-2 complex. We observed a greater reduction in the phosphorylation of the AP-2 complex for our pan-NAK inhibitor **18** than for the dual AAK1/GAK inhibitor **16**. Also, in terms of its antiviral activity against 3 different RNA viruses, the antiviral potency of compound **16** was low, whereas a slightly better activity was seen with the closely related derivative **27**, which was slightly more potent at BIKE. We therefore assume that for antiviral activity, pan NAK activity is probably more efficacious, at least for the viruses investigated. Nevertheless, we were able to determine the binding mode of the macrocyclic pyrazolo[1,5-a]pyrimidines to AAK1 by cocrystal structure analysis and also provide a rationale how to achieve selectivity between the two closely related kinases. While the improved isoform selectivity did not improve anti viral activity, or study provided insights on the target profiles that need to be inhibited within the NAK family to prevent viral entry.

## Experimental Section Materials

### Chemistry

Detailed information on the synthesis of compounds and the analytical data for the final compounds can be found in the Supporting Information. All commercial chemicals were purchased from common suppliers with a purity ≥ 95% and were used without further purification. The solvents with an analytical grade were obtained from VWR Chemicals and Merck and all dry solvents from Acros Organics. All reactions were proceeded under an argon atmosphere. The thin layer chromatography was done with silica gel on aluminum foils (60 Å pore diameter) obtained from Macherey-Nagel and visualized with ultraviolet light (λ = 254 and 365 nm). The purification of the compounds was done by flash chromatography. A puriFlash XS 420 device with a UV-VIS multiwave detector (200−400 nm) from Interchim was used with pre-packed normal-phase PF-SIHP silica columns with particle sizes of 15 and 30 μm (Interchim). Preparative purification by HPLC was carried out on an Agilent 1260 Infinity II device using an Eclipse XDB-C18 (Agilent, 21.2 x 250mm, 7µm) reversed phase column. A suitable gradient (flow rate 21 ml/min.) was used, with 0.1% TFA in water (A) and 0.1% TFA S25 in acetonitrile (B), as a mobile phase. The nuclear magnetic resonance spectroscopy (NMR) was performed with DPX250, AV300, AV400 or AV500 MHz spectrometers from Bruker. Chemical shifts (δ) are reported in parts per million (ppm). DMSO-d6, chloroform-d and methylene chloride-d2 was used as a solvent, and the spectra were calibrated to the solvent signal: 2.50 ppm (1H NMR) or 39.52 ppm (13C NMR) for DMSO-d6, 7.26 ppm (1H NMR) or 77.16 ppm (13C NMR) for chloroform-d and 5.32 ppm (1H NMR) or 54.00 ppm (13C NMR) for methylene chloride-d2. Coupling constants (J) were reported in hertz (Hz) and multiplicities were designated as followed: s (singlet), d (doublet), dd (doublet of doublet), t (triplet), dt (doublet of triplets), td (triplet of doublets), ddd (doublet of doublet of doublet), q (quartet), m (multiplet). Mass spectra were measured on a Surveyor MSQ device from ThermoFisher measuring in the positive- or negative-ion mode. Final compounds were additionally characterized by HRMS using a MALDI LTQ Orbitrap XL from ThermoScientific. The purity of the final compounds was determined by HPLC using an Agilent 1260 Infinity II device with a 1260 DAD HS detector (G7117C; 254 nm, 280 nm, 310 nm) and a LC/MSD device (G6125B, ESI pos. 100-1000). The compounds were analyzed on a Poroshell 120 EC-C18 (Agilent, 3 x 150 mm, 2.7 µm) reversed phase column using 0.1% formic acid in water (A) and 0.1% formic acid in acetonitrile (B) as a mobile phase. The following gradient was used: 0 min 5% B - 2 min 5% B - 8 min 98% B - 10 min 98% B (flow rate of 0.5 mL/min). UV-detection was performed at 254, 280 and 310 nm and all compounds used for further biological characterizations showed a purity ≥95%. No unexpected or unusually high safety hazards were encountered.

Error! Reference source not found.:

Recombinant protein kinase domains with a concentration of 2 μM were mixed with a 10 μM compound solution in DMSO, using a final buffer consisting of 20 mM HEPES, pHLJ7.5, and 500 mM NaCl. SYPRO Orange (5000×, Invitrogen) was added as a fluorescence probe (1 µl per mL) in a final concentration of 5x. Subsequently, temperature-dependent protein unfolding profiles were measured, using the QuantStudio™ 5 realtime PCR machine (Thermo Fisher). Excitation and emission filters were set to 465 nm and 590 nm. The temperature was raised with a step rate of 3°C per minute. Data points were analysed with the internal software (Thermal Shift SoftwareTM Version 1.4, Thermo Fisher) using the Boltzmann equation to determine the inflection point of the transition curve. Differences in melting temperature are given as ΔTm values in °C. Measurements were performed in triplicates.

Error! Reference source not found.

The NanoBRET Assay was conducted following the methodology described previously by Schwalm et al.(57) HEK293T cells (Human embryonic kidney 293T cells) were transfected in separate experiments with one of the following plasmid vectors: AAK1-NanoLuc Fusion Vector (Promega, NV1001), BMP2K-NanoLuc Fusion Vector (Promega, NV1091), GAK-NanoLuc Fusion Vector (Promega, NV1421), or STK16-NanoLuc Fusion Vector (Promega, NV2091). Each vector was used to express a specific kinase fused with an N-terminal NanoLuc Luciferase: full-length AAK1 kinase, full-length BMP2K kinase, full-length GAK kinase, or full-length STK16 kinase, respectively. This approach ensured that each population of transfected cells expressed one type of kinase-NanoLuc fusion protein. After trypsinization and resuspension in Opti-MEM (Life Technologies), the cells were seeded into white 384-well plates (Greiner 784075) at a density of 2 × 10^5 cells/mL. They were then transfected using FuGENE HD (Promega, E2312). The proteins were allowed to express for 20 h at 37 °C/5% CO2. Using an ECHO acoustic dispenser (Labcyte), the transfected cells were exposed to increasing compound concentrations and a cell-permeable fluorescent NanoBRET Tracer. A NanoBRET Tracer, that binds the specific target protein was chosen for each experiment. Tracer K10 (Promega, N2642) was used for the cells overexpressing AAK1 kinase or STK16 kinase, Tracer K5 (Promega, N2482) was used for the cells overexpressing BMP2K kinase or GAK kinase. The cells were treated with a fixed concentration of the NanoBRET Tracers corresponding to the Tracer KD app value, as given in the TracerDB (tracerdb.org). To establish equilibrium, the system was incubated for 2 h at 37 °C/5% CO2. In accordance with the manufacturer’s protocol, NanoBRET NanoGlo Substrate (Promega, N1573) was added, and filtered luminescence was measured using the PHERAstar plate reader (BMG Labtech) equipped with a luminescence filter pair (450 nm BP filter for the donor and 610 nm LP filter for the acceptor). Subsequently, competitive displacement data were analyzed and graphed using GraphPad Prism 10, employing a normalized 3-parameter curve fit with the following equation: Y = 100/(1 + 10^(X – log IC^_50_)). For the generation of the permeabilized data, the transfected cells were permeabilized using digitonin (0.05 µg/µL). After an equilibration period of 10 min, luminescence measurements were taken, and the displacement data were analyzed as described previously.

Error! Reference source not found.

### Protein Expression and Purification

For protein expression, the plasmid containing DNA encoding AAK1KD (residues T27-A365) cloned into the kanamycin resistant vector pNIC-CTH0 was transformed into competent Rosetta (DE3) E. coli cells (Novagen). Cultures were grown in Terrific broth medium (4×1L) at 37°C. Protein expression was induced using IPTG and the cultures incubated overnight at 18°C. The cells were harvested by centrifugation (6000 rpm, 15 min, 4°C) and stored at −20°C.

For the purification of AAK1KD, the cell pellets were resuspended in lysis buffer (50 mM HEPES pH 7.4, 500 mM NaCl, 20 mM imidazole, 0.5 mM TCEP, 5% glycerol) and lysis was performed by sonication. The lysate was cleared by centrifugation (23000 rpm, 30 min, 4°C) and the supernatant then loaded on a Ni-NTA column. After elution of the flowthrough, the resin loaded with the His-tagged protein was washed with lysis buffer (150 mL) prior to elution with lysis buffer containing 300 mM imidazole. Cleavage of the His-tag was performed by addition of TEV protease and incubation overnight in dialysis buffer (50 mM HEPES pH 7.4, 500 mM NaCl, 0.5 mM TCEP, 5% glycerol). The cleaved His-tag was removed by IMAC using Ni-NTA resin and the protein containing flowthrough concentrated prior to purification by SEC using gel filtration buffer (20 mM HEPES pH 7.4, 150 mM NaCl, 0.5 mM TCEP, 5% glycerol) and a HiLoad 16/600 Superdex 200 pg gel filtration column connected to an AKTA Xpress system. Protein containing fractions were pooled, concentrated to 2-3 mg/ml and flash-frozen in liquid nitrogen before storage at −80°C.

### Crystallization

For the cocrystallization of AAK1KD with compound 18, the ligand was added to the protein (2 3 mg/ml) at 500 mM final ligand concentration. After incubation on ice for 10 min and centrifugation (13000 rpm, 10 min, 4°C) the sample was concentrated to a final protein concentration of 16 mg/ml. Crystallization drops were set up using a mosquito® liquid handler (TTP Labtech) in ratios of 2:1, 1:1 and 1:2 of protein to precipitant solution. Crystals of AAK1KD with compound 18 grew at 20°C within several days in a crystallization condition containing 0.2 M ammonium sulfate and 30% PEG4000. Crystals were cryoprotected in precipitant solution containing 25% ethylene glycol.

### Data collection and structure determination

Diffraction data was collected at Swiss Light Source beamline X06SA (Villigen, CH) at a wavelength of 1.00002 Å and processed with the SLS automated data processing(58) pipline. The structure was solved by molecular replacement with Phaser (McCoy et al., 2005 (59)) using AAK1 (PDB ID: 4WSQ) as a model. Subsequent rounds of manual structure building and refinement were performed using Coot (Emsley and Cowtan, 2004 (60)) and Refmac5 (Murshudov et al., 1997(61)). Final validation was performed using MolProbity (Chen et al., 2010 (62)). The model of AAK1KD with Compound 18 has been deposited to the PDB with the PDB ID 9QB5.

Error! Reference source not found.

Compressed CETSA MS assays, also known as PISA (Proteome Integral Stability Alteration assay), were performed as previously described by Chernobrovkin et. al., with some minor changes.(63) K562 cells were obtained from ATCC. K562 cells were mixed with an equal volume of compound solution, final concentration of 30 M, in a buffer (20 mMHEPES, 138 mM NaCl, 5 mM KCl, 2 mM CaCl2, 1 mM MgCl2, pH 7.4). One percent and DMSO was used as a control. Cells were incubated at +37 C for 60 min. The cell suspension was split into 12 aliquots which were heated at a different temperature between 44 and 66 C for 3 min. After the heating, the cells were lysed through freeze–thawing them 3 times and the lysed aliquots were pooled together. Cell debris and aggregates were removed through centrifugation (20 min at 30.000 g).

Error! Reference source not found.

Compounds were dissolved in DMSO at 10 mM stock solution and stored at −20°C. hTERT RPE-1 cells (ATCC) were incubated in the presence of inhibitors at indicated concentration or the equivalent concentration of DMSO for 2 hours in complete medium at 37°C. To compare the levels of T156 phosphorylation of the AP-2 μ2 subunit across inhibitor-treated samples, ratiometric analysis of phospho-T156 of AP-2 μ2 and AP-2 α subunit levels, obtained by western blotting, was employed. For this, the AP2 complex was immunoprecipitated using the AP.6 antibody. Cells were washed three times with PBS, lysed in 0.1% NP40 buffer, sonicated, and lysates were clarified by centrifugation. After incubation with supernatant, protein A/G Sepharose beads were washed three times with lysis buffer and boiled with Laemmli sample buffer before SDS-PAGE gel loading. Nitrocellulose membranes were used for transfer and then probed with antibodies D4F3 (Cell Signalling) to detect and quantify p-T156 AP-2 μ2, while the AP-2 α subunit was detected using AC1-M11 (Abcam). Data are presented as a mean across three independent biological replicates.

Error! Reference source not found.

### Cell lines for DENV-2, VEEV (TC-83), and SARS-CoV-2 infection assays

Human hepatoma (Huh7, Apath, L.L.C, New York, NY, USA), U-87 MG (ATCC), BHK-21 (ATCC), Vero E6/TMPRSS2 (JCRB cell bank, #cat JCRB1819) cells were grown in Dulbecco’s Modified Eagle’s medium DMEM (Gibco) supplemented with 10% FCS (Biowest), 1% nonessential amino acids (Corning), 1% L-glutamine (Gibco), and 1% penicillin-streptomycin (Gibco). Vero E6 (ATCC) cells were maintained in DMEM supplemented with 10% FCS, 1% L-glutamine, 1% Pen-strep, 1% NEAA, 1% HEPES (Gibco), 1% Sodium pyruvate (Thermo Fisher Scientific). cells were grown in All the cells were maintained in a humidified incubator with 5% CO_2_ at 37 °C and tested negative for mycoplasma by MycoAlert (Lonza, Morristown, NJ). Cells from passages 14–15 (P14–15) were used for this study.

### Virus constructs

DENV-2 (New Guinea C strain) TSV01 Renilla reporter plasmid (pACYC NGC FL) used for the production of (DENV2-Rluc) was a gift from Pei-Yong Shi (The University of Texas Medical Branch).(64) The plasmid encoding infectious VEEV (TC-83) with a nanoluciferase reporter (VEEV-TC-83-Cap-nLuc-Tav) used for virus production (VEEV-TC-83-nLuc) was a gift from Dr. William B. Klimstra (Department of Immunology, University of Pittsburgh, Pittsburgh).(65) Plasmids used to produce SARS-CoV-2 (SARS-CoV-2 expressing Nluc-reporter gene) were a gift from Dr. Luis Martinez-Sobrido.(66)

### Virus production

DENV2-Rluc RNA was transcribed in vitro using mMessage/mMachine (Ambion) kits. DENV2-Rluc was produced by electroporating RNA into BHK-21 cells, supernatants were harvested on day 10 post-electroporation. The VEEV-TC-83-nLuc virus was harvested from the supernatant 24 h post-electroporation. Viral stocks for SARS-CoV-2-nluc were generated as previously described. Viruses produced in Vero E6/TMPRSS2 cells and passaged 3–4 times were used for the experiments.(67) Supernatants were collected, clarified, and stored at −80 °C, and viral titers were determined via plaque assays on BHK-21 (DENV, VEEV) or Vero E6 cells (SARS-CoV-2).

### Infection assays and drug screening

Huh7, U-87 MG and Vero e6/TMPRSS2 cells were pretreated with the compounds or DMSO for 1 h prior to infection with DENV2-Rluc, VEEV-TC-83-nLuc, and SARS-CoV-2-Nluc at a multiplicity of infection (MOI) of 0.01 (DENV-2, VEEV) and MOI of 0.5 (SARS-CoV-2) (n = 4). The inhibitors were maintained for the duration of the experiment. Overall viral infection was measured at 45-48 h post-infection using a Renilla luciferase substrate via luciferase assays (DENV2) and 18 h post-infection (VEEV (TC-83)) and 24 h post-infection (SARS-CoV-2) using a nanoluciferase assay. The relative light units (RLUs) were normalized to DMSO-treated cells (set as 100%).

### Viability assays

Cell viability was assessed using alamarBlue reagent (Invitrogen) according to the manufacturer’s protocol. Fluorescence was detected at 560 nm on GloMax Discover Microplate Reader (Promega).

### Statistical analysis

All data were analyzed with GraphPad Prism software. Half-maximal effective concentrations (EC_50_) and half-maximal cytotoxic concentrations (CC_50_) were measured by fitting of data to a 3-parameter logistic curve.

### Study Approval

SARS-CoV-2 work was conducted in biosafety level 3 (BSL3) facility at Stanford University, according to CDC and institutional guidelines.

## Associated content

The Supporting Information is available free of charge at #

It includes:

- Synthetic procedures, DSF kinase selectivity assay within the NAK family, EC_50_-values against STK16 (NanoBRET), HPLC and NMR spectra (PDF)
- In-house DSF kinase selectivity data (XLSX)
- KINOMEscan^®^ selectivity data from Eurofins (XLSX)
- CETSA shifts from PELAGO BIOSCIENCE (XLSX)
- EC_50_ values for the complete compound set determined via nanoBRET

## Author Contributions

T.E.M., S.K. and T.H. designed the project; T.E.M., C.G.K., J.A.A. synthesized the compounds; T.A.L.E. performed NanoBRET assay; S.K., T.H., S.E. and S.M. supervised research. T.E.M., C.G.K., J.A.A. performed DSF measurements; D.T., M.K., and S.E. performed antiviral and cell viability assays; F.P. and S.M. co-crystallized and refined the structure of AAK1, T.T. and D.M.-M. performed the CETSA experiments; Z.K. performed the Western Blot experiments; the manuscript was written by T.E.M., S.K. and T.H. with contributions from all coauthors.

## Acknowledgements

The authors are grateful for support by the Structural Genomics Consortium (SGC), a registered charity (no: 1097737) that receives funds from Bayer AG, Boehringer Ingelheim, Bristol Myers Squibb, Genentech, Genome Canada through Ontario Genomics Institute [OGI-196], EU/EFPIA/OICR/McGill/KTH/Diamond Innovative Medicines Initiative 2 Joint Undertaking [EUbOPEN grant 875510], Janssen, Pfizer and Takeda. S.K. would like to acknowledge funding from the Frankfurt Cancer Institute (FCI), an institute supported by LOEWE. T.H. and S.K. are grateful for support by the ENABLE project “Unraveling mechanisms driving cellular homeostasis, inflammation and infection to enable new approaches in translational medicine”. SK and SM are also funded by the German Cancer Aid translational grant TACTIC. ZK would like to acknowledge funding from the Wellcome Trust Fellowship 220597/Z/20/Z.

## Abbreviations

AAK1: adaptor associated kinase 1
ACN: acetonitrile
ALK: anaplastic lymphoma kinase
AP2: adaptor protein 2
ATP: Adenosine Triphosphate
BIKE: BMP-2-inducible kinase
BOC: tert-Butyloxycarbonyl
CCV: Clathrin-coated vesicles
CME: Clathrin mediated endocytosis
DAAs: Direct-acting antivirals
DCM: Dichloromethane
DENV: Dengue virus
DIAD: Diisopropyl azodicarboxylate
DMF: Dimethylformamide
GAK: cyclin G associated kinase
HCV: Hepatitis C virus
MeOH: Methanol
MPSK1: myristoylated and palmitoylated serine/threonine kinase 1
NAK: Numb associated kinases
NBS: N-Bromosuccinimide
on.: Over night
PyAOP: (7-Azabenzotriazol-1-yloxy)tripyrrolidinophosphonium hexafluorophosphate
PPh_3_: Triphenylphosphine
RIPK2: Receptor-interacting serine/threonine-protein kinase 2
RNA: Ribonucleic acid
RT: Room temperature
STK17A: Serine/ Threonine kinase 17A
TBAF: Tertabutylammoniume fluoride
TBDMS: Tertbutyldimethylsilyl
TBDMSCl: Tertbutyldimethylsilylchloride
TEA: Triethylamine
THF: Tetrahydrofurane
VEEV: Venezuelan Equine Encephalitis virus

## Footnotes

1 ΔT_m_-values were determined as average of all measurements performed.

2 Staurosporine was used as Reference Compound to determine the respective percentage Shifts for each kinase

## Notes

### Competing Interest Statement

The authors have declared no competing interest.

## References

1. Ahmad, T.; Haroon; Baig, M.; Hui, J. Coronavirus Disease 2019 (COVID-19) Pandemic and Economic Impact. Pakistan Journal of Medical Sciences 2020, 36 (COVID19-S4), S73–S78. DOI: 10.12669/pjms.36.COVID19-S4.2638.

2. Bekerman, E.; Neveu, G.; Shulla, A.; Brannan, J.; Pu, S.-Y.; Wang, S.; Xiao, F.; Barouch-Bentov, R.; Bakken, R. R.; Mateo, R.; Govero, J.; Nagamine, C. M.; Diamond, M. S.; Jonghe, S. de; Herdewijn, P.; Dye, J. M.; Randall, G.; Einav, S. Anticancer kinase inhibitors impair intracellular viral trafficking and exert broad-spectrum antiviral effects. The Journal of clinical investigation 2017, 127 (4), 1338–1352. DOI: 10.1172/JCI89857.

3. Bekerman, E.; Einav, S. Infectious disease. Combating emerging viral threats. Science (New York, N.Y.) 2015, 348 (6232), 282–283. DOI: 10.1126/science.aaa3778.

4. Grove, J.; Marsh, M. The cell biology of receptor-mediated virus entry. The Journal of cell biology 2011, 195 (7), 1071–1082. DOI: 10.1083/jcb.201108131.

5. Robinson, M.; Schor, S.; Barouch-Bentov, R.; Einav, S. Viral journeys on the intracellular highways. Cellular and Molecular Life Sciences 2018, 75 (20), 3693–3714. DOI: 10.1007/s00018-018-2882-0.

6. Strazic Geljic, I.; Kucan Brlic, P.; Musak, L.; Karner, D.; Ambriović-Ristov, A.; Jonjic, S.; Schu, P.; Rovis, T. L. Viral Interactions with Adaptor-Protein Complexes: A Ubiquitous Trait among Viral Species. International Journal of Molecular Sciences 2021, 22 (10), 5274. DOI: 10.3390/ijms22105274.

7. Helenius, A.; Kartenbeck, J.; Simons, K.; Fries, E. On the entry of Semliki forest virus into BHK-21 cells. J Cell Biol 1980, 84 (2), 404–420. DOI: 10.1083/jcb.84.2.404.

8. Briant, K.; Redlingshöfer, L.; Brodsky, F. M. Clathrin’s life beyond 40: Connecting biochemistry with physiology and disease. Current Opinion in Cell Biology 2020, 65, 141–149. DOI: 10.1016/j.ceb.2020.06.004.

9. Owen, D. J.; Collins, B. M.; Evans, P. R. Adaptors for clathrin coats: structure and function. Annual Review of Cell and Developmental Biology 2004, 20 (Volume 20, 2004), 153–191. DOI: 10.1146/annurev.cellbio.20.010403.104543.

10. Ramesh, S. T.; Navyasree, K. V.; Sah, S.; Ashok, A. B.; Qathoon, N.; Mohanty, S.; Swain, R. K.; Umasankar, P. K. BMP2K phosphorylates AP-2 and regulates clathrin-mediated endocytosis. Traffic 2021, 22 (11), 377–396. DOI: 10.1111/tra.12814.

11. Olusanya, O.; Andrews, P. D.; Swedlow, J. R.; Smythe, E. Phosphorylation of threonine 156 of the mu2 subunit of the AP2 complex is essential for endocytosis in vitro and in vivo. Current biology : CB 2001, 11 (11), 896–900. DOI: 10.1016/s0960-9822(01)00240-8.

12. Conner, S. D.; Schmid, S. L. Identification of an adaptor-associated kinase, AAK1, as a regulator of clathrin-mediated endocytosis. J Cell Biol 2002, 156 (5), 921–929. DOI: 10.1083/jcb.200108123.

13. Umeda, A.; Meyerholz, A.; Ungewickell, E. Identification of the universal cofactor (auxilin 2) in clathrin coat dissociation. European Journal of Cell Biology 2000, 79 (5), 336–342. DOI: 10.1078/S0171-9335(04)70037-0.

14. Pu, S.; Schor, S.; Karim, M.; Saul, S.; Robinson, M.; Kumar, S.; Prugar, L. I.; Dorosky, D. E.; Brannan, J.; Dye, J. M.; Einav, S. BIKE regulates dengue virus infection and is a cellular target for broad-spectrum antivirals. Antiviral Research 2020, 184, 104966. DOI: 10.1016/j.antiviral.2020.104966.

15. Wrobel, A. G.; Kadlecova, Z.; Kamenicky, J.; Yang, J.-C.; Herrmann, T.; Kelly, B. T.; McCoy, A. J.; Evans, P. R.; Martin, S.; Müller, S.; Salomon, S.; Sroubek, F.; Neuhaus, D.; Höning, S.; Owen, D. J. Temporal Ordering in Endocytic Clathrin-Coated Vesicle Formation via AP2 Phosphorylation. Developmental Cell 2019, 50 (4), 494–508.e11. DOI: 10.1016/j.devcel.2019.07.017.

16. Kirchhausen, T.; Owen, D.; Harrison, S. C. Molecular structure, function, and dynamics of clathrin-mediated membrane traffic. Cold Spring Harb Perspect Biol 2014, 6 (5), a016725. DOI: 10.1101/cshperspect.a016725.

17. Borner, G. H. H.; Antrobus, R.; Hirst, J.; Bhumbra, G. S.; Kozik, P.; Jackson, L. P.; Sahlender, D. A.; Robinson, M. S. Multivariate proteomic profiling identifies novel accessory proteins of coated vesicles. The Journal of cell biology 2012, 197 (1), 141–160. DOI: 10.1083/jcb.201111049.

18. Robinson, M.; Schor, S.; Barouch-Bentov, R.; Einav, S. Viral journeys on the intracellular highways. Cell. Mol. Life Sci. 2018, 75 (20), 3693–3714. DOI: 10.1007/s00018-018-2882-0.

19. Yamauchi, Y.; Helenius, A. Virus entry at a glance. Journal of cell science 2013, 126 (Pt 6), 1289–1295. DOI: 10.1242/jcs.119685.

20. Sorrell, F. J.; Szklarz, M.; Abdul Azeez, K. R.; Elkins, J. M.; Knapp, S. Family-wide Structural Analysis of Human Numb-Associated Protein Kinases. Structure (London, England : 1993) 2016, 24 (3), 401–411. DOI: 10.1016/j.str.2015.12.015.

21. Asquith, C. R. M.; Berger, B.-T.; Wan, J.; Bennett, J. M.; Capuzzi, S. J.; Crona, D. J.; Drewry, D. H.; East, M. P.; Elkins, J. M.; Fedorov, O.; Godoi, P. H.; Hunter, D. M.; Knapp, S.; Müller, S.; Torrice, C. D.; Wells, C. I.; Earp, H. S.; Willson, T. M.; Zuercher, W. J. SGC-GAK-1: A Chemical Probe for Cyclin G Associated Kinase (GAK). Journal of medicinal chemistry 2019, 62 (5), 2830–2836. DOI: 10.1021/acs.jmedchem.8b01213.

22. Hartz, R. A.; Ahuja, V. T.; Nara, S. J.; Kumar, C. M. V.; Manepalli, R. K. V. L. P.; Sarvasiddhi, S. K.; Honkhambe, S.; Patankar, V.; Dasgupta, B.; Rajamani, R.; Muckelbauer, J. K.; Camac, D. M.; Ghosh, K.; Pokross, M.; Kiefer, S. E.; Brown, J. M.; Hunihan, L.; Gulianello, M.; Lewis, M.; Lippy, J. S.; Surti, N.; Hamman, B. D.; Allen, J.; Kostich, W. A.; Bronson, J. J.; Macor, J. E.; Dzierba, C. D. Bicyclic Heterocyclic Replacement of an Aryl Amide Leading to Potent and Kinase-Selective Adaptor Protein 2-Associated Kinase 1 Inhibitors. Journal of medicinal chemistry 2022, 65 (5), 4121–4155. DOI: 10.1021/acs.jmedchem.1c01966.

23. Martinez-Gualda, B.; Schols, D.; Jonghe, S. de. A patent review of adaptor associated kinase 1 (AAK1) inhibitors (2013-present). Expert Opinion on Therapeutic Patents 2021, 31 (10), 911–936. DOI: 10.1080/13543776.2021.1928637.

24. Kearns, A. E.; Donohue, M. M.; Sanyal, B.; Demay, M. B. Cloning and characterization of a novel protein kinase that impairs osteoblast differentiation in vitro. The Journal of biological chemistry 2001, 276 (45), 42213–42218. DOI: 10.1074/jbc.M106163200.

25. Xiao, F.; Wang, S.; Barouch-Bentov, R.; Neveu, G.; Pu, S.; Beer, M.; Schor, S.; Kumar, S.; Nicolaescu, V.; Lindenbach, B. D.; Randall, G.; Einav, S. Interactions between the Hepatitis C Virus Nonstructural 2 Protein and Host Adaptor Proteins 1 and 4 Orchestrate Virus Release. mBio 2018, 9 (2). DOI: 10.1128/mBio.02233-17.

26. Neveu, G.; Ziv-Av, A.; Barouch-Bentov, R.; Berkerman, E.; Mulholland, J.; Einav, S. AP-2-associated protein kinase 1 and cyclin G-associated kinase regulate hepatitis C virus entry and are potential drug targets. Journal of virology 2015, 89 (8), 4387–4404. DOI: 10.1128/JVI.02705-14.

27. Karim, M.; Saul, S.; Ghita, L.; Sahoo, M. K.; Ye, C.; Bhalla, N.; Lo, C.-W.; Jin, J.; Park, J.-G.; Martinez-Gualda, B.; East, M. P.; Johnson, G. L.; Pinsky, B. A.; Martinez-Sobrido, L.; Asquith, C. R. M.; Narayanan, A.; Jonghe, S. de; Einav, S. Numb-associated kinases are required for SARS-CoV-2 infection and are cellular targets for antiviral strategies. Antiviral Research 2022, 204, 105367. DOI: 10.1016/j.antiviral.2022.105367.

28. Alconada, A.; Bauer, U.; Hoflack, B. A tyrosine-based motif and a casein kinase II phosphorylation site regulate the intracellular trafficking of the varicella-zoster virus glycoprotein I, a protein localized in the trans-Golgi network. The EMBO Journal 1996, 15 (22), 6096–6110. DOI: 10.1002/j.1460-2075.1996.tb00998.x.

29. Bhattacharyya, S.; Hope, T. J.; Young, J. A. T. Differential requirements for clathrin endocytic pathway components in cellular entry by Ebola and Marburg glycoprotein pseudovirions. Virology 2011, 419 (1), 1–9. DOI: 10.1016/j.virol.2011.07.018.

30. Dutta, D.; Chakraborty, S.; Bandyopadhyay, C.; Valiya Veettil, M.; Ansari, M. A.; Singh, V. V.; Chandran, B. EphrinA2 regulates clathrin mediated KSHV endocytosis in fibroblast cells by coordinating integrin-associated signaling and c-Cbl directed polyubiquitination. PLOS Pathogens 2013, 9 (7), e1003510. DOI: 10.1371/journal.ppat.1003510.

31. Huang, H.-C.; Chen, C.-C.; Chang, W.-C.; Tao, M.-H.; Huang, C. Entry of hepatitis B virus into immortalized human primary hepatocytes by clathrin-dependent endocytosis. Journal of virology 2012, 86 (17), 9443–9453. DOI: 10.1128/JVI.00873-12.

32. Humphries, A. C.; Dodding, M. P.; Barry, D. J.; Collinson, L. M.; Durkin, C. H.; Way, M. Clathrin potentiates vaccinia-induced actin polymerization to facilitate viral spread. Cell Host & Microbe 2012, 12 (3), 346–359. DOI: 10.1016/j.chom.2012.08.002.

33. Neveu, G.; Barouch-Bentov, R.; Ziv-Av, A.; Gerber, D.; Jacob, Y.; Einav, S. Identification and targeting of an interaction between a tyrosine motif within hepatitis C virus core protein and AP2M1 essential for viral assembly. PLOS Pathogens 2012, 8 (8), e1002845. DOI: 10.1371/journal.ppat.1002845.

34. Ohka, S.; Ohno, H.; Tohyama, K.; Nomoto, A. Basolateral sorting of human poliovirus receptor alpha involves an interaction with the mu1B subunit of the clathrin adaptor complex in polarized epithelial cells. Biochemical and biophysical research communications 2001, 287 (4), 941–948. DOI: 10.1006/bbrc.2001.5660.

35. Ohno, H.; Aguilar, R. C.; Fournier, M. C.; Hennecke, S.; Cosson, P.; Bonifacino, J. S. Interaction of endocytic signals from the HIV-1 envelope glycoprotein complex with members of the adaptor medium chain family. Virology 1997, 238 (2), 305–315. DOI: 10.1006/viro.1997.8839.

36. Agrawal, T.; Schu, P.; Medigeshi, G. R. Adaptor protein complexes-1 and 3 are involved at distinct stages of flavivirus life-cycle. Scientific reports 2013, 3, 1813. DOI: 10.1038/srep01813.

37. Pu, S.-Y.; Xiao, F.; Schor, S.; Bekerman, E.; Zanini, F.; Barouch-Bentov, R.; Nagamine, C. M.; Einav, S. Feasibility and biological rationale of repurposing sunitinib and erlotinib for dengue treatment. Antiviral Research 2018, 155, 67–75. DOI: 10.1016/j.antiviral.2018.05.001.

38. Liu, F.; Wang, J.; Yang, X.; Li, B.; Wu, H.; Qi, S.; Chen, C.; Liu, X.; Yu, K.; Wang, W.; Zhao, Z.; Wang, A.; Chen, Y.; Wang, L.; Gray, N. S.; Liu, J.; Zhang, X.; Liu, Q. Discovery of a Highly Selective STK16 Kinase Inhibitor. ACS chemical biology 2016, 11 (6), 1537–1543. DOI: 10.1021/acschembio.6b00250.

39. Chaikuad, A.; Keates, T.; Vincke, C.; Kaufholz, M.; Zenn, M.; Zimmermann, B.; Gutiérrez, C.; Zhang, R.-G.; Hatzos-Skintges, C.; Joachimiak, A.; Muyldermans, S.; Herberg, F. W.; Knapp, S.; Müller, S. Structure of cyclin G-associated kinase (GAK) trapped in different conformations using nanobodies. The Biochemical journal 2014, 459 (1), 59–69. DOI: 10.1042/BJ20131399.

40. Garcia Jimenez, D.; Poongavanam, V.; Kihlberg, J. Macrocycles in Drug Discovery─Learning from the Past for the Future. Journal of medicinal chemistry 2023, 66 (8), 5377–5396. DOI: 10.1021/acs.jmedchem.3c00134.

41. Amrhein, J. A.; Knapp, S.; Hanke, T. Synthetic Opportunities and Challenges for Macrocyclic Kinase Inhibitors. Journal of medicinal chemistry 2021, 64 (12), 7991–8009. DOI: 10.1021/acs.jmedchem.1c00217.

42. Fang, Z.; Song, Y.; Zhan, P.; Zhang, Q.; Liu, X. Conformational restriction: an effective tactic in ‘follow-on’-based drug discovery. Future medicinal chemistry 2014, 6 (8), 885–901. DOI: 10.4155/fmc.14.50.

43. Amrhein, J. A.; Berger, L. M.; Balourdas, D.-I.; Joerger, A. C.; Menge, A.; Krämer, A.; Frischkorn, J. M.; Berger, B.-T.; Elson, L.; Kaiser, A.; Schubert-Zsilavecz, M.; Müller, S.; Knapp, S.; Hanke, T. Synthesis of Pyrazole-Based Macrocycles Leads to a Highly Selective Inhibitor for MST3. Journal of medicinal chemistry 2024, 67 (1), 674–690. DOI: 10.1021/acs.jmedchem.3c01980.

44. Gerninghaus, J.; Zhubi, R.; Krämer, A.; Karim, M.; Tran, D. H. N.; Joerger, A. C.; Schreiber, C.; Berger, L. M.; Berger, B.-T.; Ehret, T. A. L.; Elson, L.; Lenz, C.; Saxena, K.; Müller, S.; Einav, S.; Knapp, S.; Hanke, T. Back-Pocket Optimization of 2-Aminopyrimidine-Based Macrocycles Leads to Potent EPHA2/GAK Kinase Inhibitors. Journal of medicinal chemistry 2024, 67 (15), 12534–12552. DOI: 10.1021/acs.jmedchem.4c00411.

45. Kostich, W.; Hamman, B. D.; Li, Y.-W.; Naidu, S.; Dandapani, K.; Feng, J.; Easton, A.; Bourin, C.; Baker, K.; Allen, J.; Savelieva, K.; Louis, J. V.; Dokania, M.; Elavazhagan, S.; Vattikundala, P.; Sharma, V.; Das, M. L.; Shankar, G.; Kumar, A.; Holenarsipur, V. K.; Gulianello, M.; Molski, T.; Brown, J. M.; Lewis, M.; Huang, Y.; Lu, Y.; Pieschl, R.; O’Malley, K.; Lippy, J.; Nouraldeen, A.; Lanthorn, T. H.; Ye, G.; Wilson, A.; Balakrishnan, A.; Denton, R.; Grace, J. E.; Lentz, K. A.; Santone, K. S.; Bi, Y.; Main, A.; Swaffield, J.; Carson, K.; Mandlekar, S.; Vikramadithyan, R. K.; Nara, S. J.; Dzierba, C.; Bronson, J.; Macor, J. E.; Zaczek, R.; Westphal, R.; Kiss, L.; Bristow, L.; Conway, C. M.; Zambrowicz, B.; Albright, C. F. Inhibition of AAK1 Kinase as a Novel Therapeutic Approach to Treat Neuropathic Pain. The Journal of pharmacology and experimental therapeutics 2016, 358 (3), 371–386. DOI: 10.1124/jpet.116.235333.

46. Liu, Q.; Bautista-Gomez, J.; Higgins, D. A.; Yu, J.; Xiong, Y. Dysregulation of the AP2M1 phosphorylation cycle by LRRK2 impairs endocytosis and leads to dopaminergic neurodegeneration. Science signaling 2021, 14 (693). DOI: 10.1126/scisignal.abg3555.

47. Mushtaq, I. Role Of Endocytic Machinery Regulators in EGFR Traffic and Viral Entry. Theses & Dissertations 2021.

48. Fedorov, O.; Niesen, F. H.; Knapp, S. Kinase inhibitor selectivity profiling using differential scanning fluorimetry. Methods in molecular biology (Clifton, N.J.) 2012, 795, 109–118. DOI: 10.1007/978-1-61779-337-0_7.

49. Wells, C.; Couñago, R. M.; Limas, J. C.; Almeida, T. L.; Cook, J. G.; Drewry, D. H.; Elkins, J. M.; Gileadi, O.; Kapadia, N. R.; Lorente-Macias, A.; Pickett, J. E.; Riemen, A.; Ruela-de-Sousa, R. R.; Willson, T. M.; Zhang, C.; Zuercher, W. J.; Zutshi, R.; Axtman, A. D. SGC-AAK1-1: A Chemical Probe Targeting AAK1 and BMP2K. ACS medicinal chemistry letters 2020, 11 (3), 340–345. DOI: 10.1021/acsmedchemlett.9b00399.

50. Karaman, M. W.; Herrgard, S.; Treiber, D. K.; Gallant, P.; Atteridge, C. E.; Campbell, B. T.; Chan, K. W.; Ciceri, P.; Davis, M. I.; Edeen, P. T.; Faraoni, R.; Floyd, M.; Hunt, J. P.; Lockhart, D. J.; Milanov, Z. V.; Morrison, M. J.; Pallares, G.; Patel, H. K.; Pritchard, S.; Wodicka, L. M.; Zarrinkar, P. P. A quantitative analysis of kinase inhibitor selectivity. Nature biotechnology 2008, 26 (1), 127–132. DOI: 10.1038/nbt1358.

51. Fabian, M. A.; Biggs, W. H.; Treiber, D. K.; Atteridge, C. E.; Azimioara, M. D.; Benedetti, M. G.; Carter, T. A.; Ciceri, P.; Edeen, P. T.; Floyd, M.; Ford, J. M.; Galvin, M.; Gerlach, J. L.; Grotzfeld, R. M.; Herrgard, S.; Insko, D. E.; Insko, M. A.; Lai, A. G.; Lélias, J.-M.; Mehta, S. A.; Milanov, Z. V.; Velasco, A. M.; Wodicka, L. M.; Patel, H. K.; Zarrinkar, P. P.; Lockhart, D. J. A small molecule-kinase interaction map for clinical kinase inhibitors. Nature biotechnology 2005, 23 (3), 329–336. DOI: 10.1038/nbt1068.

52. Kurz, C. G.; Preuss, F.; Tjaden, A.; Cusack, M.; Amrhein, J. A.; Chatterjee, D.; Mathea, S.; Berger, L. M.; Berger, B.-T.; Krämer, A.; Weller, M.; Weiss, T.; Müller, S.; Knapp, S.; Hanke, T. Illuminating the Dark: Highly Selective Inhibition of Serine/Threonine Kinase 17A with Pyrazolo1,5-apyrimidine-Based Macrocycles. Journal of medicinal chemistry 2022, 65 (11), 7799–7817. DOI: 10.1021/acs.jmedchem.2c00173.

53. Krämer, A.; Kurz, C. G.; Berger, B.-T.; Celik, I. E.; Tjaden, A.; Greco, F. A.; Knapp, S.; Hanke, T. Optimization of pyrazolo1,5-apyrimidines lead to the identification of a highly selective casein kinase 2 inhibitor. European Journal of Medicinal Chemistry 2020, 208, 112770. DOI: 10.1016/j.ejmech.2020.112770.

54. Ricotta, D.; Conner, S. D.; Schmid, S. L.; Figura, K. von; Honing, S. Phosphorylation of the AP2 mu subunit by AAK1 mediates high affinity binding to membrane protein sorting signals. J Cell Biol 2002, 156 (5), 791–795. DOI: 10.1083/jcb.200111068.

55. Kelly, B. T.; Graham, S. C.; Liska, N.; Dannhauser, P. N.; Höning, S.; Ungewickell, E. J.; Owen, D. J. Clathrin adaptors. AP2 controls clathrin polymerization with a membrane-activated switch. Science (New York, N.Y.) 2014, 345 (6195), 459–463. DOI: 10.1126/science.1254836.

56. Redlingshöfer, L.; Brodsky, F. M. Antagonistic regulation controls clathrin-mediated endocytosis: AP2 adaptor facilitation vs restraint from clathrin light chains. Cells & Development 2021, 168, 203714. DOI: 10.1016/j.cdev.2021.203714.

57. Schwalm, M. P.; Krämer, A.; Dölle, A.; Weckesser, J.; Yu, X.; Jin, J.; Saxena, K.; Knapp, S. Tracking the PROTAC degradation pathway in living cells highlights the importance of ternary complex measurement for PROTAC optimization. Cell Chemical Biology 2023, 30 (7), 753–765.e8. DOI: 10.1016/j.chembiol.2023.06.002.

58. Wojdyla, J. A.; Kaminski, J. W.; Panepucci, E.; Ebner, S.; Wang, X.; Gabadinho, J.; Wang, M. DA+ data acquisition and analysis software at the Swiss Light Source macromolecular crystallography beamlines. Journal of synchrotron radiation 2018, 25 (Pt 1), 293–303. DOI: 10.1107/S1600577517014503.

59. McCoy, A. J.; Grosse-Kunstleve, R. W.; Storoni, L. C.; Read, R. J. Likelihood-enhanced fast translation functions. Acta crystallographica. Section D, Biological crystallography 2005, 61 (Pt 4), 458–464. DOI: 10.1107/S0907444905001617.

60. Emsley, P.; Cowtan, K. Coot: model-building tools for molecular graphics. Acta crystallographica. Section D, Biological crystallography 2004, 60 (Pt 12 Pt 1), 2126–2132. DOI: 10.1107/S0907444904019158.

61. Murshudov, G. N.; Vagin, A. A.; Dodson, E. J. Refinement of macromolecular structures by the maximum-likelihood method. Acta crystallographica. Section D, Biological crystallography 1997, 53 (Pt 3), 240–255. DOI: 10.1107/S0907444996012255.

62. Chen, V. B.; Arendall, W. B.; Headd, J. J.; Keedy, D. A.; Immormino, R. M.; Kapral, G. J.; Murray, L. W.; Richardson, J. S.; Richardson, D. C. MolProbity: all-atom structure validation for macromolecular crystallography. Acta crystallographica. Section D, Biological crystallography 2010, 66 (Pt 1), 12–21. DOI: 10.1107/S0907444909042073.

63. Chernobrovkin, A. L.; Cázares-Körner, C.; Friman, T.; Caballero, I. M.; Amadio, D.; Martinez Molina, D. A Tale of Two Tails: Efficient Profiling of Protein Degraders by Specific Functional and Target Engagement Readouts. SLAS discovery : advancing life sciences R & D 2021, 26 (4), 534–546. DOI: 10.1177/2472555220984372.

64. Zou, G.; Xu, H. Y.; Qing, M.; Wang, Q.-Y.; Shi, P.-Y. Development and characterization of a stable luciferase dengue virus for high-throughput screening. Antiviral Research 2011, 91 (1), 11–19. DOI: 10.1016/j.antiviral.2011.05.001.

65. Sun, C.; Gardner, C. L.; Watson, A. M.; Ryman, K. D.; Klimstra, W. B. Stable, high-level expression of reporter proteins from improved alphavirus expression vectors to track replication and dissemination during encephalitic and arthritogenic disease. Journal of virology 2014, 88 (4), 2035–2046. DOI: 10.1128/JVI.02990-13.

66. Chiem, K.; Morales Vasquez, D.; Park, J.-G.; Platt, R. N.; Anderson, T.; Walter, M. R.; Kobie, J. J.; Ye, C.; Martinez-Sobrido, L. Generation and Characterization of recombinant SARS-CoV-2 expressing reporter genes. Journal of virology 2021, 95 (7). DOI: 10.1128/JVI.02209-20.

67. Saul, S.; Karim, M.; Ghita, L.; Huang, P.-T.; Chiu, W.; Durán, V.; Lo, C.-W.; Kumar, S.; Bhalla, N.; Leyssen, P.; Alem, F.; Boghdeh, N. A.; Tran, D. H.; Cohen, C. A.; Brown, J. A.; Huie, K. E.; Tindle, C.; Sibai, M.; Ye, C.; Khalil, A. M.; Chiem, K.; Martinez-Sobrido, L.; Dye, J. M.; Pinsky, B. A.; Ghosh, P.; Das, S.; Solow-Cordero, D. E.; Jin, J.; Wikswo, J. P.; Jochmans, D.; Neyts, J.; Jonghe, S. de; Narayanan, A.; Einav, S. Anticancer pan-ErbB inhibitors reduce inflammation and tissue injury and exert broad-spectrum antiviral effects. The Journal of clinical investigation 2023, 133 (19). DOI: 10.1172/JCI169510

