## Supporting Information for "Development of pyrazolo[1,5-a]pyrimidine based macrocyclic kinase inhibitors targeting AAK1"

**Table of Content:**

Supplementary Tables **S1** – **S6**

### Materials and Methods

#### Chemistry

### Analytical data of compounds **2-30**

**Table S1:** DSF assay of compounds **14-30** and control compounds **LP-935509, AAK1-SGC-1N, AAK1-SGC, SGC-GAK-1N** and **SGC-GAK** against the three NAK family members AAK1, BIKE and GAK, the corresponding ΔTm-shifts given in percentage shift of the respective ΔTm-shift of the reference compound Staurosporine.

| **Compound** | **AAK1 [%]** | **BIKE [%]** | **GAK [%]** |
| --- | --- | --- | --- |
| **14** | 58 | 50 | 48 |
| **15** | 77 | 81 | 89 |
| **16** | 61 | 11 | 21 |
| **17** | 50 | 22 | 21 |
| **18** | 61 | 34 | 39 |
| **19** | 31 | 11 | 12 |
| **20** | 32 | 19 | 17 |
| **21** | 67 | 27 | 44 |
| **22** | 76 | 37 | 72 |
| **23** | 71 | 51 | 59 |
| **24** | 60 | 60 | 59 |
| **25** | 61 | 25 | 21 |
| **26** | 67 | 72 | 71 |
| **27** | 79 | 45 | 38 |
| **28** | 54 | 53 | 41 |
| **29** | 61 | 62 | 63 |
| **30** | 69 | 32 | 74 |
| **AAK1-SGC-1N** | 17 | 14 | 5 |
| **AAK1-SGC** | 61 | 66 | 13 |
| **SGC-GAK-1N** | 2 | 3 | 10 |
| **SGC-GAK** | 1 | 2 | 84 |
| **LP-935509** | 90 | 77 | 41 |

**Table S2:** DSF assay of compounds **14-30** against a panel of 89 kinases with their corresponding ΔTm-shifts, given in percentage shift of the respective Δ Tm-shift of a reference compound. (*Excel*)

**Table S3:** Determination of EC_50_ values for **16**, **22-24**, **27–28**, **LP-935509** and **Staurosporine** in a cellular NanoBRET target engagement assay in intact cells against STK16.

| **Compound** | **Average [M]** | **Average [µM]** | **Standard deviation [µM]** |
| --- | --- | --- | --- |
| **16** | 7.8E-07 | 0.8 | 0.1 |
| **22** | 7.9E-07 | 0.8 | 0.1 |
| **23** | 5.3E-06 | 5.3 | 0.2 |
| **24** | 6.0E-06 | 6.0 | 0.6 |
| **27** | 5.6E-06 | 5.6 | 0.5 |
| **28** | 8.5E-06 | 8.5 | 0.1 |
| **LP-935509** | 6.5E-05 | > 25 | / |
| **Staurosporine** | 8.3E-07 | 0.8 | 0.1 |

**Table S4: KINOMEscan®** of **16, 18** and **27** @ 100 nM, 1 μM and 100 nM respectively, against 409 kinases from **Eurofins** (formerly DiscoverX). (*Excel*)

**Table S5**: CETSA shifts of **16** and **20 – 30,** conducted by **PELAGO BIOSCIENCE.** (*Excel*)

**Table S6:** Determination of EC_50_ values of compounds **14-30, SGC-AAK1-1, SGC-AAK1-1N, SGC-GAK-1, SGC-GAK-1N, LP-935509** in a cellular NanoBRET target engagement assay in intact and lysed cells against AAK1, GAK and BIKE (BMP2K). (*Excel*)

**Materials and Methods**

**Chemistry**

The synthesis of compounds will be explained in the following and the analytical data for them can be found in the Supporting Information. All commercial chemicals were purchased from common suppliers with a purity ≥ 95% and were used without further purification. The solvents with an analytical grade were obtained from VWR Chemicals and Merck and all dry solvents from Acros Organics. All reactions were proceeded under an argon atmosphere. The thin layer chromatography was done with silica gel on aluminum foils (60 Å pore diameter) obtained from Macherey-Nagel and visualized with ultraviolet light (λ = 254 and 365 nm). The purification of the compounds was done by flash chromatography. A puriFlash XS 420 device with a UV-VIS multiwave detector (200−400 nm) from Interchim was used with pre-packed normal-phase PF-SIHP silica columns with particle sizes of 15 and 30 μm (Interchim). Preparative purification by HPLC was carried out on an Agilent 1260 Infinity II device using an Eclipse XDB-C18 (Agilent, 21.2 x 250mm, 7µm) reversed phase column. A suitable gradient (flow rate 21 ml/min.) was used, with 0.1% TFA in water (A) and 0.1% TFA S25 in acetonitrile (B), as a mobile phase. The nuclear magnetic resonance spectroscopy (NMR) was performed with DPX250, AV300, AV400 or AV500 MHz spectrometers from Bruker. Chemical shifts (δ) are reported in parts per million (ppm). DMSO-d6, chloroform-d and methylene chloride-d2 was used as a solvent, and the spectra were calibrated to the solvent signal: 2.50 ppm (1H NMR) or 39.52 ppm (13C NMR) for DMSO-d6, 7.26 ppm (1H NMR) or 77.16 ppm (13C NMR) for chloroform-d and 5.32 ppm (1H NMR) or 54.00 ppm (13C NMR) for methylene chloride-d2. Coupling constants (J) were reported in hertz (Hz) and multiplicities were designated as followed: s (singlet), d (doublet), dd (doublet of doublet), t (triplet), dt (doublet of triplets), td (triplet of doublets), ddd (doublet of doublet of doublet), q (quartet), m (multiplet). Mass spectra were measured on a Surveyor MSQ device from ThermoFisher measuring in the positive- or negative-ion mode. Final compounds were additionally characterized by HRMS using a MALDI LTQ Orbitrap XL from ThermoScientific. The purity of the final compounds was determined by HPLC using an Agilent 1260 Infinity II device with a 1260 DAD HS detector (G7117C; 254 nm, 280 nm, 310 nm) and a LC/MSD device (G6125B, ESI pos. 100-1000). The compounds were analyzed on a Poroshell 120 EC-C18 (Agilent, 3 x 150 mm, 2.7 µm) reversed phase column using 0.1% formic acid in water (A) and 0.1% formic acid in acetonitrile (B) as a mobile phase. The following gradient was used: 0 min 5% B - 2 min 5% B - 8 min 98% B - 10 min 98% B (flow rate of 0.5 mL/min). UV-detection was performed at 254, 280 and 310 nm and all compounds used for further biological characterizations showed a purity ≥95%. No unexpected or unusually high safety hazards were encountered.

**Synthesis of 3-bromo-5-chloropyrazolo[1,5-*a*]pyrimidine (2)**

5,7-dichloropyrazolo[1,5-*a*]pyrimidine (20.00 g, 1.0 eq , 130.23 mmol) (**1)** was suspended in 250 mL anh. ACN. Then NBS (25.50 g, 1.1 eq, 143.26 mmol) was added to the suspension, which was subsequently stirred at rt for 3 hrs. The removal of the solvent under reduced pressure was followed by the addition of 300 mL H_2_O to the residue. The precipitant was filtered, washed, and dried under reduced pressure, to obtain **2** as a brown solid (28.90 g, 96%).

^1^H NMR (250MHz, DMSO-d_6_): δ 9.21 (d, J = 7.3 Hz, 1H, HetH), 8.43 (s, 1H, HetH), 7.22 (d, J=7.3 Hz, 1H, HetH) ppm.

**Synthesis of 2-[2-({3-bromopyrazolo[1,5-*a*]pyrimidine-5 yl}amino)ethoxy]ethan-1-ol (3a)**

To a solution of 3-bromo-5-chloropyrazolo [1,5-*a*]pyrimidine (28.90 g, 1.0 eq, 124.32 mmol) (**2**) in 380 mL acetonitrile (anh.), 2-(2-aminoethoxy)ethanol (7.46 g, 1.1 eq, 136.75 mmol) and *N,N*-diisopropylethylamine (10.01 g, 1.2 eq, 149.18 mmol) were added. The obtained solution was stirred under reflux for 16 hrs. Afterwards, the solvent was evaporated, and the residue was taken up in ethyl acetate and washed several times with water and brine. After drying over MgSO_4_ the solvent was evaporated under reduced pressure and the crude product was purified with silica flash column chromatography with a mobile phase of hexanes and ethyl acetate (ratio gradually ranging from 3:1 to 0:1). **3a** was obtained as a yellow oil (30.32 g, 81%).

1H NMR (250 MHz, DMSO-*d6*): *δ* 8.45 (d, *J* = 7.5 Hz, 1H Het*H*), 7.87 (s, 1H, Het*H*), 7.70 (t, *J* = 5.0 Hz, 1H, N*H*), 6.34 (d, *J* = 7.6 Hz, 1H, Het*H*), 4.57 (s, 1H, O*H*), 3.66 – 3.45 (m, 8H, C*H*2) ppm.

**Synthesis of 6-((3-bromopyrazolo[1,5-*a*]pyrimidin-5-yl)amino)hexan-1-ol (3b)**

To a solution of 3-bromo-5-chloropyrazolo [1,5-*a*]pyrimidine (0.70 g, 1.0 eq, 3.01 mmol) (**2**) in acetonitrile (anh.), 6-aminohexan-1-ol (0.423 g, 1.2 eq, 3.61 mmol) and *N,N*-diisopropylethylamine (0.63 mL, 1.2 eq, 3.61 mmol) were added. The obtained solution was stirred under reflux for 16 hrs. Afterwards, the solvent was evaporated, and the residue was taken up in ethyl acetate and washed several times with water and brine. After drying over MgSO_4_ the solvent was evaporated under reduced pressure and the crude product was purified with silica flash column chromatography with a mobile phase of hexanes and ethyl acetate (ratio gradually ranging from 3:1 to 0:1). **3b** was obtained as a yellow oil (0.656 g, 70%).

1H NMR (250 MHz, DMSO-d6): δ 8.42 (d, J = 7.4 Hz, 1H, HetH), 7.85 (s, 1H, HetH), 7.62 (t, J = 5.3 Hz, 1H, NH), 6.28 (d, J = 7.6 Hz, 1H, HetH), 4.32 (t, J = 5.1 Hz, 1H OH), 3.44 – 3.32 (m, 4H, CH2), 1.65 – 1.50 (m, 2H, CH2), 1.49 – 1.26 (m, 6H, CH2) (ppm).

**Synthesis of 3-bromo-5-(8,8,9,9-tetramethyl-4,7-dioxa-1-aza-8-siladecan-1-yl)pyrazolo-[1,5-*a*]pyrimidine (4a)**

2-[2-({3-Bromopyrazolo[1,5-*a*]pyrimidine-5-yl}amino)ethoxy]ethan-1-ol (**3a**) (30.30 g,1.0 eq, 100.62 mmol) and triethylamine (20.36 g, 2.0 eq, 201.24 mmol) were dissolved in dry dimethylformamide (280 mL). While stirring, *tert*-butyldimethylsilyl chloride (22.75 g, 1.5 eq, 150.93 mmol) was added in small portions. After stirring for 1 h at room temperature, the solvent was removed under reduced pressure and the residue was dissolved in ethyl acetate. The organic layer was washed with water and brine. Afterwards it was dried over MgSO4, the solvent was removed under reduced pressure and further purificated by a silica gel column chromatography with a mobile phase of *n-*hexane an ethyl acetate (ratio of 1:1). The title compound was obtained as a white solid (31.40 g, 75%). 1H NMR (250 MHz, DMSO-d6): δ 8.44 (d, J = 7.6 Hz, 1H, HetH), 7.86 (s, 1H, HetH), 7.70 (t, J = 5.4 Hz, 1H, NH), 6.33 (d, J = 7.6 Hz, 1H, HetH), 3.74 – 3.67 (m, 2H, CH2), 3.64 – 3.48 (m, 6H, CH2), 0.83 (s, 9H, (CH3)3), 0.02 (s, 6H, Si(CH3)2) ppm.

**Synthesis of 3-bromo-N-(6-((tert-butyldimethylsilyl)oxy)hexyl)pyrazolo-[1,5-a]-pyrimidin-5-amine (4b)**

6-((3-bromopyrazolo[1,5-*a*]pyrimidin-5-yl)amino)hexan-1-ol (**3b**) (0.656 g, 1.0 eq, 2.09 mmol). and triethylamine (0.57 mL, 2.0 eq, 4.19 mmol) were dissolved in dry dimethylformamide (20 mL). While stirring, *tert*-butyldimethylsilyl chloride (0.475 g, 1.5 eq, 3.14 mmol) was added in small portions. After 1 h at room temperature the solvent was removed under reduced pressure and the residue was dissolved in water. The aqueous layer was extracted with ethyl acetate, the organic layer washed with brine and dried over MgSO4. The solvent was removed under reduced pressure and further purification was carried out by a silica gel column chromatography with a mobile phase of *n-*hexane an ethyl acetate (ratio of 1:1). The title compound was obtained as a white solid (0.79 g, 88%). 1H NMR (250 MHz, DMSO-d6): δ 8.42 (d, J = 7.4 Hz, 1H, HetH), 7.85 (s, 1H, HetH), 7.62 (t, J = 5.3 Hz, 1H, NH), 6.28 (d, J = 7.6 Hz, 1H, HetH), 4.32 (t, J = 5.1 Hz, 1H OH), 3.44 – 3.32 (m, 4H, CH2), 1.65 – 1.50 (m, 2H, CH2), 1.49 – 1.26 (m, 6H, CH2) (ppm).

**Synthesis of tert-butyl N-{3-bromopyrazolo[1,5-a]pyrimidin-5-yl}-N-(2-{2-[(tert-butyl-dimethylsilyl)oxy]ethoxy}ethyl)carbamate (5a)**

To a solution of 3-bromo-5-(8,8,9,9- tetramethyl-4,7-dioxa-1-aza-8-siladecan-1-yl)pyrazolo[1,5-a]pyrimidine (**4a**) (29.23 g, 1.0 eq, 70.37 mmol), di-tert-butyldicarbonate (38.39 g, 2.50 eq, 175.91 mmol) and 4(dimethylamino) pyridine (171.93 g, 0.02 eq, 1.41 mmol) in tetrahydrofuran (260 mL), Triethylamine (8.54 g, 1.2 eq, 84.44 mmol) was added. The reaction mixture was heated to reflux for 3 h and afterwards, the solvent was evaporated. The residue was taken up in ethyl acetate and washed with water and brine. After drying over MgSO4 the organic layer was removed under reduced pressure. Further purification occurred by silica gel column chromatography with a mobile phase of n-hexane an ethyl acetate (ratio gradually ranging from 1:0 to 0:1). The desired compound was yielded as a yellowish oil (34.50 g, 95%). 1H NMR (250 MHz, DMSO-d6): δ 8.95 (d, J = 7.8 Hz, 1H, HetH), 8.24 (s, 1H, HetH), 7.42 (d, J = 7.8 Hz, 1H, HetH), 4.18 (t, J = 6.1 Hz, 2H, CH2), 3.69 (t, J = 6.0f Hz, 2H, CH2), 3.61 – 3.55 (m, 2H, CH2), 3.50 – 3.44 (m, 2H, CH2), 1.51 (s, 9H, (CH3)3), 0.79 (s, 9H, (CH3)3), -0.06 (s, 6H, Si(CH3)2) ppm.

**Synthesis of tert-butyl (3-bromopyrazolo[1,5-a]pyrimidin-5-yl)(6-((tertbutyldimethylsilyl) oxy)hexyl)carbamate (5b)**

Triethylamine (0.302 mL, 1.2 eq, 2.22 mmol) was added to a solution of 3-bromo-N-(6-((tert-butyldimethylsilyl)oxy)hexyl)pyrazolo-[1,5-a]-pyrimidin-5-amine (0.79 g, 1.0 eq, 1.85 mmol), di-tert-butyldicarbonate (1.01 g, 2.50 eq, 4.62 mmol) and 4(dimethylamino) pyridine (0.005 g, 0.02 eq, 0.04 mmol) in tetrahydrofuran (20 mL). The reaction mixture was heated to reflux for 3 h and afterwards, the solvent was evaporated. The residue was taken up in ethyl acetate and washed with water and brine. After drying over MgSO4 the organic layer was removed under reduced pressure. Further purification was done with silica gel column chromatography with a mobile phase of n-hexane an ethyl acetate (ratio gradually ranging from 1:0 to 0:1). The desired compound was yielded as a yellowish oil (0.93 g, 95%). 1H NMR (250 MHz, DMSO-d6): δ 8.94 (d, J = 7.8 Hz, 1H, HetH), 8.23 (s, 1H, HetH), 7.48 (d, J = 7.8 Hz, 1H, HetH), 4.02 – 3.92 (m, 2H, CH2), 3.55 (t, J = 6.1 Hz, 2H, CH2), 1.75 – 1.60 (m, 2H, CH2), 1.51 (s, 9H, C(CH3)3), 1.48 – 1.40 (m, 2H, CH2), 1.37 – 1.27 (m, 4H, CH2), 0.83 (s, 9H, C(CH3)3), -0.01 (s, 6H, Si(CH3)2) ppm.

**Synthesis of Methyl-4-bromo-2-hydroxybenzoate (7)**

4-bromo-2-hydroxybenzoic acid (2.00 g, 1.0 eq, 9.22 mmol) (**6**) was dissolved in 20 mL anh. MeOH. H_2_SO_4_ (0.1 mL, 1.85 mmo1, 0.2 eq) was then added and the reaction solution was stirred at °C for 65 h. Afterwards, the solvent was removed under reduced pressure and the crude product extracted with a sat. NaHCO_3_ solution. The organic phase was dried over MgSO_4_ and the solvent removed under reduced pressure. The product was obtained as an off-white solid (1.34 g, 63 %). 1H NMR (250 MHz, DMSO-d6): δ 10.89 (s, 1H), 7.93 (d = 8.8 Hz, 1H), 7.14 (d, J =2.5 Hz, 1H), 1.06 (dd, J = 8.8, 2.5 Hz, 1H), 3.90 (d, J = 6.3 Hz, 3H) ppm.

**Synthesis of methyl 2-hydroxy-4(4,4,5,5-tetramethyl-1,3,2-dioxaborolan-2-yl)benzoate (8)**

To a solution of Methyl-4-bromo-2-hydroxybenzoate (1.34 g, 1.0 eq, 5.81 mmol) (**7**) in dry 1,4-dioxane (20 mL), bis(pinacolato)diboron (1.48 g, 1.0 eq, 5.81 mmol), potassium acetate (1.71 g, 3.0 eq, 17.44 mmol) and Pd(dppf)Cl_2_·DCM (0.213 g, 0.05 eq, 0.290 mmol) were added. The reaction solution was flushed with argon for 10 min and stirred at 100 °C for 3.5 h. Subsequently the reaction was diluted with ethyl acetate and water and the layers were separated. The organic layer was washed with brine and dried over MgSO_4_. After removal of the solvent under reduced pressure, the crude product was purified with silica gel column chromatography with a mobile phase of n-hexane and ethyl acetate (ratio gradually ranging from 1:0 to 30:1). The title compound was obtained as a white solid (1.47 g, 91%). 1H NMR (250 MHz, DMSO-d6): δ 10.35 (s, 1H, OH), 7.76 (d, J = 7.8 Hz, 1H, PhH), 7.21 – 7.18 (m, 2H, PhH), 3.88 (s, 3H, OCH3), 1.30 (s, 12H, CH3) ppm.

**Synthesis of methyl 4-(5-((tert-butoxycarbonyl)(2-(2-((tert-butyldimethylsilyl)oxy)-ethoxy)ethyl)amino)pyrazolo[1,5-a]pyrimidin-3-yl)-2-hydroxybenzoate (9a)**

tert-Butyl-N-{3-bromopyrazolo[1,5-a]pyrimidin-5-yl}-N-(2-{2-[(tert-butyldimethylsilyl) oxy]ethoxy}ethyl)carbamate (**5a**) (4.63 g, 1.0 eq, 8.99 mmol), methyl 2-hydroxy- 4(4,4,5,5-tetramethyl-1,3,2-dioxaborolan-2-yl)benzoate (**8**) (5.00 g, 2.0 eq 17.98 mmol), Pd2(dba)3 (823 mg, 0.1 eq, 0.90 mmol), XPhos (429 mg, 0.1 eq, 0.90 mmol) and potassium phosphate (7.63 g, 4.0 eq, 35.96 mmol) were suspended in 1,4-dioxane (50 mL) and water (20 mL). The reaction was stirred at 110 °C for 40 min. After diluting with ethyl acetate, the layers were separated. The organic phase was washed with water and brine and dried over MgSO4. The solvent was evaporated in vacuo. Purification was performed by flash silica gel column chromatography with a mobile phase of n-hexane an ethyl acetate (ratio gradually ranging from 1:0 to 0:1). The bright yellow oil obtained was the desired compound (4.85 g, 91%). 1H NMR (250 MHz, DMSO-d6): δ 10.62 (s, 1H, OH), 8.99 (d, J = 7.8 Hz, 1H, HetH), 8.74 (s, 1H, HetH), 7.79 (d, J = 8.2 Hz, 1H, PhH), 7.71 – 7.66 (m, 2H, PhH), 7.49 (d, J = 7.8 Hz, 1H, HetH), 4.23 (t, J = 6.2 Hz, 2H, CH2), 3.90 (s, 3H, OCH3), 3.77 (t, J = 6.2 Hz, 2H, CH2), 3.61 (t, J = 5.2 Hz, 2H, CH2), 3.48 (t, J = 5.0 Hz, 2H, CH2), 1.53 (s, 9H, (CH3)3), 0.76 (s, 9H, (CH3)3), -0.08 (s, 6H, Si(CH3)2) ppm.

**Synthesis of methyl 4-(5-((tert-butoxycarbonyl)(6-((tert-butyldimethylsilyl)-oxy)hexyl)-amino)pyrazolo[1,5-a]pyrimidin-3-yl)-2-hydroxybenzoate (9b)**

tert-butyl(3-bromopyrazolo[1,5-a]pyrimidin-5-yl)(6-((tertbutyldimethylsilyl)oxy)hexyl) carbamate (0.93 g, 1.0 eq, 1.76 mmol) (**5b**),methyl 2-hydroxy- 4(4,4,5,5-tetramethyl-1,3,2-dioxaborolan-2-yl)benzoate (**8**) (0.59 g, 1.2 eq 2.12 mmol), Pd2(dba)3 (0.07 g, 0.05 eq, 0.09 mmol), XPhos (0.04 g, 0.05 eq, 0.09 mmol) and potassium phosphate (0.94 g, 2.5 eq, 4.41 mmol) were suspended in 1,4-dioxane (10 mL) and water (10 mL). The reaction was stirred at 110 °C for 40 min. After diluting with ethyl acetate, the layers were separated. The organic one was washed with water and brine and dried over MgSO_4_. The solvent was evaporated in vacuo. Purification was performed by flash silica gel column chromatography with a mobile phase of n-hexane an ethyl acetate (ratio gradually ranging from 1:0 to 0:1). The compound was obtained as bright yellow oil (0.91 g, 91%). 1H NMR (250 MHz, CDCl3-d): δ 10.76 (s, 1H, OH), 8.46 (d, J = 7.9 Hz, 1H, HetH), 8.38 (s, 1H, HetH), 7.83 (d, J = 8.4 Hz, 1H, PhH), 7.72 (s, 1H, PhH), 7.70 (d, J = 5.9 Hz, 1H, HetH), 7.55 (dd, J = 8.4, 1.7 Hz, 1H, PhH), 4.14 – 4.08 (m, 2H, CH2), 3.95 (s, 3H, OCH3), 3.60 (t, J = 6.4 Hz, 2H, CH2), 1.89 – 1.77 (m, 2H, CH2), 1.58 (s, 9H, C(CH3)3), 1.56 – 1.39 (m, 6H, CH2), 0.87 (s, 9H, C(CH3)3), 0.02 (s, 6H, Si(CH3)2) (ppm).

**Synthesis of methyl 4-(5-{[(tert-butoxy)carbonyl][2-(2-hydroxyethoxy)-ethyl]amino}-pyrazolo[1,5-a]pyrimidin-3-yl)-2-hydroxybenzoate (10a)**

To a solution of methyl 4-(5-((tert-butoxycarbonyl)(2-(2-((tert-butyldimethylsilyl)oxy)- ethoxy)ethyl)amino)pyrazolo[1,5-a]pyrimidin-3-yl)-2-hydroxybenzoate (**9a**) (4.13 g, 1.0 eq, 7.04 mmol) in tetrahydrofuran (60 mL) 1 M tetrabutylammonium fluoride solution (10.6 mL, 10.56 mmol) in tetrahydrofuran) was added. The reaction was stirred at room temperature. After 3 h the solvent was removed under reduced pressure. Subsequently the residue was taken up in ethyl acetate, washed with water and brine and dried over MgSO_4_. Following, the solvent was evaporated under reduced pressure. Further purification occurred through flash silica gel column chromatography with a mobile phase of n-hexane an ethyl acetate (ratio gradually ranging from 1:0 to 0:1). The title compound was yielded as yellow solid (2.81 g, 85%). 1H NMR (500 MHz, DMSO-d6): δ 10.62 (s, 1H, OH), 8.99 (d, J = 7.8 Hz, 1H, HetH), 8.75 (s, 1H, HetH), 7.81 (d, J = 8.3 Hz, 1H, PhH), 7.71 – 7.68 (m, 2H, PhH), 7.49 (d, J = 7.8 Hz, 1H, HetH), 4.52 (t, J = 5.1 Hz, 1H, OH), 4.22 (t, J = 6.3 Hz, 2H, CH2), 3.91 (s, 3H, OCH3), 3.77 (t, J = 6.3 Hz, 2H, CH2), 3.48 – 3.42 (m, 4H, CH2), 1.53 (s, 9H, (CH3)3) ppm.

13C NMR (126 MHz, DMSO-d6): δ 169.44, 160.83, 153.83, 152.82, 143.78, 143.09, 139.79, 136.58, 130.15, 116.35, 112.68, 109.33, 106.19, 104.73, 82.53, 72.24, 67.98, 60.21, 52.34, 45.82, 27.64 ppm.

**Synthesis of methyl 4-(5-((tert-butoxycarbonyl)(6-hydroxyhexyl)amino)-pyrazolo[1,5-a]-pyrimidin-3-yl)-2-hydroxybenzoate (10b)**

To a solution of methyl 4-(5-((tert-butoxycarbonyl)(6-((tert-butyldimethylsilyl)-oxy)hexyl)-amino)pyrazolo[1,5-a]pyrimidin-3-yl)-2-hydroxybenzoate (**9b**) (0.91 g, 1.0 eq, 1.52 mmol) in tetrahydrofuran (20 mL) was added a 1 M tetrabutylammonium fluoride solution (2.31 mL, 1.5 eq, 2.31 mmol) in anh. tetrahydrofuran. The reaction was stirred at room temperature. After 3 h the solvent was removed under reduced pressure. Subsequently the residue was taken up with ethyl acetate, washed with water and brine and the solvent was evaporated again. Further purification was done by flash silica gel column chromatography with a mobile phase of n-hexane an ethyl acetate (ratio gradually ranging from 1:0 to 0:1). The title compound was yielded as yellow solid (0.72 g, 98%). 1H NMR (250 MHz, CDCl3-d): δ 10.78 (s, 1H, OH), 8.47 (d, J = 7.9 Hz, 1H, HetH), 8.38 (s, 1H, HetH), 7.83 (d, J = 8.4 Hz, 1H, PhH), 7.75 (d, J = 1.7 Hz, 1H, PhH), 7.70 (d, J = 7.9 Hz, 1H, HetH), 7.51 (dd, J = 8.4, 1.7 Hz, 1H, PhH), 4.19 – 4.06 (m, 2H, CH2), 3.95 (s, 3H, OCH3), 3.65 (t, J = 6.3 Hz, 2H, CH2), 1.94 – 1.77 (m, 2H, CH2), 1.67 – 1.60 (m, 2H, CH2), 1.58 (s, 9H, C(CH3)3), 1.57 – 1.34 (m, 6H, CH2) (ppm).

**Synthesis of 9-(tert-butyl) 24-methyl (13Z,14E)-3,6-dioxa-9-aza-1(3,5)- pyrazolo[1,5-a]pyrimidina-2(1,3)-benzenacyclononaphane-24,9- dicarboxylate (11a)**

To a suspension of triphenylphosphine (832 mg, 3.0 eq 3.17 mmol) and sodium sulfate in dry toluene (240 mL) was added a solution of diisopropyl azodicarboxylate (642 mg, 3.0 eq, 3.17 mmol) in dry toluene (75 mL), dropwise. Afterwards methyl 4-(5-{[(tert-butoxy)carbonyl][2-(2-hydroxyethoxy)ethyl]amino}pyrazolo[1,5-a]pyrimidin- 3-yl)-2-hydroxybenzoate (**10a**) (500 mg, 1.0 eq, 1.06 mmol) was dissolved in 2-methyltetrahydrofuran (50 mL) and this solution was added dropwise. The reaction was stirred for 16 h at 90 °C under argon atmosphere. Sodium sulfate then was separated by filtration and the solvent was evaporated in vacuo. The purification was achieved by silica gel column chromatography with a mobile phase of n-hexane an ethyl acetate (ratio gradually ranging from 9:1 to 0:1). The title compound was obtained as a yellowish solid (417 mg, 87%).

1H NMR (500 MHz, DMSO-d6): δ 9.01 (d, J = 7.8 Hz, 1H, HetH), 8.75 (d, J = 1.2 Hz, 1H, PhH), 8.71 (s, 1H, HetH), 7.69 (d, J = 8.1 Hz, 1H, PhH), 7.55 (d, J = 7.8 Hz, 1H, HetH), 7.43 (dd, J = 8.1, 1.4 Hz, 1H, PhH), 4.39 (t, J = 6.0 Hz, 2H, CH2), 4.09 (t, J = 6.6 Hz, 2H, CH2), 3.92 (t, J = 6.6 Hz, 2H, CH2), 3.88 (t, J = 6.0 Hz, 2H, CH2), 3.78 (s, 3H, OCH3), 1.54 (s, 9H, (CH3)3) ppm.

13C NMR (126 MHz, DMSO-d6): δ 165.78, 158.86, 153.40, 152.43, 143.08, 142.97, 137.60, 136.52, 131.23, 116.61, 112.64, 106.34, 104.16, 82.78, 67.51, 66.80, 66.57, 51.69, 46.57, 27.66 ppm.

MS (ESI+) m/z: 455.03 [M + H]+.

HRMS m/z: [M + H]+ calcd for C23H27N4O6, 455.19251; found 455.19082.

HPLC (III): tR = 16.694, purity ≥ 95%.

**Synthesis of 10-(tert-butyl) 24-methyl (13Z,14E)-3-oxa-10-aza-1(3,5)- pyrazolo[1,5-a]pyrimidina-2(1,3)-benzenacyclodecaphane-24,10-dicarboxylate (11b)**

A suspension of triphenylphosphine (TPP) (1,17 g, 3.0 eq 4.46 mmol) and sodium sulfate in dry toluene (240 mL) was added dropwise with a solution of diisopropyl azodicarboxylate (DIAD) (0.88 mL, 3.0 eq, 4.46 mmol) in dry toluene (75 mL). Afterwards methyl 4-(5-((tert-butoxycarbonyl)(6-hydroxyhexyl)amino)-pyrazolo[1,5-a]-pyrimidin-3-yl)-2-hydroxybenzoate (**10b**) (0.72 mg, 1.0 eq, 1.49 mmol) was dissolved in 2-methyltetrahydrofuran (50 mL) and was added dropwise to the TPP and DIAD solution. Reaction took place over 16 h at 90 °C under argon atmosphere. Sodium sulfate was separated by filtration and the solvent was evaporated in vacuo. Purification was achieved by silica gel column chromatography with a mobile phase of n-hexane an ethyl acetate (ratio gradually ranging from 9:1 to 0:1). The yellowish solid obtained was the title compound (0.48 g, 70%).

1H NMR (500 MHz, DMSO-d6): δ 8.98 – 8.90 (m, 1H, HetH), 8.72 – 8.66 (m, 1H, HetH), 8.00 – 7.93 (m, 1H, PhH), 7.71 – 7.66 (m, 1H, PhH), 7.52 – 7.46 (m, 1H, HetH), 7.46 – 7.41 (m, 1H, PhH), 4.24 – 4.14 (m, 2H, CH2), 3.90 – 3.82 (m, 2H, CH2), 3.78 (s, 3H, OCH3), 1.91 – 1.75 (m, 4H, CH2), 1.52 (s, 9H, C(CH3)3), 1.48 (s, 4H, CH2) (ppm).

13C NMR (126 MHz, DMSO-d6): δ 165.84, 158.04, 154.02, 152.62, 143.76, 142.80, 137.61, 136.29, 131.52, 117.27, 116.94, 110.86, 106.52, 104.31, 82.33, 68.35, 51.65, 46.83, 27.65, 24.33, 24.02, 23.56, 21.42 (ppm).

MS (ESI+) m/z: 466.60 [M + H]+.

HRMS m/z: [M + Na]+ calcd for C25H30N4O5Na, 489.21084; found 489.21036.

HPLC (I): tR = 19.128, purity ≥ 95%.

**Synthesis of (13Z,14E)-3,6-dioxa-9-aza-1(3,5)-pyrazolo[1,5-a]-pyrimidina-2(1,3)-benzenacyclononaphane-24-carboxylate (12a)**

To a solution of 9-(tert-butyl) 24-methyl (13Z,14E)-3,6-dioxa-9-aza-1(3,5)- pyrazolo[1,5-a]pyrimidina-2(1,3)-benzenacyclononaphane-24,9-dicarboxylate (**11a**) (1.05 g, 1.0 eq, 2.31 mmol) in dry methylene chloride (40 mL), trifluoroacetic acid (8.3 mL g, 47.0 eq, 108.59 mmol) was added dropwise and the mixture was stirred for 16 h at room temperature. Afterwards the solvent was removed under reduced pressure and the residue was taken up in ethyl acetate. The organic phase was washed with saturated potassium carbonate solution and the aqueous phase was extracted with ethyl acetate. The combined organic phases were washed with brine. Subsequently, the solution was filtrated, and the solvent was removed again. The residue was taken up with ethyl acetate, it was washed with water and brine once more and dried over MgSO4. After removal of the organic solvent under vacuum, the purification occured by flash column chromatography with a mobile phase of ethyl acetate (isocratic ratio). The title compound was yielded as a white solid (0.69 g, 84%). 1H NMR (500 MHz, DMSO-d6): δ 8.83 (d, J = 1.5 Hz, 1H, HetH), 8.57 (d, J = 7.6 Hz, 1H, HetH), 8.39 (s, 1H, PhH), 7.94 (t, J = 5.4 Hz, 1H, NH), 7.70 (d, J = 8.1 Hz, 1H, PhH), 7.29 (dd, J = 8.1, 1.4 Hz, 1H, PhH), 6.34 (d, J = 7.6 Hz, 1H, HetH), 4.37 (t, J = 7.9 Hz, 2H, CH2), 4.00 – 3.97 (m, 2H, CH2), 3.89 – 3.86 (m, 2H, CH2), 3.76 (s, 3H, OCH3), 3.56 – 3.50 (m, 2H, CH2) ppm.

13C NMR (126 MHz, DMSO-d6): δ 165.66, 158.08, 156.18, 145.33, 141.81, 139.04, 135.82, 131.61, 115.64, 114.82, 110.45, 103.40, 100.21, 65.35, 65.27, 51.57 ppm.

MS (ESI+) m/z: 354.96 [M + H]+.

HRMS m/z: [M + H]+ calcd for C18H19N4O4, 355.14008; found 355.14078.

HPLC (III): tR = 12.868, purity ≥ 95%.

**Synthesis of Synthesis of methyl (13Z,14E)-3-oxa-10-aza-1(3,5)-pyrazolo[1,5-a]-pyrimidina-2(1,3)-benzenacyclodecaphane-24-carboxylate (12b)**

To a solution of 10-(tert-butyl) 24-methyl (13Z,14E)-3-oxa-10-aza-1(3,5)- pyrazolo[1,5-a]pyrimidina-2(1,3)-benzenacyclodecaphane-24,10-dicarboxylate (**11b**) (0.48 g, 1.0 eq, 1.03 mmol) in dry methylene chloride (25 mL), trifluoroacetic acid (1.97 mL g, 25.0 eq, 25.72 mmol) was added dropwise and it was stirred for 16 h at room temperature. Afterwards, saturated K_2_CO_3_ solution was added to the stirring solution. Subsequently, the solution was filtrated, and the solvent was removed, and the residue was taken up in ethyl acetate. The organic phase was washed with saturated K_2_CO_3_ solution, and the aqueous phase was extracted with ethyl acetate. The combined organic phases were washed with brine and dried over MgSO4. After removal of the organic solvent under reduced pressure, the purification was done by flash column chromatography with a mobile phase of ethyl acetate (isocratic ratio). The title compound was yielded as a white solid (0.35 g, 93%).

1H NMR (500 MHz, DMSO-d6): δ 8.48 (d, J = 7.5 Hz, 1H, HetH), 8.42 (s, 1H, HetH), 8.35 (d, J = 1.5 Hz, 1H, PhH), 7.98 (t, J = 5.9 Hz, 1H, NH), 7.66 (d, J = 8.2 Hz, 1H, PhH), 7.34 (dd, J = 8.3, 1.4 Hz, 1H, PhH), 6.28 (d, J = 7.6 Hz, 1H, HetH), 4.25 – 4.16 (m, 2H, CH2), 3.76 (s, 3H, OCH3), 3.35 – 3.30 (m, 2H, CH2) 1.96 – 1.85 (m, 2H, CH2), 1.85 – 1.73 (m, 2H, CH2), 1.56 – 1.41 (m, 4H, CH2) (ppm).

13C NMR (126 MHz, DMSO-d6): δ 165.88, 158.31, 156.22, 145.38, 142.17, 139.16, 135.44, 131.47, 115.45, 115.32, 109.78, 103.14, 100.57, 68.48, 51.49, 24.03, 23.80, 23.11, 21.42 (ppm).

MS (ESI+) m/z: 366.40 [M + H]+.

HRMS m/z: [M + Na]+ calcd for C20H22N4O3Na, 389.15841; found 389.15814.

HPLC (I): tR = 15.499, purity ≥ 95%.

**Synthesis of (13Z,14E)-3,6-dioxa-9-aza-1(3,5)-pyrazolo[1,5-a]pyrimidina- 2(1,3)-benzenacyclononaphane-24-carboxylic acid (13a)**

(1^3^Z,1^4^E)-3,6-dioxa-9-aza-1(3,5)-pyrazolo[1,5-a]-pyrimidina-2(1,3)benzenacyclononaphane-2^4^-carboxylate (**12a**) (600 mg, 1.0 eq, 1.32 mmol) and Lithium hydroxide (1.43 g, 6.0 eq, 7.92 mmol) were solved in Methanol and water (50:50) (8 mL:8 mL) and the reaction mixture was stirred for 16 h 55 °C. Methanol was removed under reduced pressure and the aqueous suspension was acidified with 1 M HCl. The formed precipitate was filtered and washed with water and dried under reduced pressure. The title compound was obtained as a white solid (316 mg, 54%).

1H NMR (500 MHz, DMSO-d6): δ 12.24 (bs, 1H, COOH), 8.81 (s, 1H, HetH), 8.57 (d, J = 7.6 Hz, 1H, HetH), 8.38 (s, 1H, PhH), 7.93 (t, J = 5.4 Hz, 1H, NH), 7.70 (d, J = 8.1 Hz, PhH), 7.27 (dd, J = 8.1, 1.4 Hz, 1H, PhH), 6.34 (d, J = 7.6 Hz, 1H, HetH), 4.38 (t, J = 7.1 Hz, 2H, CH2), 3.99 (t, J = 7.5 Hz, 2H, CH2), 3.89 – 3.86 (m, 2H, CH2), 3.56 – 3.50 (m, 2H, CH2) ppm.

13C NMR (126 MHz, DMSO-d6): δ 166.70, 158.07, 156.14, 145.27, 141.76, 138.69, 135.80, 131.82, 115.79, 115.61, 110.39, 103.50, 100.18, 65.31, 65.22, 63.89, 59.77 ppm.

MS (ESI+) m/z: 340.93 [M + H]+.

HRMS m/z: [M + H]+ calcd for C17H17N4O4, 341.12443; found 341.12573.

HPLC (III): tR = 11.407, purity ≥ 95%.

**Synthesis of (13Z,14E)-3-oxa-10-aza-1(3,5)-pyrazolo[1,5-a]pyrimidina- 2(1,3)-benzena-cyclodecaphane-24-carboxylic acid (13b)**

Methyl(1^3^Z,1^4^E)-3-oxa-10-aza-1(3,5)-pyrazolo[1,5-a]-pyrimidina-2(1,3)benzenacyclo decaphane -2^4^-carboxylate (**12b**) (0.35 g, 1.0 eq, 0.96 mmol) and Lithium hydroxide (0.60 g, 15.0 eq, 14.3 mmol) were solved in Methanol and water (50:50) (20 mL) and the reaction mixture was stirred for 16 h 55 °C. Methanol was removed under reduced pressure and the aqueous suspension was acidified with 1 M HCl. The formed precipitate was filtered and washed with water and dried under reduced pressure. The title compound was obtained as a white solid (0.21 g, 63%).

1H NMR (500 MHz, DMSO-d6): δ 12.15 (s, 1H, COOH), 8.48 (d, J = 7.6 Hz, 1H, HetH), 8.42 (s, 1H, HetH), 8.34 (s, 1H, PhH), 7.98 (t, J = 5.8 Hz, 1H, NH), 7.67 (d, J = 8.2 Hz, 1H, PhH), 7.34 (d, J = 8.2 Hz, 1H, PhH), 6.28 (d, J = 7.6 Hz, 1H, HetH), 4.30 – 4.15 (m, 2H, CH2), 1.99 – 1.86 (m, 2H, CH2), 1.85 – 1.74 (m, 2H, CH2), 1.58 – 1.40 (m, 4H, CH2) (ppm).

13C NMR (126 MHz, DMSO-d6): δ 166.87, 158.24, 156.20, 145.33, 142.15, 138.89, 135.45, 131.72, 116.19, 115.53, 109.75, 103.22, 100.56, 68.53, 24.06, 23.84, 23.14, 21.46 (ppm).

MS (ESI+) m/z: 353.10 [M + H]+.

HRMS m/z: [M + Na]+ calcd for C19H20N4O3Na, 375.14276; found 375.14249.

HPLC (I): tR = 8.494, purity ≥ 95%.

**Synthesis of (13Z,14E)-N-(4-(hydroxymethyl)cyclohexyl)-3,6-dioxa-9-aza-1(3,5)-pyrazolo[1,5-a]pyrimidina-2(1,3)-benzenacyclononaphane-24-carboxamide (14)**

(1^3^Z,1^4^E)-3,6-dioxa-9-aza-1(3,5)-pyrazolo[1,5-a]pyrimidina- 2(1,3)-benzenacyclononaphane-24-carboxylic acid (0.03 g, 1.0 eq, 88.2 µmol), DIPEA (23.0 µL, 1.5 eq, 0.132 mmol) and 7-azabenzotriazol-1-yloxy)tripyrrolidinophosphonium hexafluorophosphate (0.083 g, 1.8 eq, 0.159 mmol) were solved in dry dimethylformamide and stirred for 10 min at room temperature. Afterwards, (2-chloropyridin-4-yl)methanamine (0.023 g, 1.8 eq, 0.159 mmol) was added and the solution was stirred for 16 h at ambient temperature. The solvent was removed under reduced pressure and to the residue was added acetonitrile. The formed precipitate was filtered and further washed with acetonitrile, to obtain the product as a white solid (40 mg, 95%)

1H NMR (250 MHz, DMSO) δ 8.84 (s, 1H), 8.77 (t, J = 6.2 Hz, 1H), 8.60 – 8.54 (m, 1H), 8.38 (s, 1H), 8.35 (d, J = 5.1 Hz, 1H), 7.92 (d, J = 5.2 Hz, 1H), 7.79 (d, J = 8.1 Hz, 1H), 7.42 (s, 1H), 7.33 (t, J = 6.4 Hz, 2H), 6.34 (d, J = 7.6 Hz, 1H), 4.54 (d, J = 6.0 Hz, 2H), 4.48 (s, 2H), 4.07 (d, J = 8.4 Hz, 2H), 3.97 – 3.83 (m, 2H), 3.56 (s, 2H), 2.07 (s, 2H) ppm.

MS (ESI+) m/z: 452.20 [M + H]+.

Precision mass m/z: [M + H]+ calcd for C24H30N5O4, 452.2292; found 452.2285

HPLC: tR = 3.534, purity ≥ 95%.

**Synthesis of (13Z,14E)-N-((2-chloropyridin-4-yl)methyl)-3,6-dioxa-9-aza-1(3,5)-pyrazolo[1,5-a]pyrimidina-2(1,3)-benzenacyclononaphane-24-carboxamide (15)**

(13Z,14E)-3,6-dioxa-9-aza-1(3,5)-pyrazolo[1,5-a]pyrimidina- 2(1,3)-benzenacyclononaphane-24-carboxylic acid acid (0.03 g, 1.0 eq, 88.2 µmol), DIPEA (23.0 µL, 1.5 eq, 0.132 mmol) and 7-azabenzotriazol-1-yloxy)tripyrrolidinophosphonium hexafluorophosphate (0.083 g, 1.8 eq, 0.159 mmol) were solved in dry dimethylformamide and stirred for 10 min at room temperature. Afterwards, (4-aminocyclohexyl)methanol (0.02 g, 1.8 eq, 0.155 mmol) was added and the solution was stirred for 16 h at ambient temperature. The solvent was removed under reduced pressure and to the residue was added acetonitrile. The formed precipitate was filtered and further washed with acetonitrile, to obtain the product as a white solid (28 mg, 68%)

1H NMR (250 MHz, DMSO) δ 8.80 (s, 1H), 8.55 (d, J = 7.6 Hz, 1H), 8.36 (s, 1H), 7.88 (d, J = 5.3 Hz, 1H), 7.83 (d, J = 7.8 Hz, 1H), 7.79 (d, J = 8.1 Hz, 1H), 7.30 (d, J = 8.1 Hz, 1H), 6.33 (d, J = 7.6 Hz, 1H), 4.52 – 4.36 (m, 3H), 4.09 – 3.97 (m, 2H), 3.96 – 3.82 (m, 2H), 3.78 – 3.61 (m, 1H), 3.55 (s, 2H), 3.23 (t, J = 5.7 Hz, 2H), 1.92 (d, J = 9.6 Hz, 2H), 1.77 (d, J = 11.7 Hz, 2H), 1.28 (dd, J = 24.2, 9.2 Hz, 3H), 0.99 (dd, J = 23.5, 11.1 Hz, 2H) ppm.

MS (ESI+) m/z: 465.10 [M + H]+.

Precision mass m/z: [M + H]+ calcd for C23H21ClN6O3, 465.1436; found 465.1425.

HPLC: tR = 3.820, purity ≥ 95%.

**Synthesis of (13Z,14E)-N-(3-chlorobenzyl)-3,6-dioxa-9-aza-1(3,5)-pyrazolo- [1,5-a]pyrimidina-2(1,3)-benzenacyclononaphane-24-carboxamide (16)**

(13Z,14E)-3,6-dioxa-9-aza-1(3,5)-pyrazolo[1,5-a]pyrimidina-2(1,3)-benzenacyclononaphane- 24-carboxylic acid (**13a**) (35 mg, 1.0 eq, 0.09 mmol) was dissolved in dry dimethylformamide (5 mL) and dry N,N-diisopropylethylamine (29 mg, 2.4 eq, 0.22 mmol) as well as (1-[bis(dimethylamino)methylene]-1H-1,2,3-triazolo[4,5-b]pyridinium 3-oxide hexafluoro-phosphate (41 mg, 1.2 eq, 0.11 mmol) were added. The mixture was stirred for 1 h at room temperature under argon atmosphere. Afterwards, (3-chlorophenyl)methanamine (23 mg, 1.2 eq, 0.16 mmol) was added and the mixture was stirred at room temperature for 18 h. After quenching with water and acidification with 1 N HCl, the precipitate was separated by vacuum filtration. The desired product was obtained as a white solid (64 mg, 94%).

1H NMR (500 MHz, DMSO-d6): δ 8.83 (d, J = 1.5 Hz, 1H, PhH), 8.71 (t, J = 6.2 Hz, 1H, NH), 8.57 (d, J = 7.6 Hz, 1H HetH), 8.38 (s, 1H, HetH), 7.91 (t, J = 5.4 Hz, 1H, NH), 7.80 (d, J = 8.1 Hz, 1H, PhH), 7.43 – 7.25 (m, 5H, PhH), 6.34 (d, J = 7.6 Hz, 1H, HetH), 4.54 – 4.43 (m, 4H, CH2), 4.10 – 4.00 (m, 2H, CH2), 3.95 – 3.84 (m, 2H, CH2), 3.61 – 3.48 (m, 2H, CH2) (ppm).

13C NMR (126 MHz, DMSO-d6): δ 164.89, 156.45, 156.10, 145.21, 142.76, 141.66, 137.67, 135.81, 132.92, 131.00, 130.15, 126.85, 126.50, 125.75, 117.96, 116.19, 110.00, 103.57,100.15, 65.41, 64.03, 42.10 (ppm).

MS (ESI+) m/z: 464.00 [M + H]+.

HRMS m/z: [M + H]+ calcd for C24H22ClN5O3, 464.14839; found 464.14759.

HPLC (I): tR = 14.783, purity ≥ 95%.

**Synthesis of (13Z,14E)-N-(4-chlorobenzyl)-3,6-dioxa-9-aza-1(3,5)-pyrazolo-[1,5-a]pyrimidina-2(1,3)-benzenacyclononaphane-24-carboxamide (17)**

The title compound was synthesized according to the procedure of **16** using (4-chlorophenyl)methanamine (23 mg, 1.2 eq, 0.16 mmol). The desired product was obtained as a white solid (66 mg, 97%).

1H NMR (500 MHz, DMSO-d6): δ 8.82 (d, J = 1.5 Hz, 1H, PhH), 8.69 (t, J = 6.2 Hz, 1H, NH), 8.56 (d, J = 7.6 Hz, 1H, HetH), 8.37 (s, 1H, HetH), 7.93 (t, J = 5.4 Hz, 1H, NH), 7.82 (d, J = 8.1 Hz, 1H, PhH), 7.41 – 7.28 (m, 5H, PhH), 6.34 (d, J = 7.6 Hz, 1H, HetH), 4.55 – 4.43 (m, 4H, CH2), 4.07 – 4.00 (m, 2H, CH2), 3.94 – 3.84 (m, 2H, CH2), 3.59 – 3.49 (m, 2H, CH2) (ppm).

13C NMR (126 MHz, DMSO-d6): δ 164.80, 156.54, 156.12, 145.23, 141.68, 139.15, 137.73, 135.80, 131.12, 131.10, 130.91, 128.97, 128.64, 128.21, 117.80, 116.20, 110.02, 103.58, 100.18, 65.46, 65.38, 64.04, 42.03 (ppm).

MS (ESI+) m/z: 463.97 [M + H]+.

HRMS m/z: [M + H]+ calcd for C24H22ClN5O3, 464.14839; found 464.14798.

HPLC (I): tR = 14.814, purity ≥ 95%.

**Synthesis of (13Z,14E)-N-benzyl-3,6-dioxa-9-aza-1(3,5)-pyrazolo[1,5-a]- pyrimidina-2(1,3)-benzenacyclononaphane-24-carboxamide (18)**

The title compound was synthesized according to the procedure of **16** using benzylamine (12 mg, 1.2 eq, 0.11 mmol). The desired product was obtained as a white solid (33 mg, 75%).

1H NMR (500 MHz, DMSO-d6): δ 8.82 (d, J = 1.5 Hz, 1H, PhH), 8.64 (t, J = 6.1 Hz, 1H, NH), 8.57 (d, J = 7.6 Hz, 1H, HetH), 8.38 (s, 1H, HetH), 7.91 (t, J = 5.4 Hz, 1H, NH), 7.83 (d, J = 8.1 Hz, 1H, PhH), 7.33 (d, J = 4.5 Hz, 5H, PhH), 7.27 – 7.19 (m, 1H, PhH), 6.34 (d, J = 7.6 Hz, 1H, HetH), 4.52 (d, J = 6.1 Hz, 2H, CH2), 4.50 – 4.43 (m, 2H, CH2), 4.04 (dd, J = 9.1, 6.3 Hz, 2H, CH2), 3.89 (dd, J = 9.2, 6.0 Hz, 2H, CH2), 3.54 (q, J = 7.2, 6.8 Hz, 2H, CH2) (ppm).

13C NMR (126 MHz, DMSO-d6): δ 164.69, 156.50, 156.10, 145.20, 141.66, 140.00, 137.64, 135.80, 131.11, 128.26, 127.02, 126.58, 117.92, 116.20, 110.02, 103.57, 100.15, 65.46, 65.36, 64.03 (ppm).

MS (ESI+) m/z: 429.95 [M + H]+.

HRMS m/z: [M + H]+ calcd for C24H23N5O3, 430.18737; found 430.18729.

HPLC (I): tR = 13.843, purity ≥ 95%.

**Synthesis of (13Z,14E)-N-(3-chlorobenzyl)-3-oxa-10-aza-1(3,5)-pyrazolo-[1,5-a]pyrimidina-2(1,3)-benzenacyclodecaphane-24-carboxamide (19)**

The title compound was synthesized according to the procedure of **16** using (3 chlorophenyl)methanamine (21 mg, 1.1 eq, 0.15 mmol). The desired product was obtained as a white solid (52 mg, 77%).

1H NMR (500 MHz, DMSO-d6): δ 8.72 (t, J = 6.2 Hz, 1H, NH), 8.48 (d, J = 7.6 Hz, 1H, HetH), 8.42 (s, 1H, HetH), 8.35 (d, J = 1.5 Hz, 1H, PhH), 7.96 (t, J = 6.0 Hz, 1H, NH), 7.78 (d, J = 8.2 Hz, 1H, PhH), 7.42 – 7.34 (m, 3H, PhH), 7.32 – 7.28 (m, 2H, PhH), 6.27 (d, J = 7.6 Hz, 1H, HetH), 4.51 (d, J = 6.1 Hz, 2H, CH2), 4.38 – 4.31 (m, 2H, CH2), 3.37 – 3.33 (m, 2H, CH2), 2.05 – 1.96 (m, 2H, CH2), 1.86 – 1.77 (m, 2H, CH2), 1.55 – 1.43 (m, 4H, CH2) (ppm).

13C NMR (126 MHz, DMSO-d6): δ 165.15, 156.58, 156.15, 145.22, 142.81, 142.06, 137.90, 135.42, 132.92, 131.01, 130.11, 126.84, 126.47, 125.75, 118.11, 115.86, 108.55, 103.31, 100.52, 68.20, 42.08, 40.17, 23.91, 23.59, 22.95, 21.24 (ppm).

MS (ESI+) m/z: 478.90 [M + H]+.

HRMS m/z: [M + H]+ calcd for C26H27 ClN5O2, 476.18478; found 476.18416.

HPLC (I): tR = 16.725, purity ≥ 95%.

**Synthesis of (13Z,14E)-N-(2,3-dichlorobenzyl)-3,6-dioxa-9-aza-1(3,5)-pyrazolo[1,5-a]pyrimidina-2(1,3)-benzenacyclononaphane-24-carboxamide (20)**

The title compound was synthesized according to the procedure of **16** using (2,3 dichlorophenyl)methanamine (25 mg, 1.2 eq, 0.14 mmol). The desired product was obtained as a white solid (46 mg, 79%).

1H NMR (500 MHz, DMSO-d6): δ 8.85 (d, J = 1.5 Hz, 1H, PhH), 8.77 (t, J = 6.1 Hz, 1H, NH), 8.57 (d, J = 7.6 Hz, 1H, HetH), 8.38 (s, 1H, HetH), 7.91 (t, J = 5.4 Hz, 1H, NH), 7.83 (d, J = 8.1 Hz, 1H, PhH), 7.55 (dd, J = 7.8, 1.8 Hz, 1H, PhH), 7.40 – 7.30 (m, 3H, PhH), 6.34 (d, J = 7.6 Hz, 1H, HetH), 4.60 (d, J = 6.1 Hz, 2H, CH2), 4.53 – 4.48 (m, 2H, CH2), 4.10 – 4.01 (m, 2H, CH2), 3.93 – 3.86 (m, 2H, CH2), 3.58 – 3.51 (m, 2H, CH2) (ppm).

13C NMR (126 MHz, DMSO-d6): δ 164.84, 156.63, 156.10, 145.22, 141.65, 139.56, 137.91, 135.77, 131.57, 131.08, 129.66, 128.75, 128.09, 126.94, 117.44, 116.24, 110.09, 103.52, 100.15, 65.61, 65.44, 64.17, 41.39 (ppm).

MS (ESI+) m/z: 498.10 [M + H]+.

HRMS m/z: [M + H]+ calcd for C24H22Cl2N5O3, 498.10942; found 498.10862.

HPLC (II): tR = 9.533, purity ≥ 95%.

**Synthesis of (13Z,14E)-N-(3-chloro-5-fluorobenzyl)-3,6-dioxa-9-aza-1(3,5)- pyrazolo[1,5-a]pyrimidina-2(1,3)-benzenacyclononaphane-24-carboxamide (21)**

The title compound was synthesized according to the procedure of **16** using (3-chloro-5- fluorophenyl)methanamine (23 mg, 1.2 eq, 0.14 mmol). The desired product was obtained as a white solid (30 mg, 53%).

1H NMR (500 MHz, DMSO-d6): δ 8.83 (d, J = 1.5 Hz, 1H, PhH), 8.74 (t, J = 6.2 Hz, 1H, NH), 8.55 (d, J = 7.5 Hz, 1H, HetH), 8.37 (s, 1H, HetH), 7.90 (t, J = 5.4 Hz, 1H, NH), 7.78 (d, J = 8.1 Hz, 1H, PhH), 7.32 (dd, J = 8.2, 1.4 Hz, 1H, PhH), 7.28 (dt, J = 8.6, 2.2 Hz, 1H, PhH), 7.26 – 7.25 (m, 1H, PhH), 7.18 – 7.13 (m, 1H, PhH), 6.34 (d, J = 7.6 Hz, 1H, HetH), 4.52 – 4.47 (m, 4H, CH2), 4.09 – 4.00 (m, 2H, CH2), 3.93 – 3.84 (m, 2H, CH2), 3.56 – 3.51 (m, 2H, CH2) (ppm).

13C NMR (126 MHz, DMSO-d6): δ 165.55, 163.59, 161.63, 156.94, 156.57, 145.68, 145.47, 145.40, 142.09, 138.22, 136.22, 134.20, 134.11, 131.43, 123.65, 123.63, 118.32, 116.67, 114.63, 114.43, 113.39, 113.21, 110.50, 104.02, 100.63, 65.95, 65.90, 64.59, 42.42 (ppm).

MS (ESI+) m/z: 482.20 [M + H]+.

HRMS m/z: [M + H]+ calcd for C24H22ClFN5O3, 482.13897; found 482.13835.

HPLC (I): tR = 14.031, purity ≥ 95%.

**Synthesis of (13Z,14E)-N-(3-cyanobenzyl)-3,6-dioxa-9-aza-1(3,5)-pyrazolo-[1,5-a]pyrimidina-2(1,3)-benzenacyclononaphane-24-carboxamide (22)**

The title compound was synthesized according to the procedure of **16** using (3-cyanophenyl)methanamine (18 mg, 1.2 eq, 0.14 mmol). The desired product was obtained as a white solid (38 mg, 71%)

1H NMR (500 MHz, DMSO-d6): δ 8.83 (d, J = 1.5 Hz, 1H, PhH), 8.75 (t, J = 6.2 Hz, 1H, NH), 8.55 (d, J = 7.6 Hz, 1H, HetH), 8.37 (s, 1H, HetH), 7.90 (t, J = 5.4 Hz, 1H, NH), 7.80 (d, J = 8.1 Hz, 1H, PhH), 7.77 – 7.75 (m, 1H, PhH), 7.69 (td, J = 16.0, 8.0, 1.5 Hz, 2H, PhH), 7.55 (t, J = 7.7 Hz, 1H, NH), 7.32 (dd, J = 8.1, 1.4 Hz, 1H, PhH), 6.34 (d, J = 7.6 Hz, 1H, HetH), 4.55 (d, J = 6.2 Hz, 2H, CH2), 4.52 – 4.47 (m, 2H, CH2), 4.08 – 4.01 (m, 2H, CH2), 3.91 – 3.85 (m, 2H, CH2), 3.56 – 3.51 (m, 2H, CH2) (ppm).

13C NMR (126 MHz, DMSO-d6): δ 165.07, 156.54, 156.13, 145.24, 141.87, 141.66, 137.76, 135.79, 132.11, 131.02, 130.61, 130.43, 129.56, 118.96, 117.88, 116.22, 111.17, 111.16, 110.05, 103.59, 100.19, 65.53, 65.45, 64.14, 42.08 (ppm).

MS (ESI+) m/z: 455.20 [M + H]+.

HRMS m/z: [M + H]+ calcd for C25H23N6O3, 455.18262; found 455.18168.

HPLC (I): tR = 12.938, purity ≥ 95%.

**Synthesis of (13Z,14E)-N-(3-chloro-2-fluorobenzyl)-3,6-dioxa-9-aza-1(3,5)-pyrazolo[1,5-a]pyrimidina-2(1,3)-benzenacyclononaphane-24-carboxamide (23)**

The title compound was synthesized according to the procedure of **16** using (3-chloro-2- fluorophenyl)methanamine (23 mg, 1.2 eq, 0.14 mmol). The desired product was obtained as a slightly yellow solid (38 mg, 67%).

1H NMR (500 MHz, DMSO-d6): δ 8.83 (d, J = 1.5 Hz, 1H, PhH), 8.71 (t, J = 6.0 Hz, 1H, NH), 8.55 (d, J = 7.5 Hz, 1H, HetH), 8.37 (s, 1H, HetH), 7.90 (t, J = 4.9 Hz, 1H, NH), 7.81 (d, J = 8.1 Hz, 1H, PhH), 7.46 (td, J = 7.6, 1.7 Hz, 1H, PhH), 7.36 – 7.29 (m, 2H, PhH), 7.23 – 7.17 (m, 1H, PhH), 6.34 (d, J = 7.6 Hz, 1H, HetH), 4.57 (d, J = 6.0 Hz, 2H), 4.52 – 4.46 (m, 2H, CH2), 4.07 – 4.01 (m, 2H, CH2), 3.90 – 3.87 (m, 2H, CH2), 3.54 – 3.52 (m, 2H, CH2) (ppm).

13C NMR (126 MHz, DMSO-d6): δ 165.38, 157.07, 156.58, 145.69, 142.10, 138.33, 136.22, 131.55, 129.33, 129.27, 129.21, 128.38, 128.35, 125.70, 125.65, 125.62, 117.98, 116.69, 110.55, 104.01, 100.64, 66.05, 64.61, 37.05 (ppm).

MS (ESI+) m/z: 482.20 [M + H]+.

HRMS m/z: [M + H]+ calcd for C24H22ClFN5O3, 482.13897; found 482.13837.

HPLC (I): tR = 13.927, purity ≥ 95%.

**Synthesis of (13Z,14E)-N-(pyridin-4-ylmethyl)-3,6-dioxa-9-aza-1(3,5)-pyrazolo[1,5-a]pyrimidina-2(1,3)-benzenacyclononaphane-24-carboxamide (24)**

The title compound was synthesized according to the procedure of **16** using pyridin-4- ylmethanamine (15 mg, 1.2 eq, 0.14 mmol). The desired product was obtained as a white solid (42 mg, 83%).

1H NMR (500 MHz, DMSO-d6): δ 8.84 (d, J = 1.5 Hz, 1H, PhH), 8.74 (t, J = 6.1 Hz, 1H, NH), 8.56 (d, J = 7.6 Hz, 1H, HetH), 8.53 – 8.47 (m, 2H, HetH), 8.38 (s, 1H, HetH), 7.91 (t, J = 5.4 Hz, 1H, NH), 7.82 (d, J = 8.1 Hz, 1H, PhH), 7.35 – 7.28 (m, 3H, HetH, PhH), 6.34 (d, J = 7.6 Hz, 1H, HetH), 4.57 – 4.46 (m, 4H, CH2), 4.09 – 4.02 (m, 2H, CH2), 3.93 – 3.86 (m, 2H, CH2), 3.59 – 3.50 (m, 2H, CH2) (ppm).

13C NMR (126 MHz, DMSO-d6): δ 164.96, 156.57, 156.09, 149.43, 149.07, 145.20, 141.63, 137.78, 135.76, 131.08, 122.00, 117.64, 116.17, 110.02, 103.54, 100.14, 65.48, 65.40, 64.08, 41.83 (ppm).

MS (ESI+) m/z: 431.20 [M + H]+.

HRMS m/z: [M + H]+ calcd for C23H23N6O3, 431.18262; found 431.18294.

HPLC (I): tR = 11.404, purity ≥ 95%.

**Synthesis of (13Z,14E)-N-(3-bromobenzyl)-3,6-dioxa-9-aza-1(3,5)-pyrazolo-[1,5-a]pyrimidina-2(1,3)-benzenacyclononaphane-24-carboxamide (25)**

The title compound was synthesized according to the procedure of **16** using (3-chlorophenyl)methanamine (26 mg, 1.2 eq, 0.14 mmol). The desired product wasobtained as a white solid (48 mg, 80%).

1H NMR (500 MHz, DMSO-d6): δ 8.83 (d, J = 1.5 Hz, 1H, PhH), 8.70 (t, J = 6.2 Hz, 1H, NH), 8.56 (d, J = 7.6 Hz, 1H, HetH), 8.37 (s, 1H, HetH), 7.90 (t, J = 5.4 Hz, 1H, NH), 7.79 (d, J = 8.1 Hz, 1H, PhH), 7.53 (t, J = 1.8 Hz, 1H, PhH), 7.43 (dt, J = 7.6, 1.7 Hz, 1H, PhH), 7.35 – 7.27 (m, 3H, PhH), 6.34 (d, J = 7.6 Hz, 1H, HetH), 4.52 – 4.48 (m, 4H, CH2), 4.09 – 4.01 (m, 2H, CH2), 3.93 – 3.84 (m, 2H, CH2), 3.56 – 3.51 (m, 2H, CH2) (ppm).

13C NMR (126 MHz, DMSO-d6): δ 164.97, 156.48, 156.13, 145.24, 143.03, 141.67, 137.69, 135.80, 131.01, 130.48, 129.76, 129.42, 126.19, 121.62, 118.03, 116.23, 110.06, 103.60, 100.19, 65.51, 65.45, 64.15, 42.09, 35.82 (ppm).

MS (ESI+) m/z: 510.20 [M + H]+.

HRMS m/z: [M + H]+ calcd for C24H23BrN5O3, 508.09788; found 508.09719.

HPLC (I): tR = 13.925, purity ≥ 95%.

**Synthesis of (13Z,14E)-N-(pyridin-3-ylmethyl)-3,6-dioxa-9-aza-1(3,5)-pyrazolo[1,5-a]pyrimidina-2(1,3)-benzenacyclononaphane-24-carboxamide (26)**

The title compound was synthesized according to the procedure of **16** using pyridin-3- ylmethanamine (15 mg, 1.2 eq, 0.14 mmol). The desired product was obtained as a white solid (40 mg, 79%).

1H NMR (500 MHz, DMSO-d6): δ 8.82 (d, J = 1.5 Hz, 1H, PhH), 8.73 (t, J = 6.1 Hz, 1H, NH), 8.56 – 8.54 (m, 2H, HetH), 8.47 – 8.42 (m, 1H, HetH), 8.37 (s, 1H, HetH), 7.90 (t, J = 5.4 Hz, 1H, NH), 7.82 (d, J = 8.1 Hz, 1H, PhH), 7.73 (dt, J = 7.9, 2.0 Hz, 1H, HetH), 7.36 (dd, J = 7.9, 4.8 Hz, 1H, PhH), 7.31 (dd, J = 8.1, 1.4 Hz, 1H, HetH), 6.33 (d, J = 7.6 Hz, 1H, HetH), 4.53 (d, J = 6.1 Hz, 2H, CH2), 4.50 – 4.44 (m, 2H, CH2), 4.08 – 3.99 (m, 2H, CH2), 3.92 – 3.83 (m, 2H, CH2), 3.53 (q, J = 6.7 Hz, 2H, CH2) (ppm).

13C NMR (126 MHz, DMSO-d6): δ 164.91, 156.56, 156.11, 148.70, 147.85, 145.23, 141.66, 137.77, 135.78, 135.57, 135.02, 131.10, 123.50, 117.72, 116.20, 110.04, 103.57, 100.17, 65.51, 65.44, 64.11, 40.44 (ppm).

MS (ESI+) m/z: 431.20 [M + H]+.

HRMS m/z: [M + H]+ calcd for C23H23N6O3, 431.18262; found 431.18114.

HPLC (I): tR = 11.177, purity ≥ 95%.

**Synthesis of (13Z,14E)-N-(2,4-dichlorobenzyl)-3,6-dioxa-9-aza-1(3,5)-pyrazolo[1,5-a]pyrimidina-2(1,3)-benzenacyclononaphane-24-carboxamide (27)**

The title compound was synthesized according to the procedure of **16** using 2,4-dichlorobenzylamine (28 mg, 1.2 eq, 0.16 mmol). The desired product was obtained as a white solid (53 mg, 72%).

1H NMR (500 MHz, DMSO-d6): δ 8.84 (d, J = 1.4 Hz, 1H, PhH), 8.74 (t, J = 6.1 Hz, 1H, NH), 8.57 (d, J = 7.6 Hz, 1H, HetH), 8.39 (s, 1H, HetH), 7.92 (t, J = 5.4 Hz, 1H, NH), 7.82 (d, J = 8.1 Hz, 1H, PhH), 7.63 (d, J = 2.2 Hz, 1H, PhH), 7.44 (dd, J = 8.4, 2.2 Hz, 1H, PhH), 7.36 (d, J = 8.4 Hz, 1H, PhH), 7.33 (dd, J = 8.2, 1.4 Hz, 1H, PhH), 6.34 (d, J = 7.6 Hz, 1H, HetH), 4.54 (d, J = 6.1 Hz, 2H, CH2), 4.52 – 4.47 (m, 2H, CH2), 4.12 – 3.99 (m, 2H, CH2), 3.94 – 3.84 (m, 2H, CH2), 3.61 – 3.48 (m, 2H, CH2) (ppm).

13C NMR (126 MHz, DMSO-d6): δ 164.86, 156.63, 156.13, 145.25, 141.70, 137.92, 136.07, 135.82, 132.68, 131.95, 131.11, 129.84, 128.51, 127.38, 117.46, 116.26, 110.08, 103.55, 100.17, 65.55, 65.40, 64.09 (ppm).

MS (ESI+) m/z: 598.01 [M + H]+.

HRMS m/z: [M + H]+ calcd for C24H23Cl2N5O3, 499.11278; found 499.11153.

HPLC (I): tR = 16.058, purity ≥ 95%.

**Synthesis of (13Z,14E)-N-(naphthalen-1-ylmethyl)-3,6-dioxa-9-aza-1(3,5)-pyrazolo[1,5-a]pyrimidina-2(1,3)-benzenacyclononaphane-24-carboxamide (28)**

The title compound was synthesized according to the procedure of **16** using naphthalen-1- ylmethanamine (19 mg, 1.2 eq, 0.12 mmol). The desired product was obtained as a white solid (31 mg, 63%).

1H NMR (500 MHz, DMSO-d6): δ 8.82 (d, J = 1.5 Hz, 1H, PhH), 8.68 (t, J = 6.0 Hz, 1H, NH), 8.56 (d, J = 7.6 Hz, 1H, HetH), 8.37 (s, 1H, HetH), 8.22 (dd, J = 8.4, 1.1 Hz, 1H, PhH), 7.99 – 7.94 (m, 1H, PhH), 7.91 (t, J = 5.4 Hz, 1H, NH), 7.86 – 7.81 (m, 2H, PhH), 7.62 – 7.53 (m, 2H, PhH), 7.52 – 7.46 (m, 2H, PhH), 7.33 (dd, J = 8.1, 1.4 Hz, 1H, PhH), 6.34 (d, J = 7.6 Hz, 1H, HetH), 4.98 (d, J = 5.9 Hz, 2H, CH2), 4.50 – 4.36 (m, 2H, CH2), 4.07 – 3.96 (m, 2H, CH2), 3.93 – 3.81 (m, 2H, CH2), 3.62 – 3.45 (m, 2H, CH2) (ppm).

13C NMR (126 MHz, DMSO-d6): δ 164.64, 156.46, 156.07, 145.18, 141.61, 137.62, 135.75, 134.94, 133.29, 131.00, 130.75, 128.54, 127.27, 126.15, 125.74, 125.48, 124.77, 123.40, 118.03, 116.21, 110.10, 103.54, 100.12, 65.57, 65.41, 64.12, 40.68 (ppm).

MS (ESI+) m/z: 480.15 [M + H]+.

HRMS m/z: [M + H]+ calcd for C28H26N5O3, 480.20302; found 480.20349.

HPLC (II): tR = 9.264, purity ≥ 95%.

**Synthesis of (13Z,14E)-N-(quinolin-4-ylmethyl)-3,6-dioxa-9-aza-1(3,5)-pyrazolo[1,5-a]pyrimidina-2(1,3)-benzenacyclononaphane-24-carboxamide (29)**

The title compound was synthesized according to the procedure of **16** using quinolin-4- ylmethanamine (21 mg, 1.2 eq, 0.11 mmol). The desired product was obtained as a slightly yellow solid (30 mg, 71%).

1H NMR (500 MHz, DMSO-d6): δ 8.89 (d, J = 4.5 Hz, 1H, HetH), 8.86 – 8.81 (m, 2H, NH, PhH), 8.57 (d, J = 7.5 Hz, 1H, HetH), 8.39 (s, 1H, HetH), 8.31 (dd, J = 8.4, 1.4 Hz, 1H, HetH), 8.08 (dd, J = 8.4, 1.3 Hz, 1H, HetH), 7.91 (t, J = 5.4 Hz, 1H, NH), 7.85 – 7.81 (m, 2H, HetH, PhH), 7.74 – 7.69 (m, 1H, HetH), 7.47 (d, J = 4.6 Hz, 1H, HetH), 7.34 (dd, J = 8.1, 1.4 Hz, 1H, PhH), 6.35 (d, J = 7.6 Hz, 1H, HetH), 5.05 (d, J = 5.9 Hz, 2H, CH2), 4.56 – 4.46 (m, 2H, CH2), 4.09 – 4.03 (m, 2H, CH2), 3.93 – 3.87 (m, 2H, CH2), 3.59 – 3.52 (m, 2H, CH2) (ppm).

13C NMR (126 MHz, DMSO-d6): δ 165.03, 156.57, 156.10, 145.22, 141.64, 137.81, 135.78, 131.04, 126.84, 125.99, 123.69, 118.84, 117.75, 116.21, 110.08, 103.55, 100.15, 65.53, 65.43, 64.12 (ppm).

MS (ESI+) m/z: 481.15 [M + H]+.

HRMS m/z: [M + Na]+ calcd for C27H24N6NaO3, 503.18021; found 503.17904.

HPLC (II): tR = 7.067, purity ≥ 95%.

**Synthesis of (13Z,14E)-N-((S)-1-phenylethyl)-3,6-dioxa-9-aza-1(3,5)-pyrazolo[1,5-a]pyrimidina-2(1,3)-benzenacyclononaphane-24-carboxamide (30)**

The title compound was synthesized according to the procedure of **16** using (S)-1-phenylethanamine (13 mg, 1.2 eq, 0.11 mmol). Purification was done by flash silica gel column chromatography with a mobile phase of ethyl acetate and ethanol with the addition of 1% acetic acid (ratio gradually ranging from 1:0 to 1:1). The obtained slightly yellow solid (32 mg, 82%) was the title compound.

1H NMR (500 MHz, DMSO-d6): δ 8.84 (s, 1H, PhH), 8.56 (d, J = 7.5 Hz, 1H, HetH), 8.41 – 8.34 (m, 2H, HetH, NH), 7.91 (t, J = 5.4 Hz, 1H, NH), 7.75 (d, J = 8.0 Hz, 1H, PhH), 7.41 – 7.30 (m, 5H, PhH), 7.26 – 7.21 (m, 1H), 6.34 (d, J = 7.6 Hz, 1H, HetH), 5.14 (p, J = 7.1 Hz, 1H, CH), 4.56 – 4.42 (m, 2H, CH2), 4.05 (t, J = 7.5 Hz, 2H, CH2), 3.95 – 3.85 (m, 2H, CH2), 3.59 – 3.51 (m, 2H, CH2), 1.47 (d, J = 7.0 Hz, 3H, CH3) (ppm).

13C NMR (126 MHz, DMSO-d6): δ 164.31, 156.89, 156.55, 145.66, 145.26, 142.09, 137.99, 136.24, 131.27, 128.82, 127.11, 126.40, 118.81, 116.74, 110.72, 104.02, 100.61, 66.28, 65.96, 64.72, 48.86, 23.28 (ppm).

MS (ESI+) m/z: 444.10 [M + H]+.

HRMS m/z: [M + Na]+ calcd for C25H25N5NaO3, 466.18496; found 466.18413.

HPLC (II): tR = 8.938, purity ≥ 95%.

Analytical data of compounds **2-30**

**Compound 2**

**^1^H NMR**

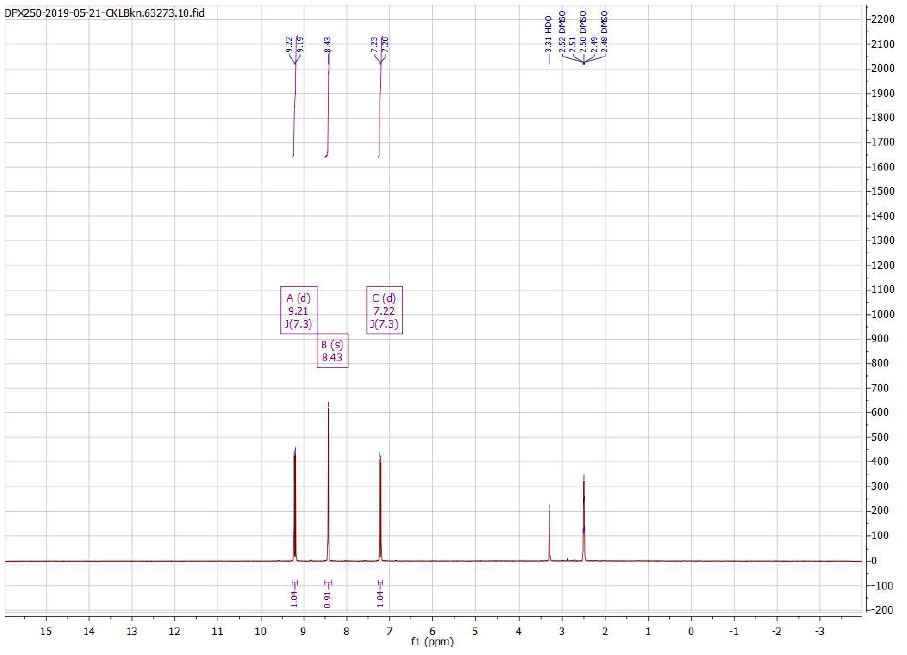

**Compound 3a**

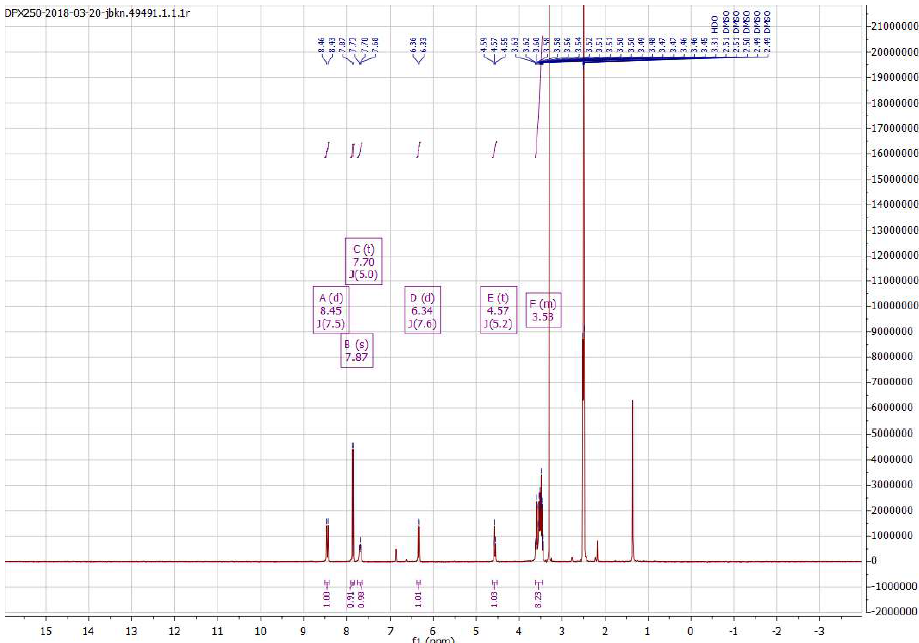
**^1^H NMR**

**Compound 3b**

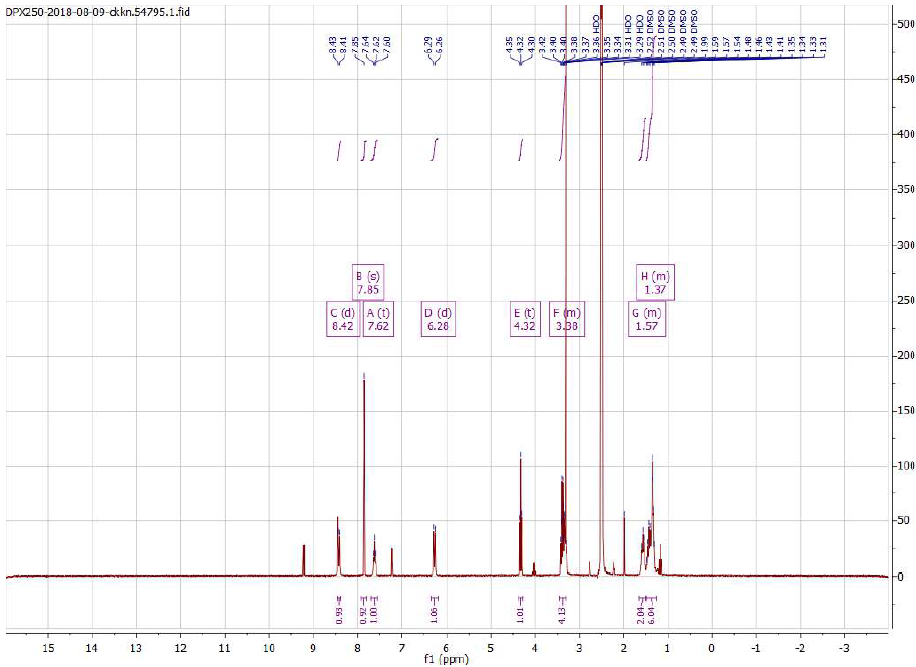
**^1^H NMR**

**Compound 4a**

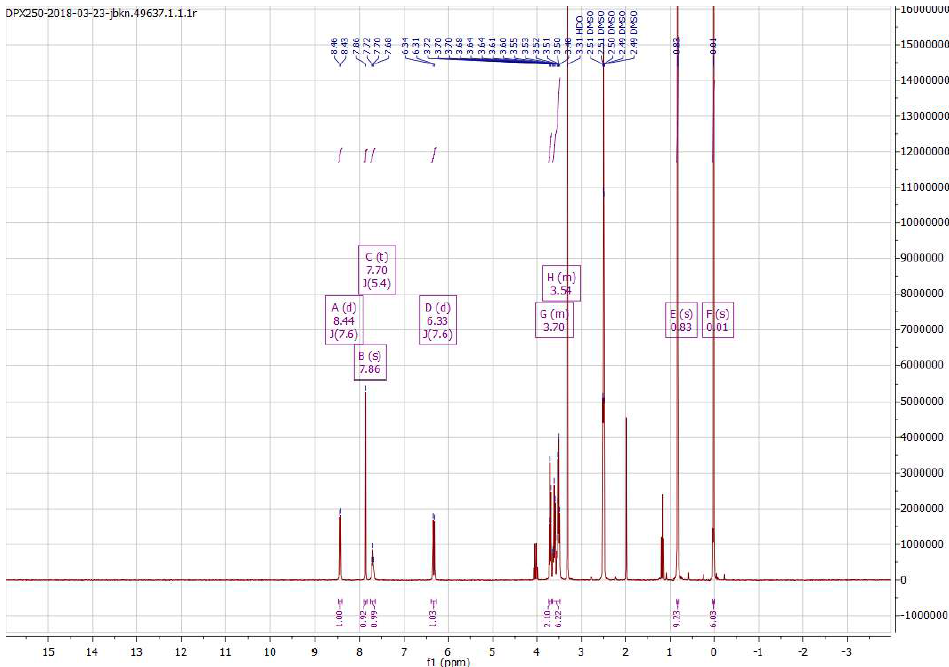
**^1^H NMR**

**Compound 4b**

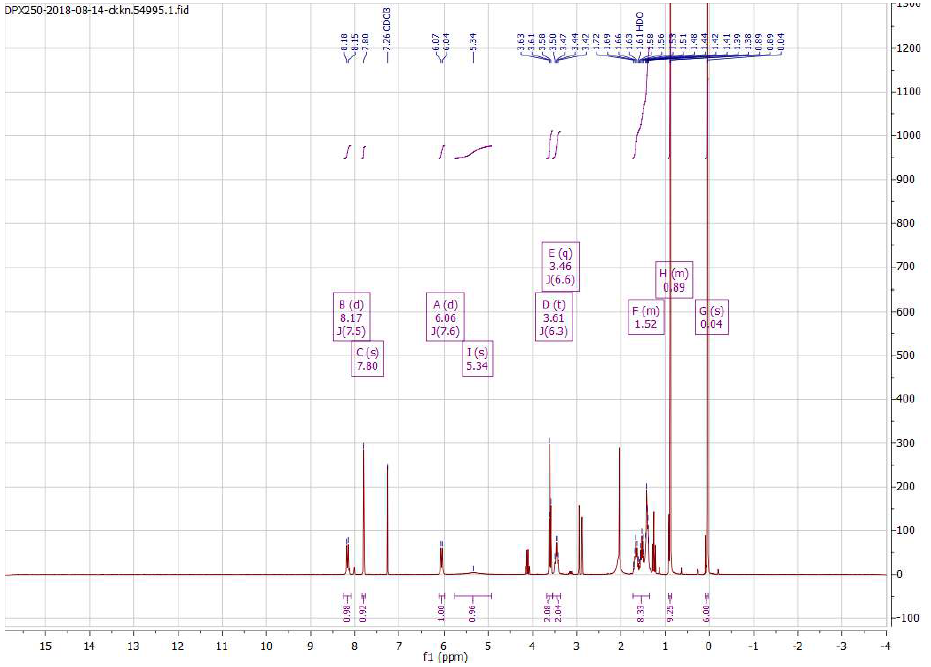
**^1^H NMR**

**Compound 5a**

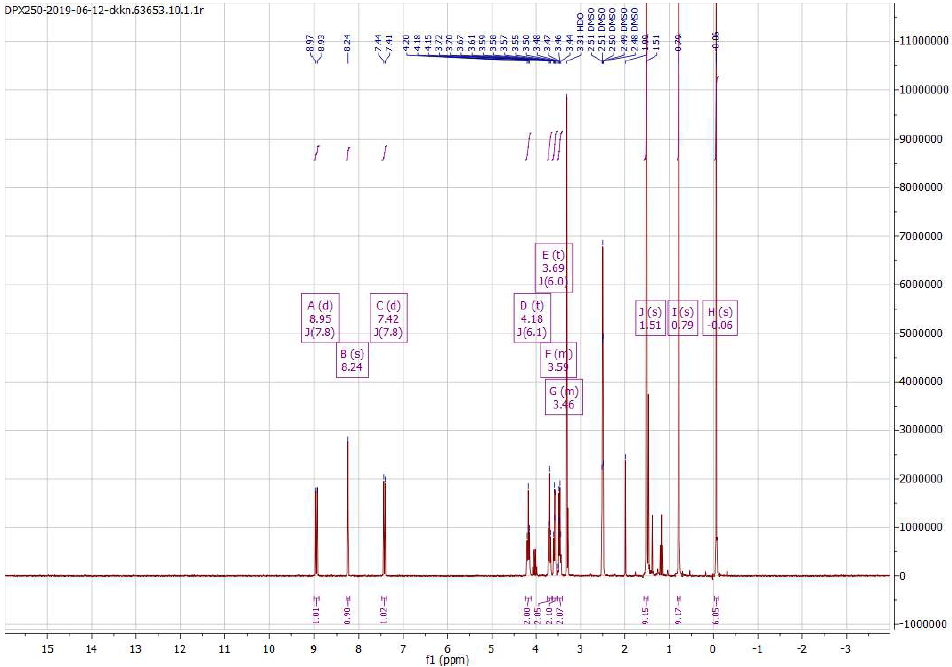
**^1^H NMR**

**Compound 5b**

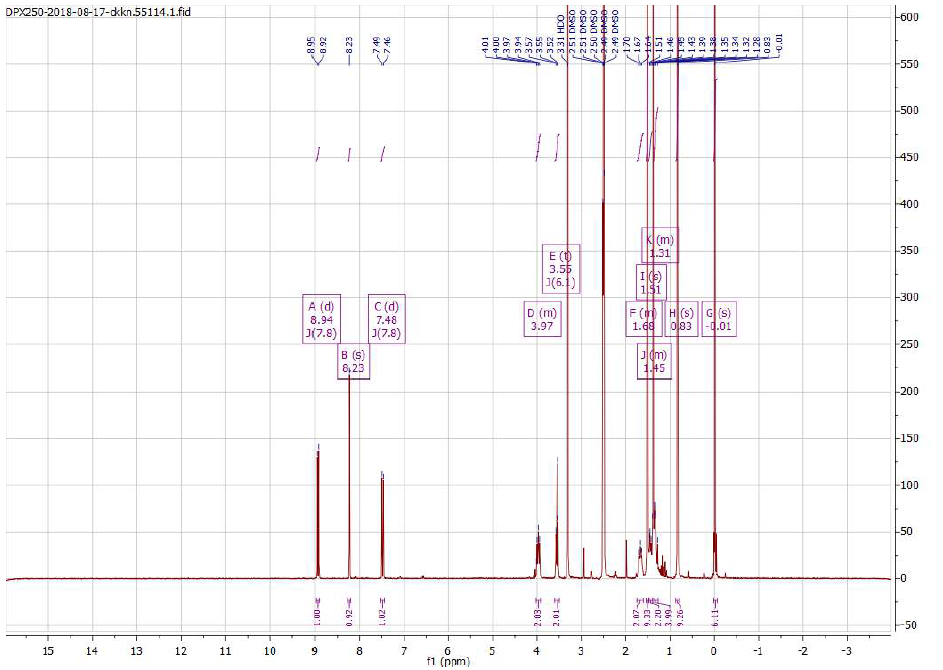
**^1^H NMR**

**Compound 7**

**^1^H NMR**

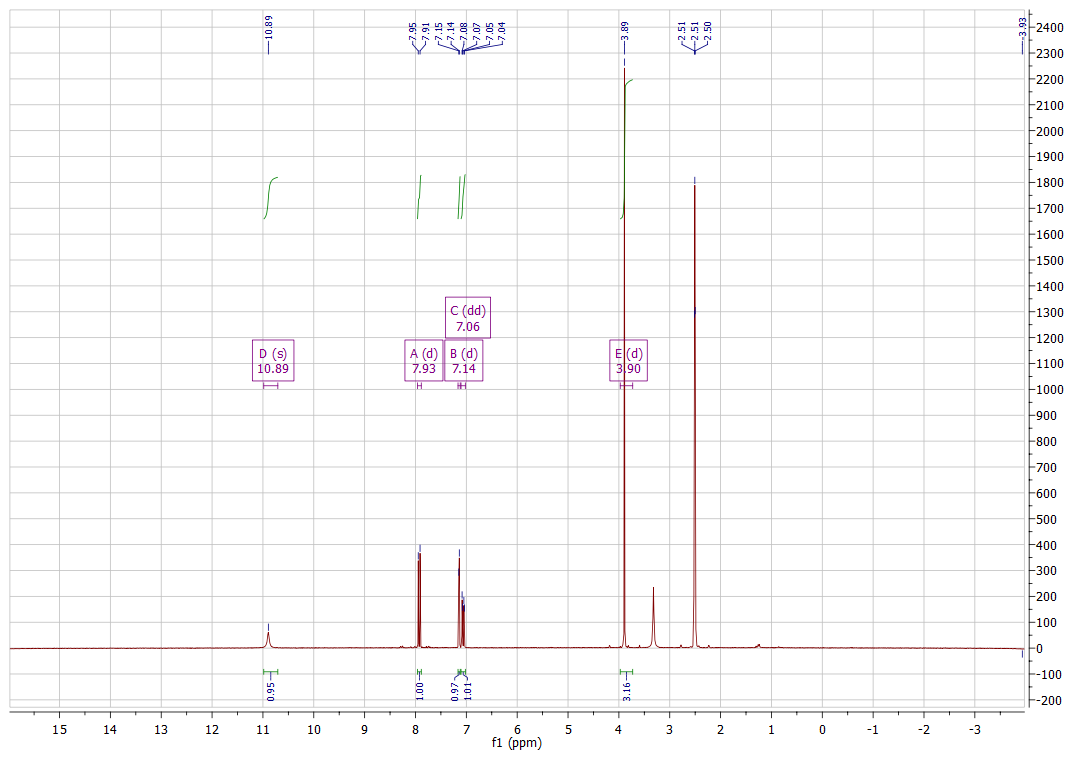

**Compound 8**

**^1^H NMR**

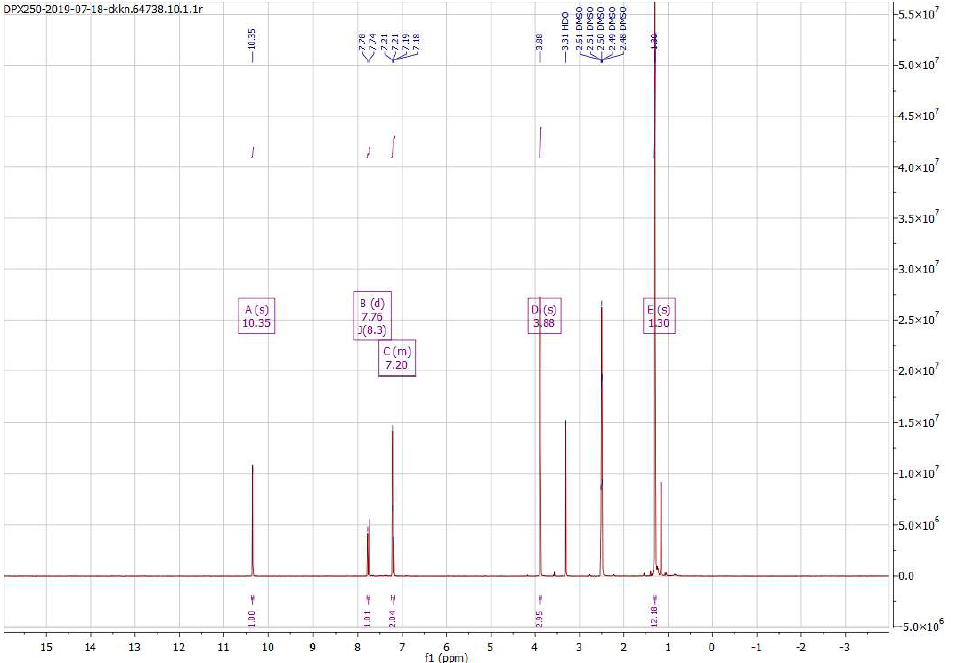

**Compound 9a**

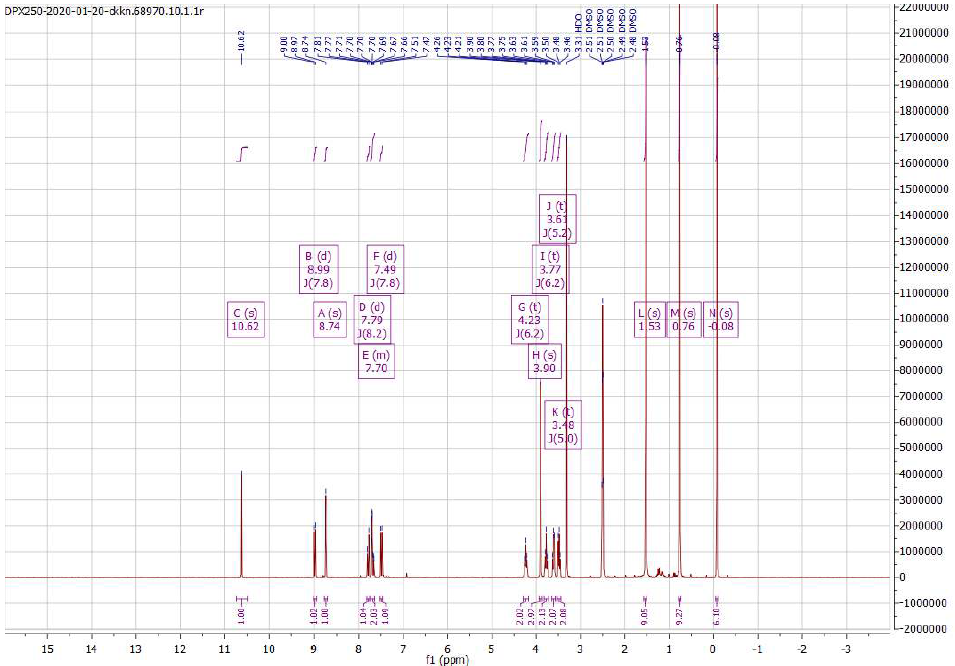
**^1^H NMR**

**Compound 9b**

**^1^H NMR**

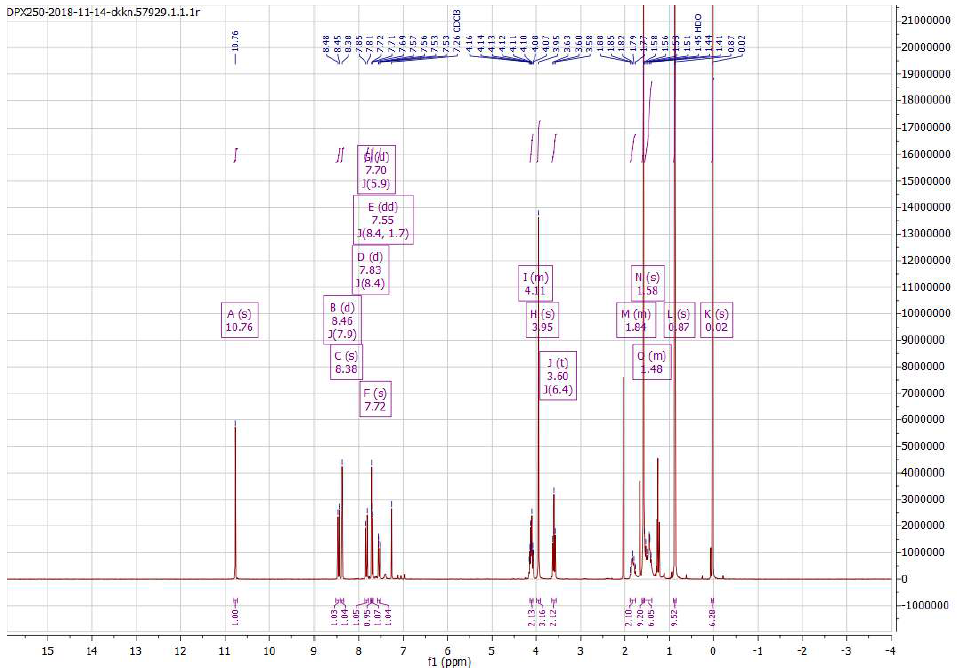

**Compound 10a**

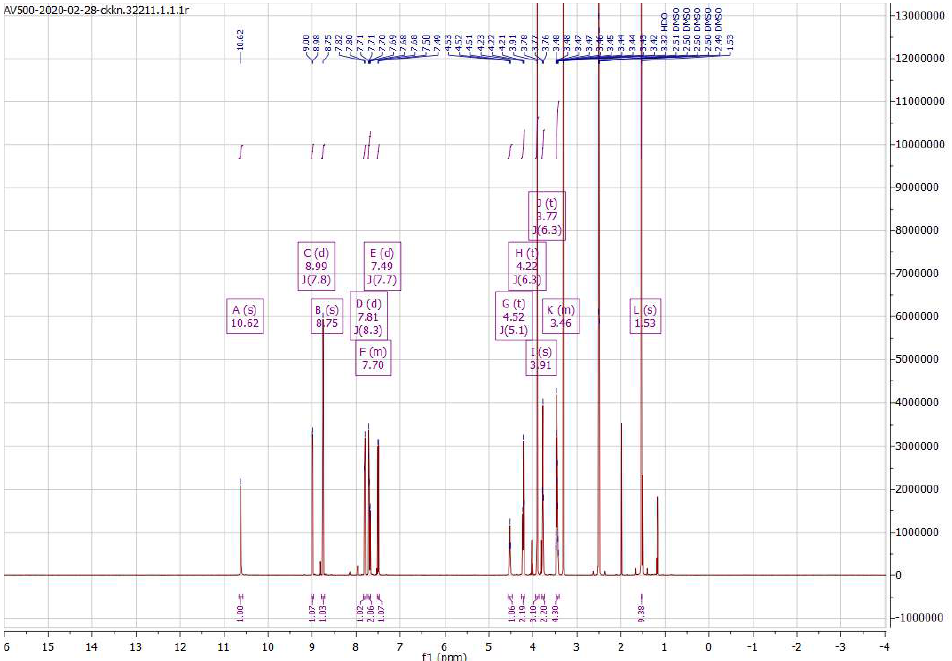
**^1^H NMR**

**Compound 10b**

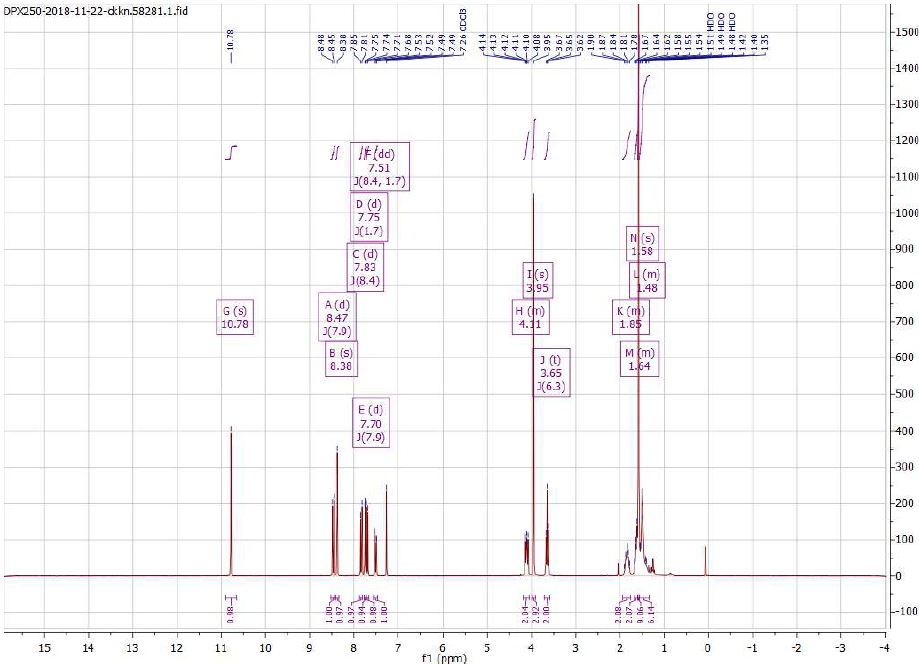
**^1^H NMR**

**Compound 11a**

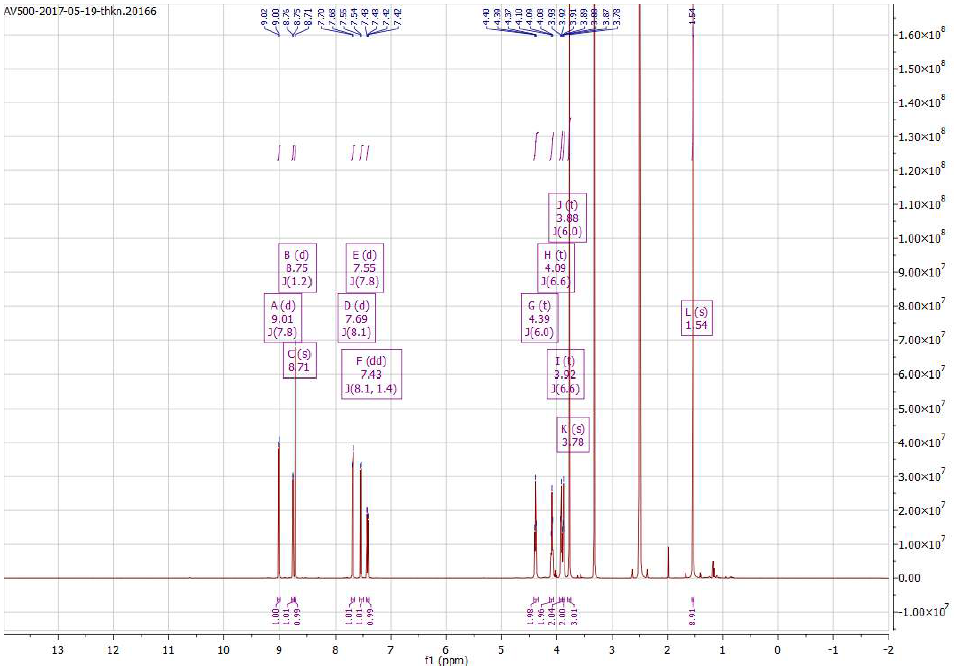
**^1^H NMR**

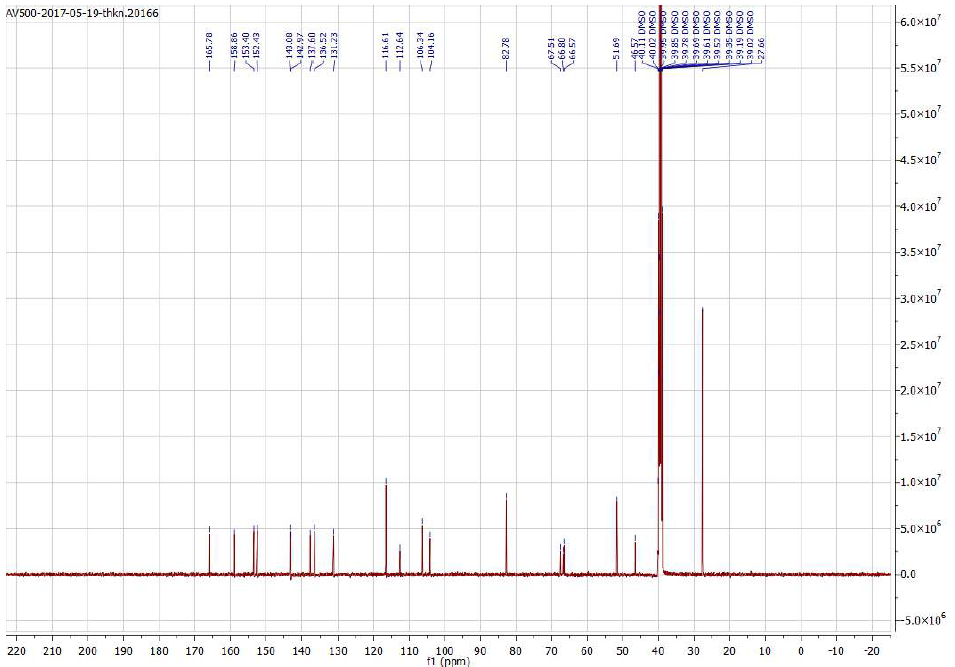
**^13^C NMR**

**Compound 11b**

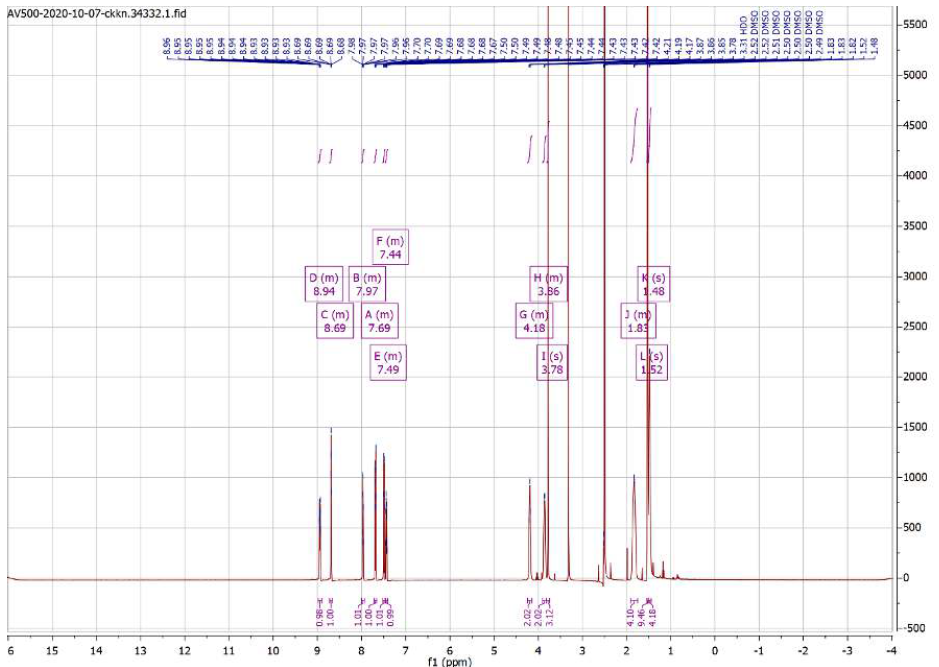
**^1^H NMR**

**^13^C NMR**

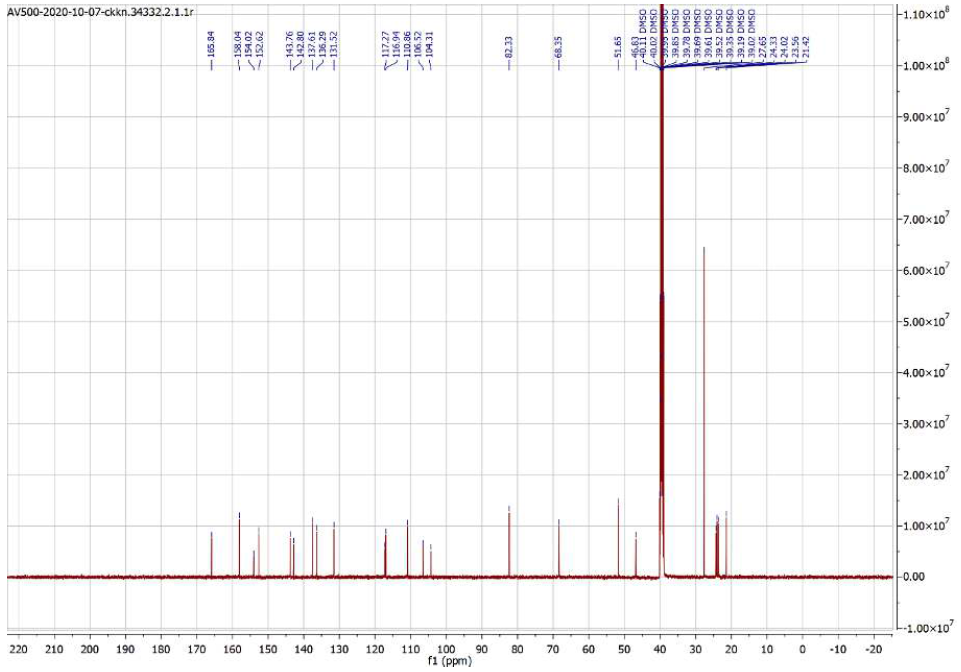

**Compound 12a**

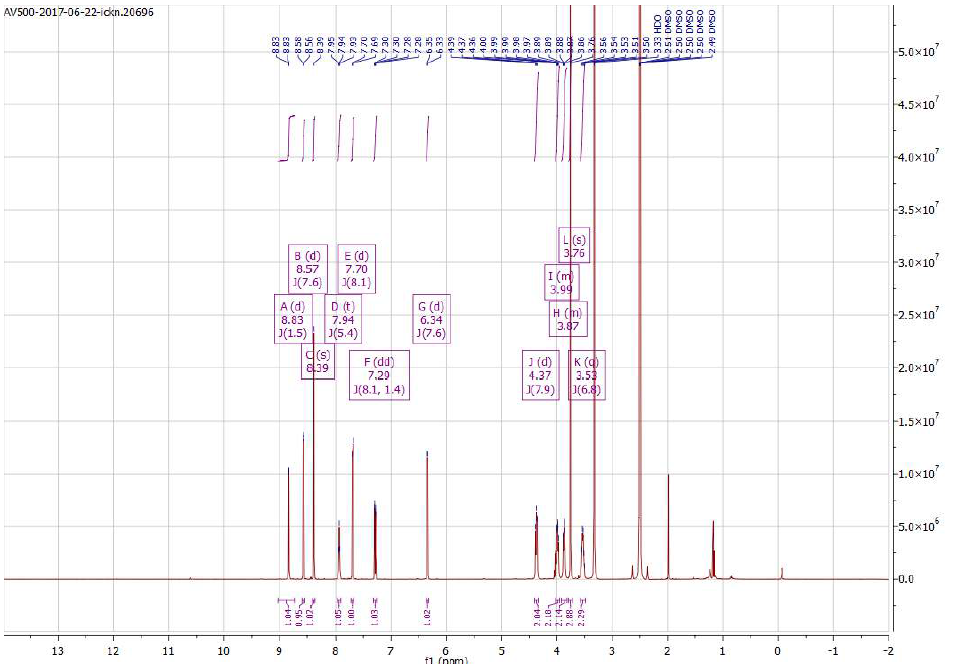
**^1^H NMR**

**^13^C NMR**

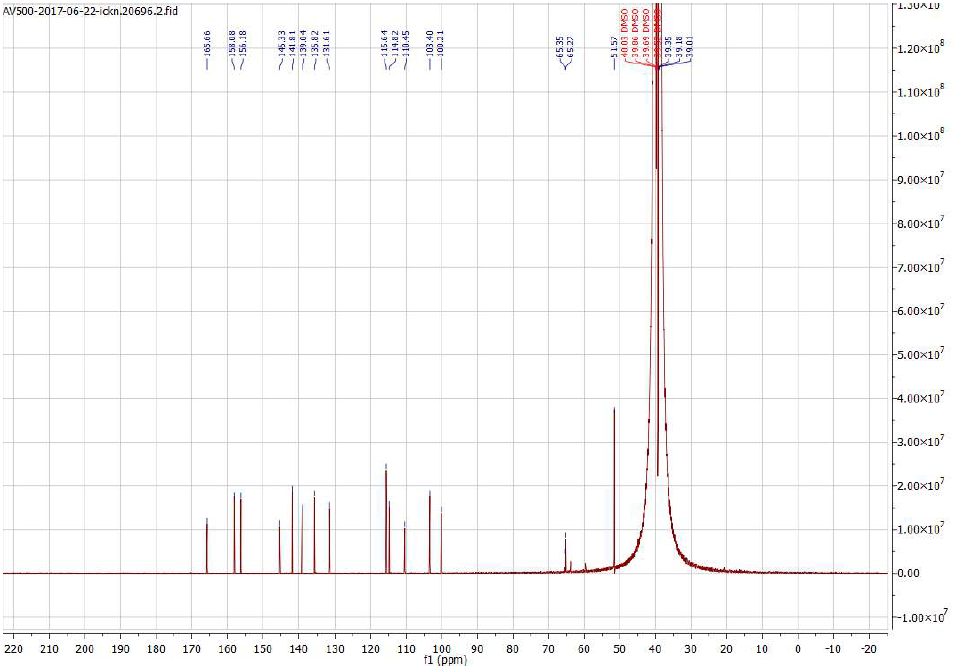

**Compound 12b**

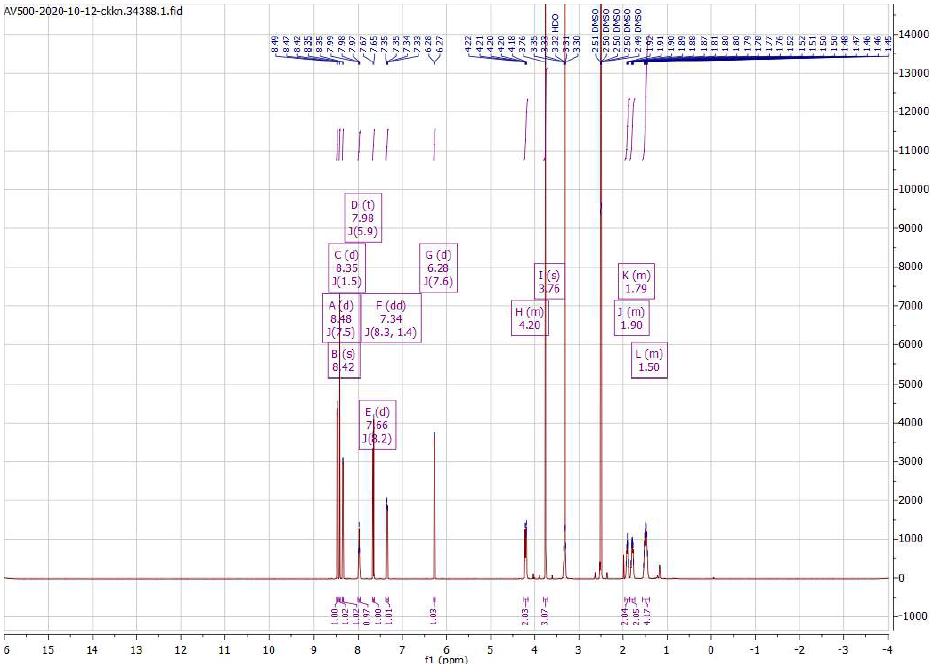
**^1^H NMR**

**^13^C NMR**

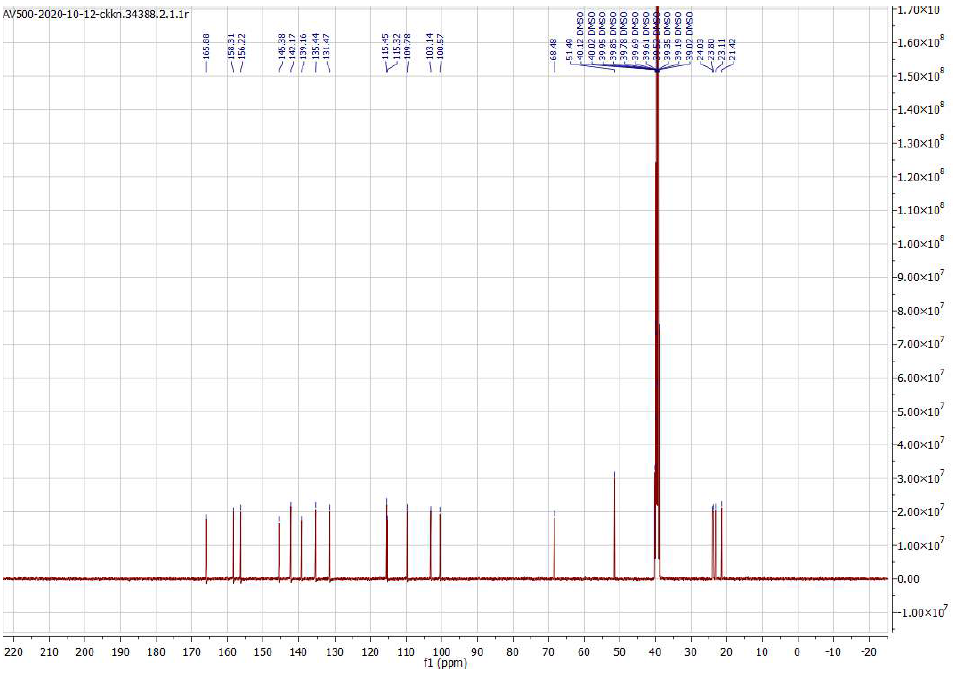

**Compound 13a**

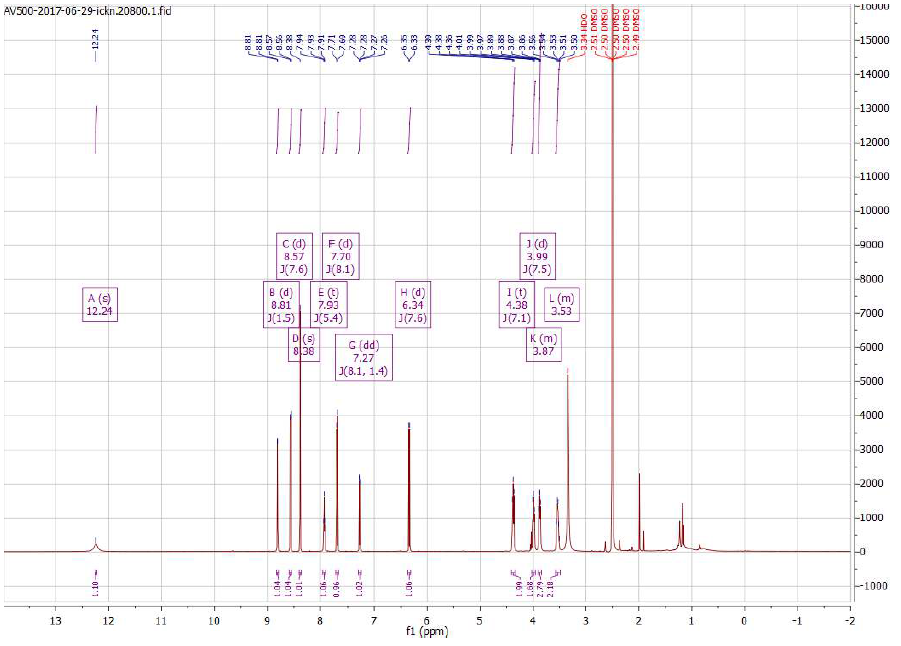
**^1^H NMR**

**^13^C NMR**

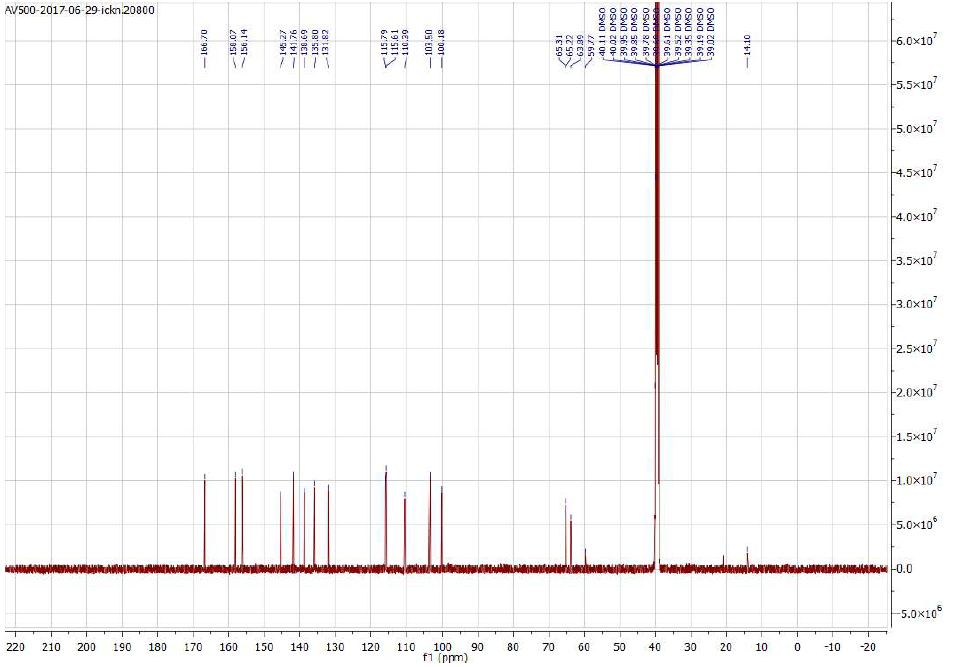

**Compound 13b**

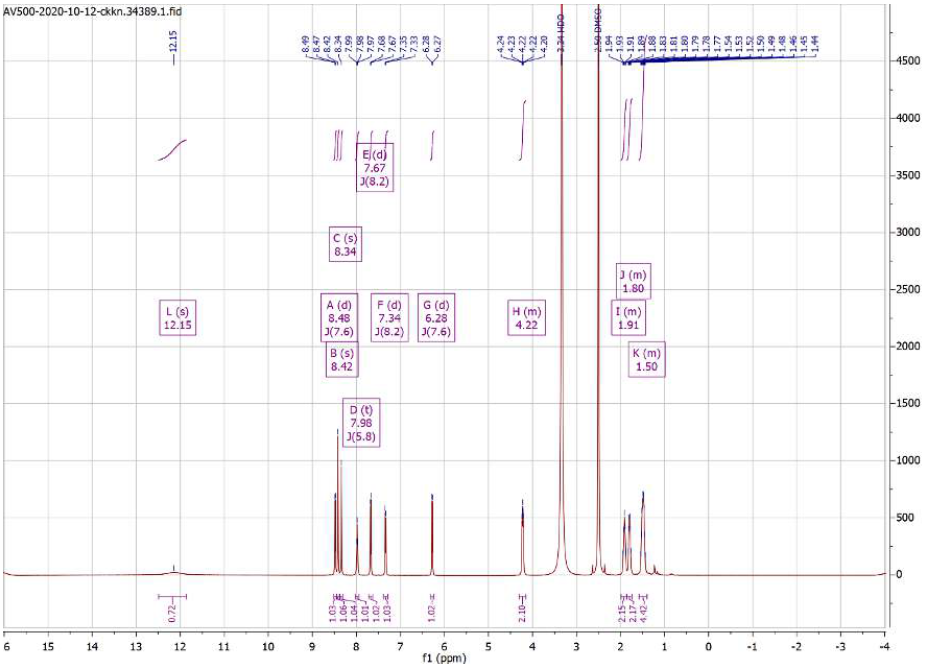
**^1^H NMR**

**^13^C NMR**

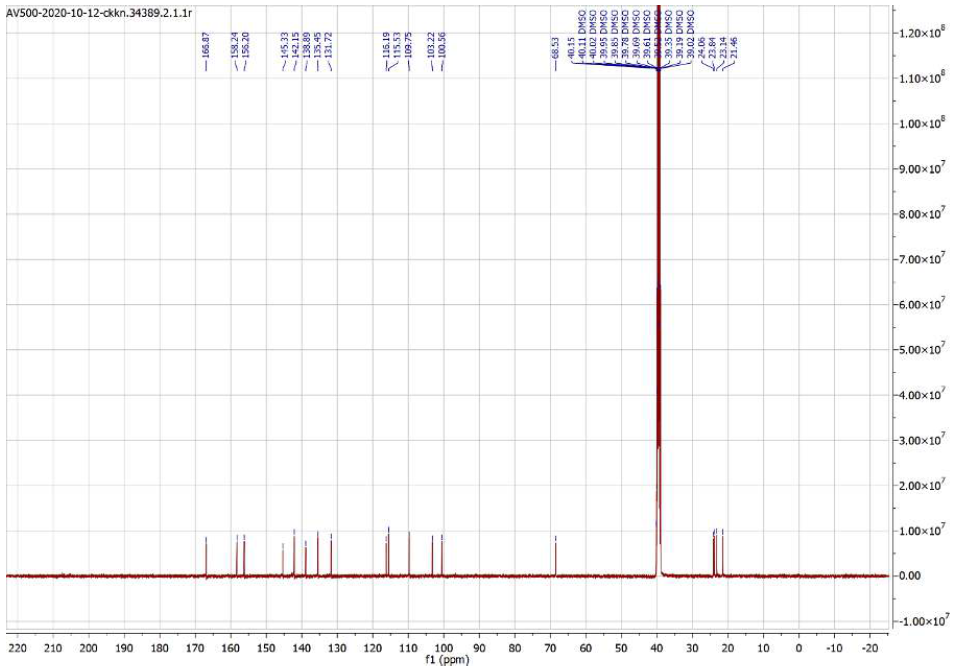

**Compound 14**

**^1^H NMR**

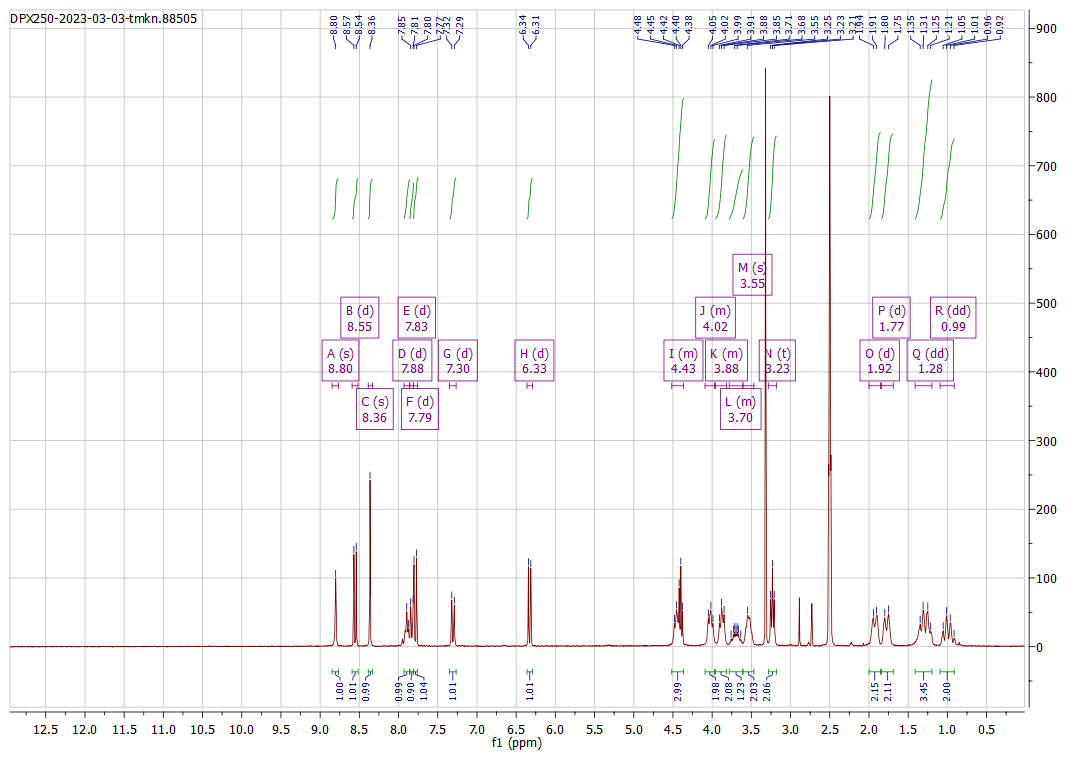

**^13^C NMR**

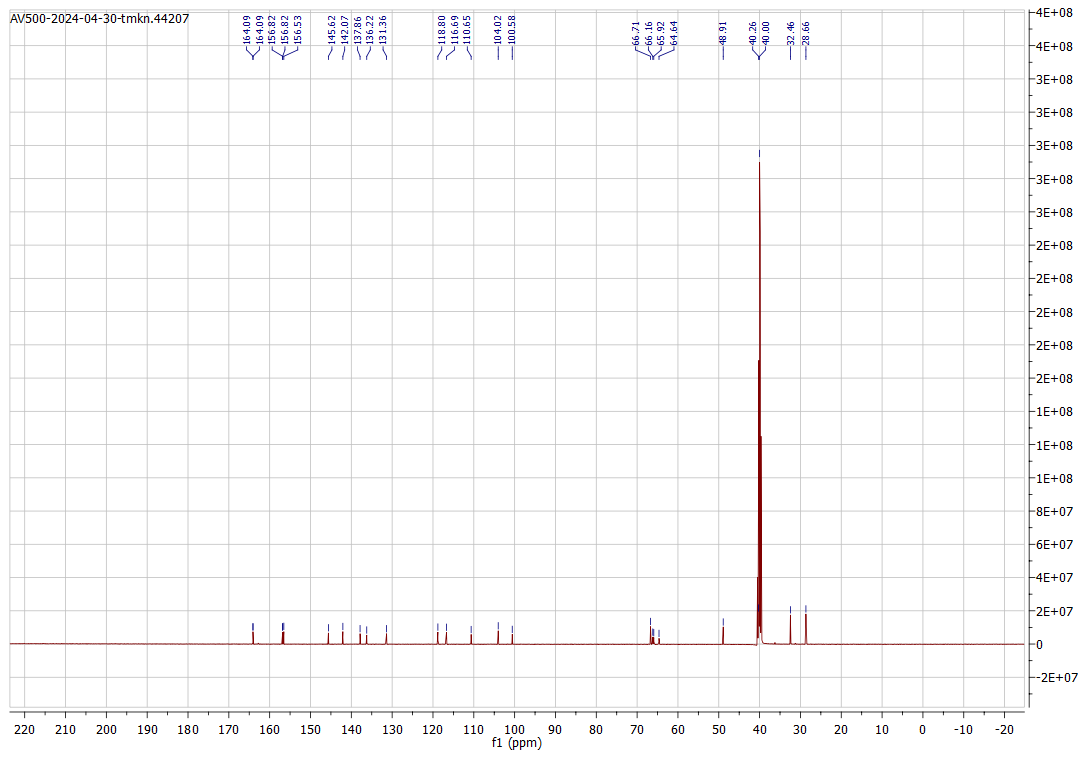

**HPLC:**

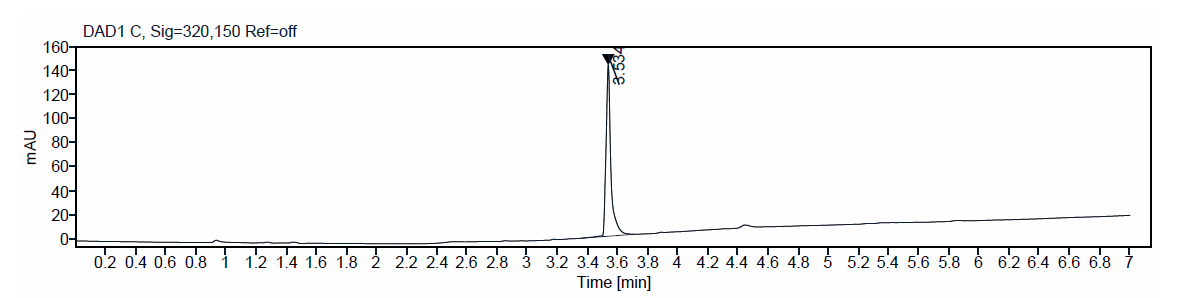

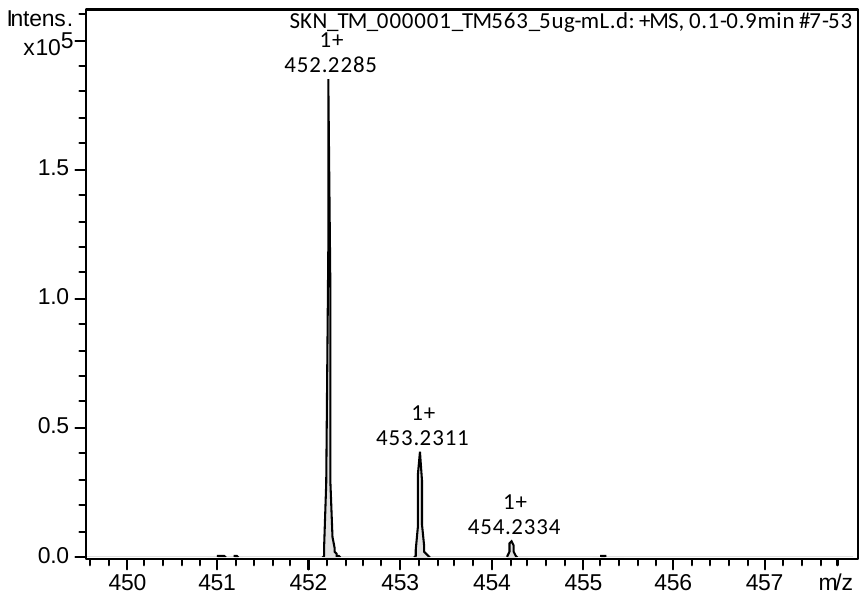
**Precision mass:**

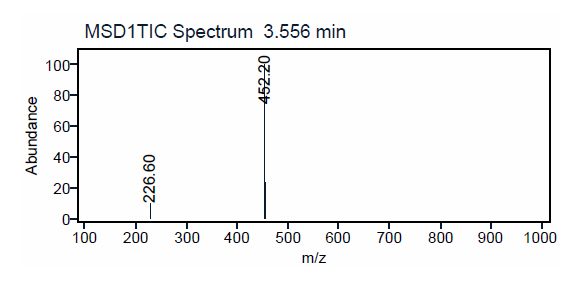
**ESI:**

**Compound 15**

**^1^H NMR**

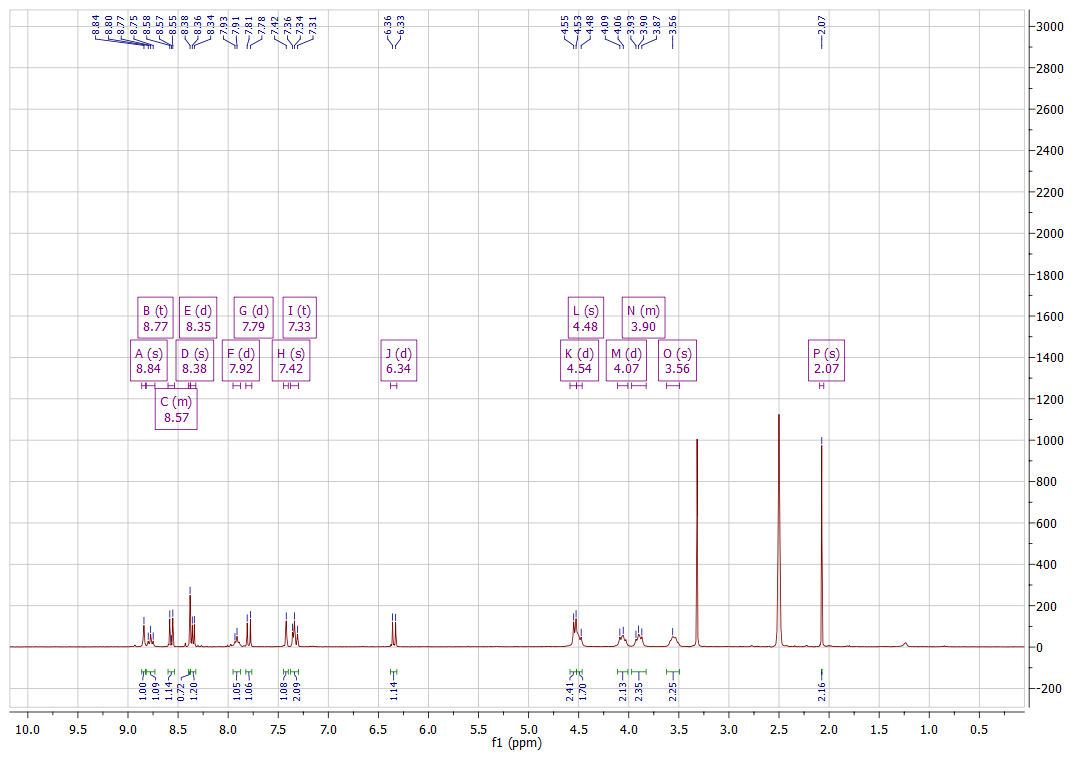

**^13^C NMR**

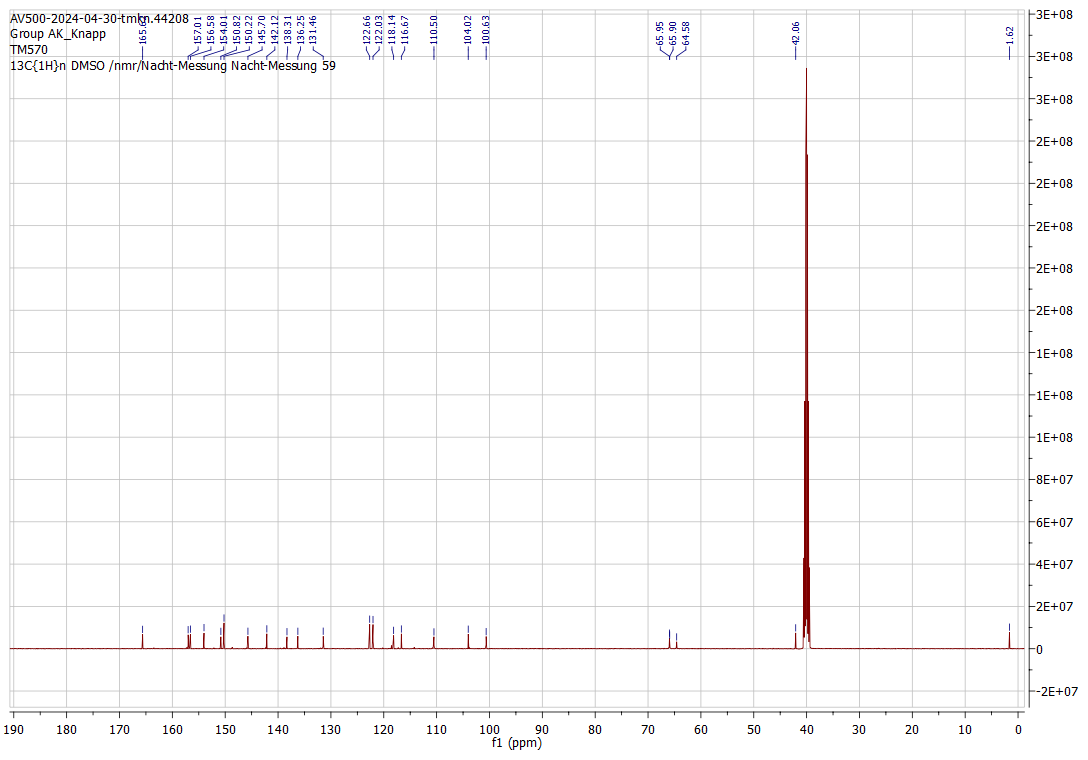

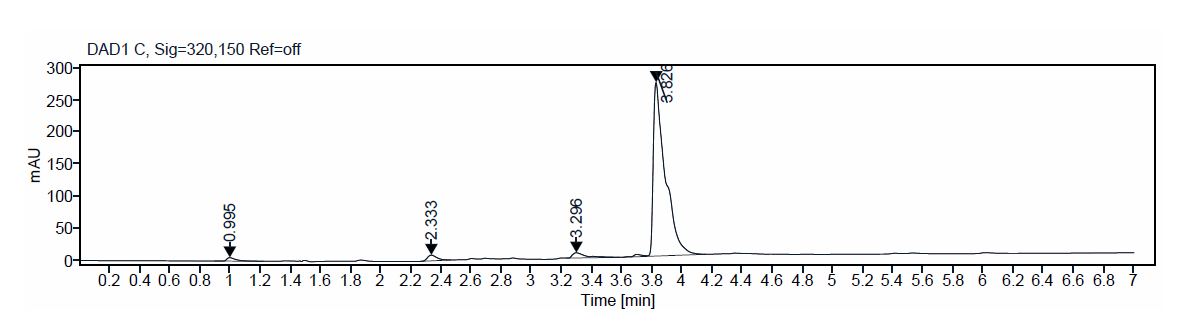
**HPLC:**

**Precision mass:**

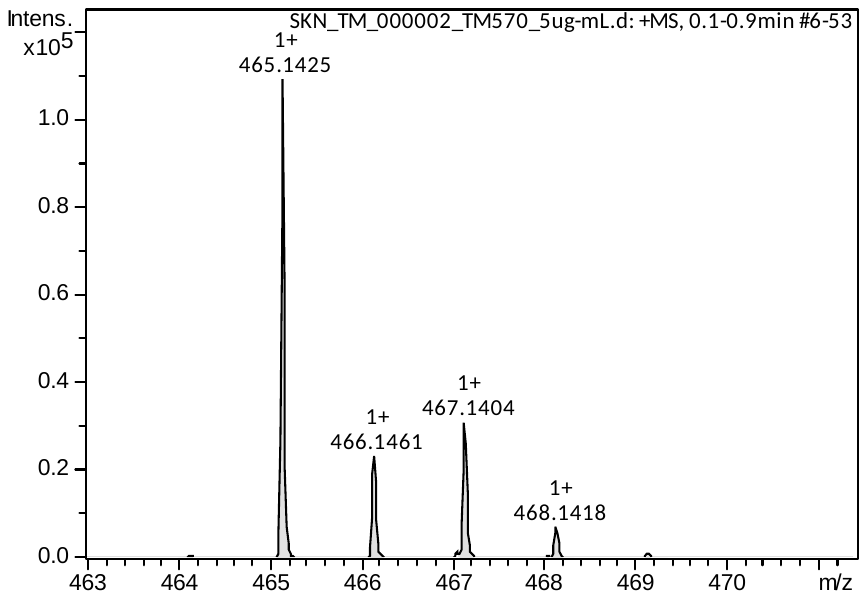

**ESI:**

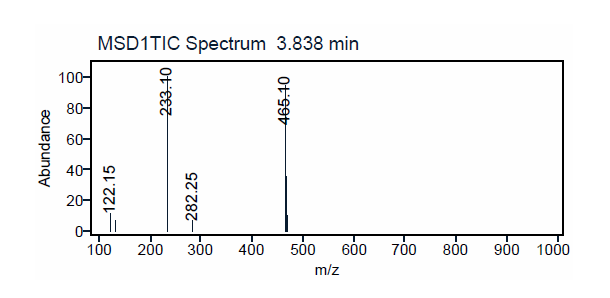

**Compound 16**

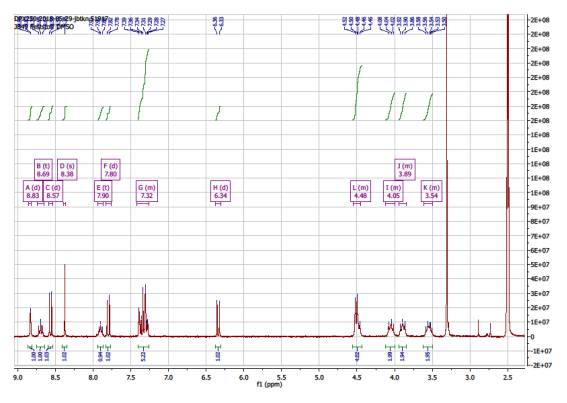
**^1^H NMR**

**^13^C NMR**

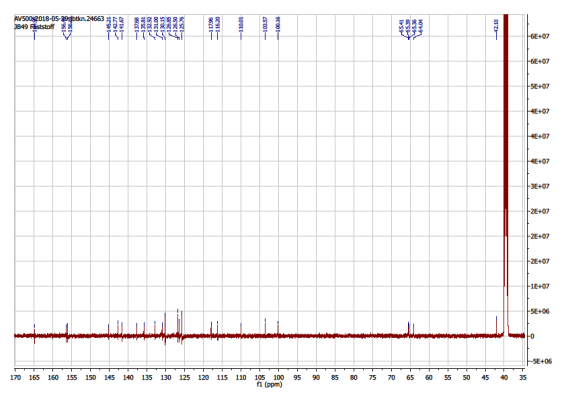

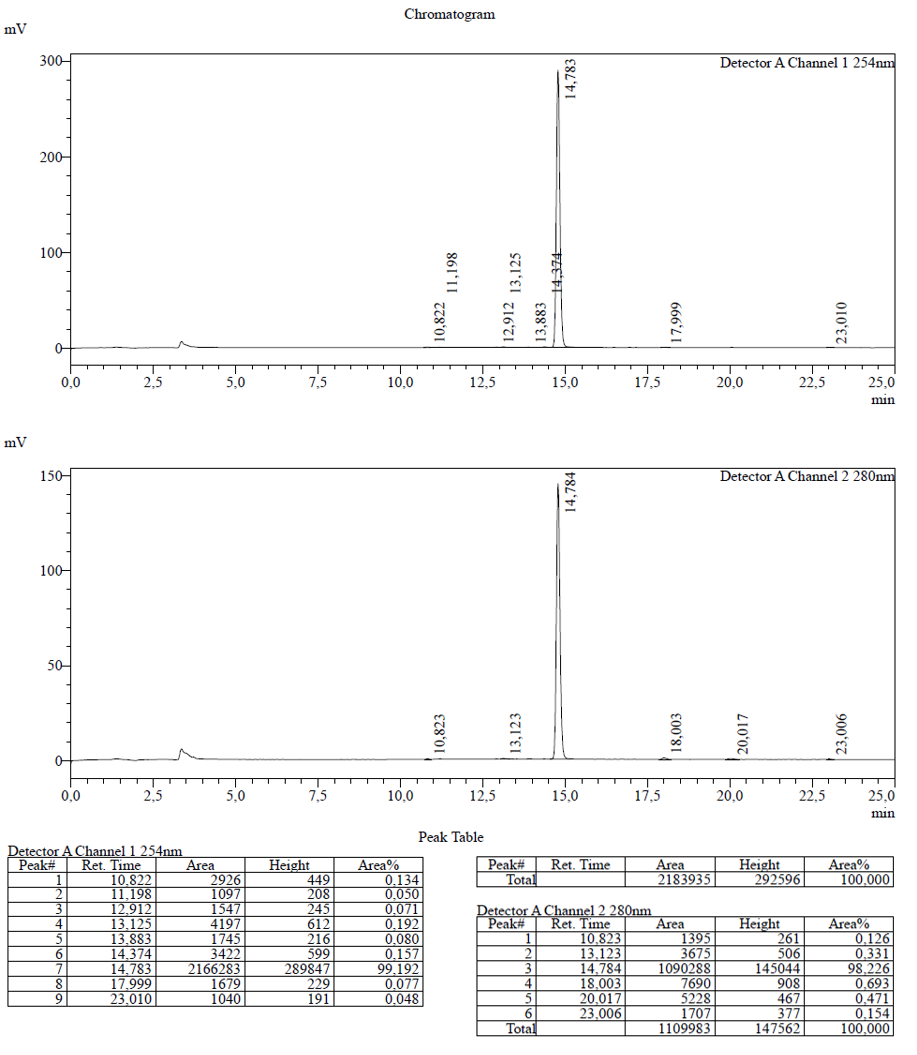
**HPLC:**

**HRMS:**
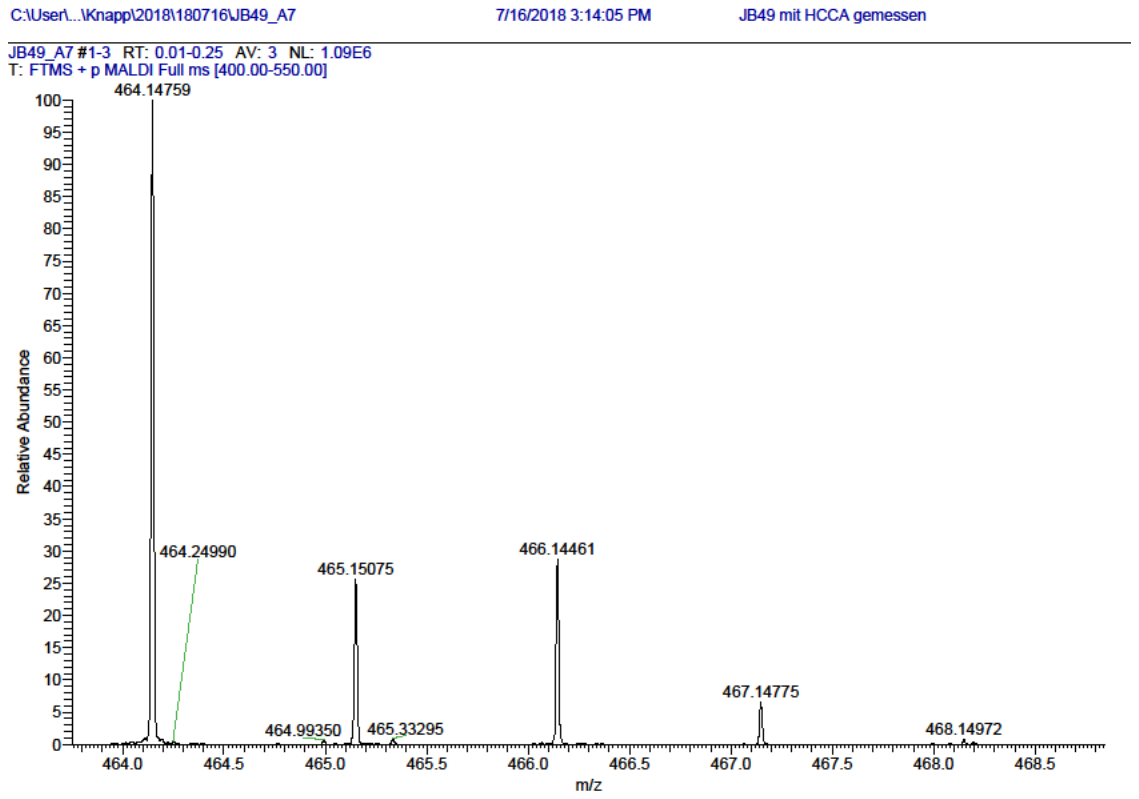

**ESI:**

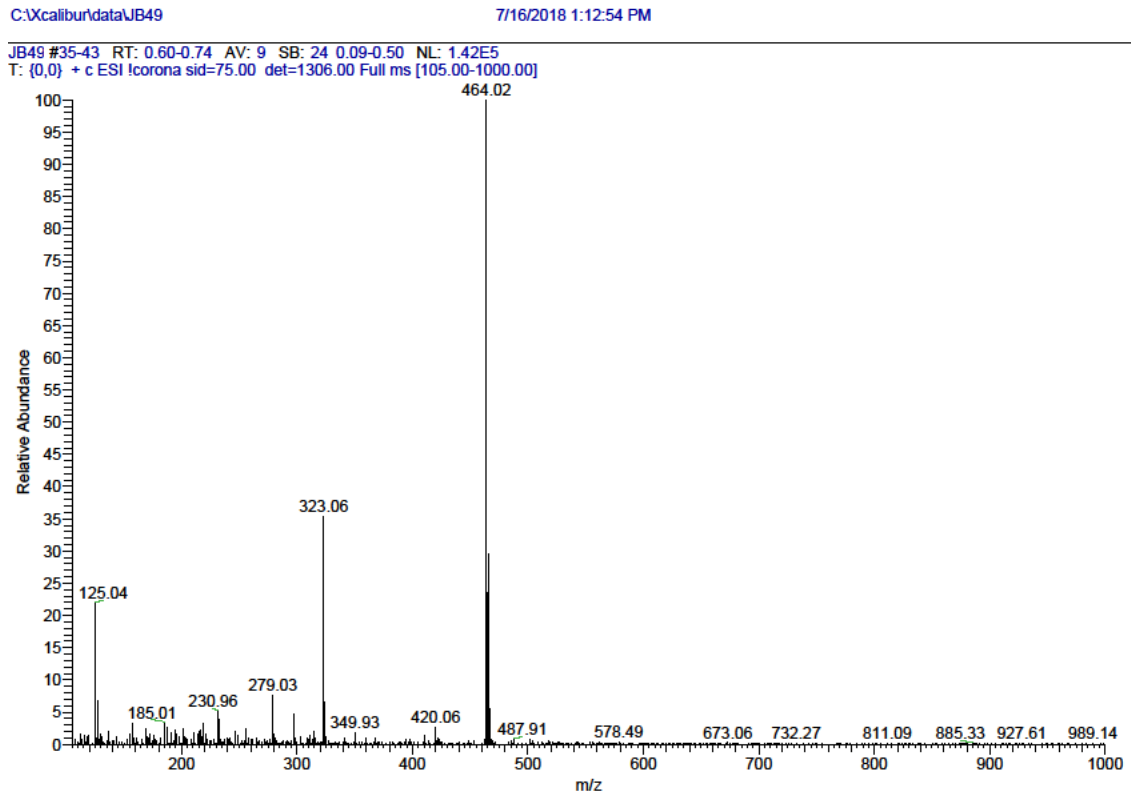

**Compound 17**

**^1^H NMR**

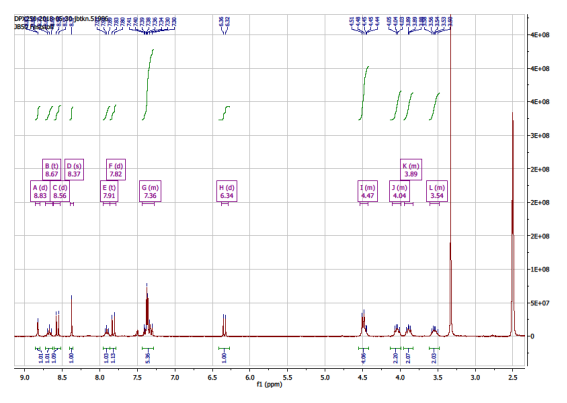

**^13^C NMR**

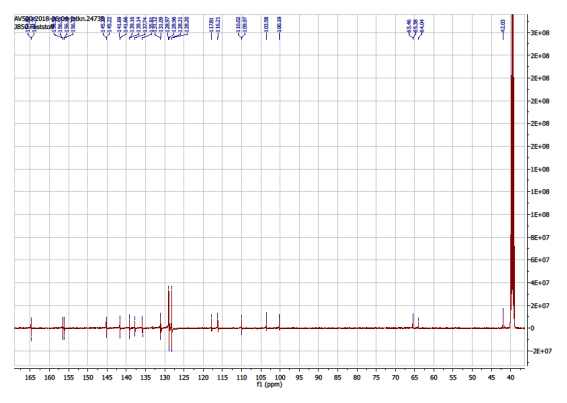

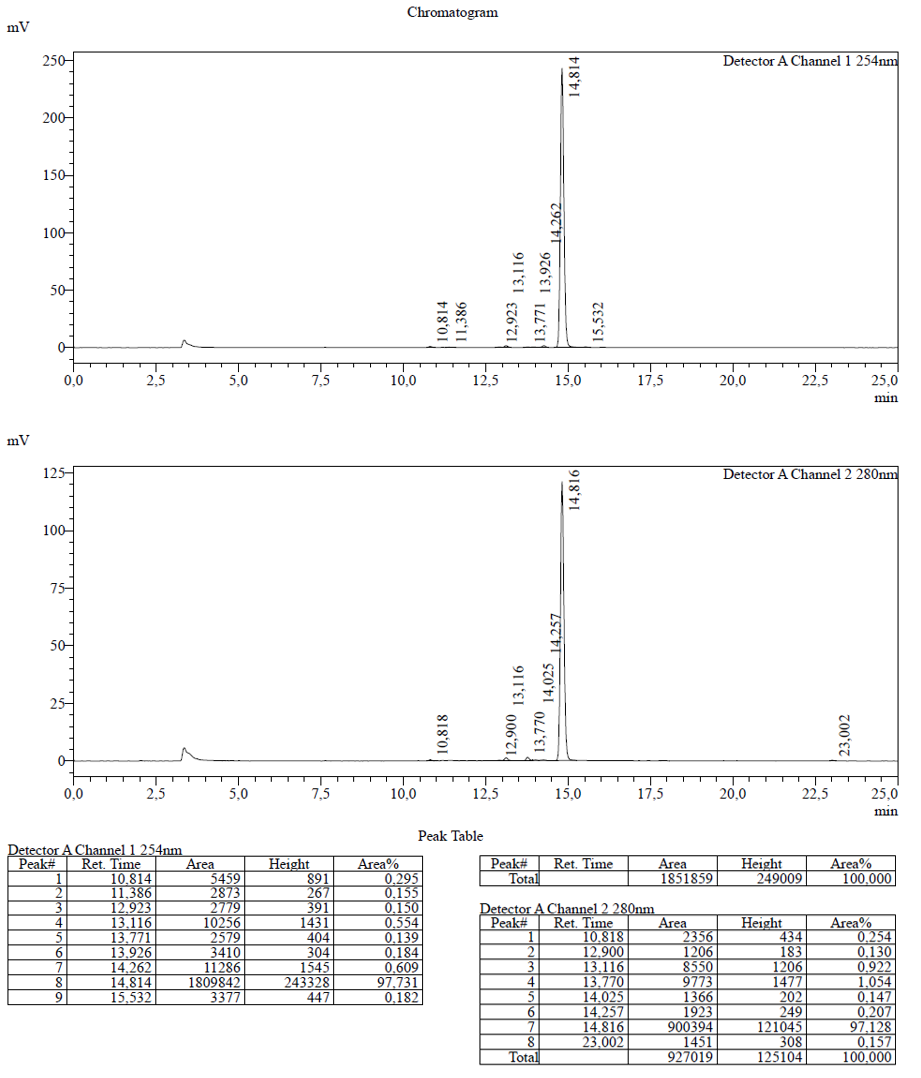
**HPLC:**

**ESI:**

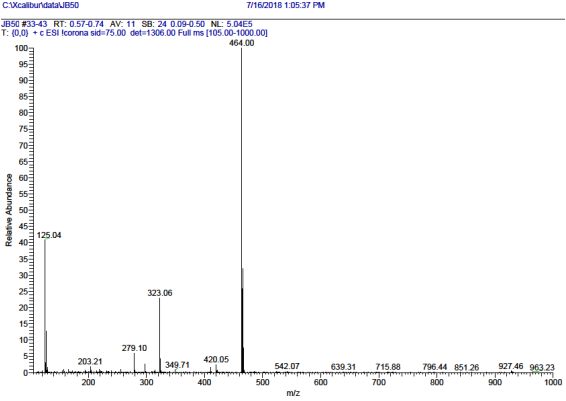

**HRMS:**

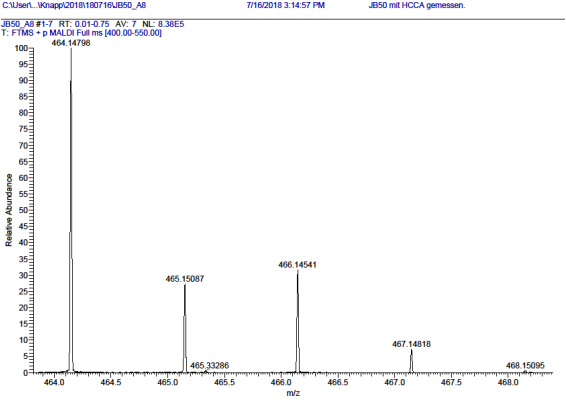

**Compound 18**

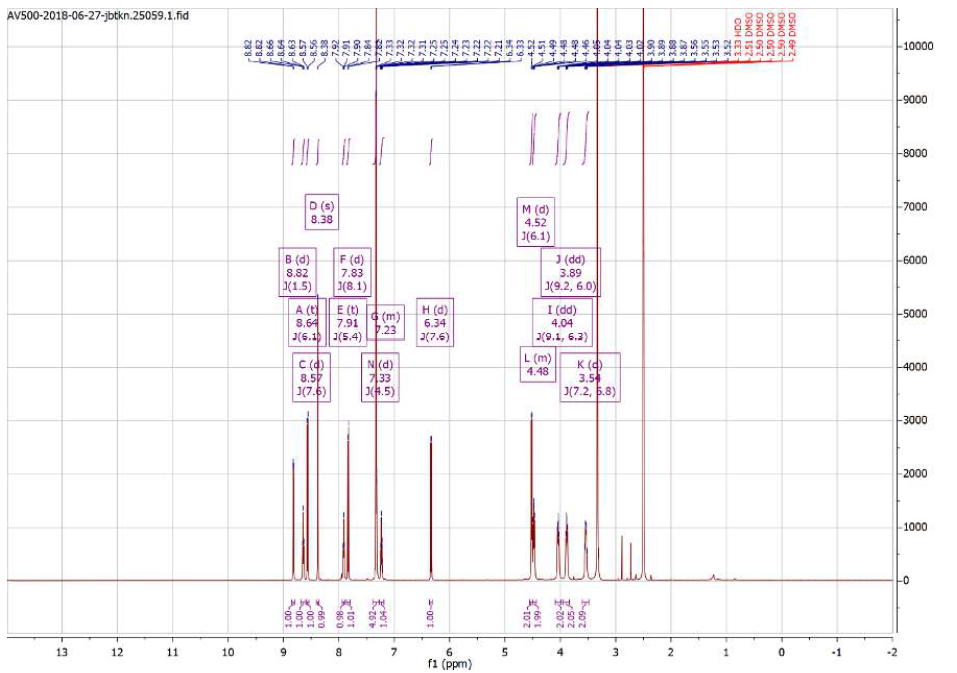
**^1^H NMR**

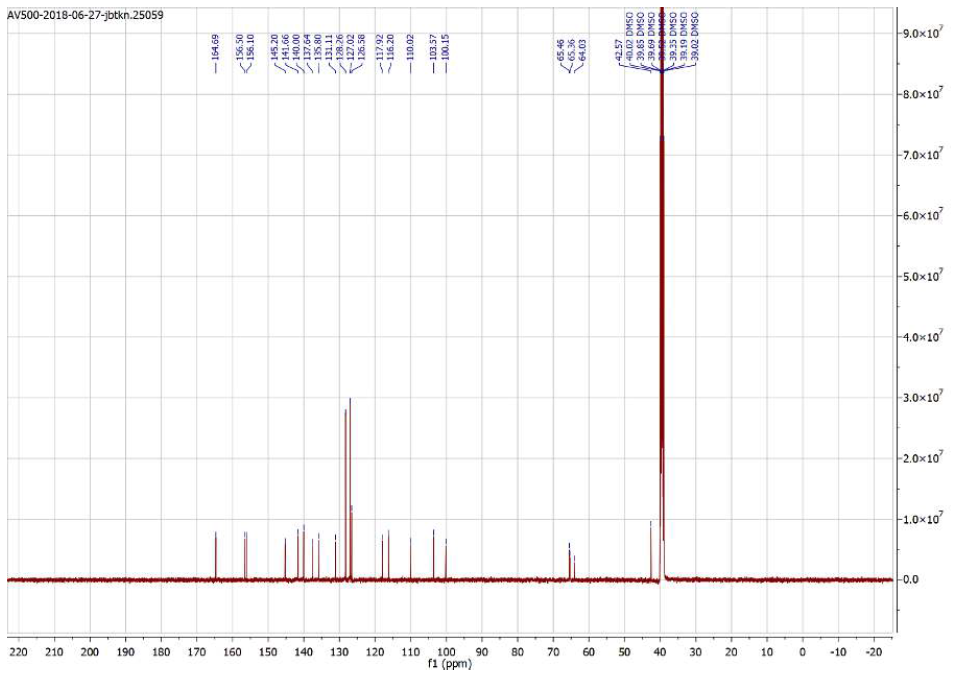
**^13^C NMR**

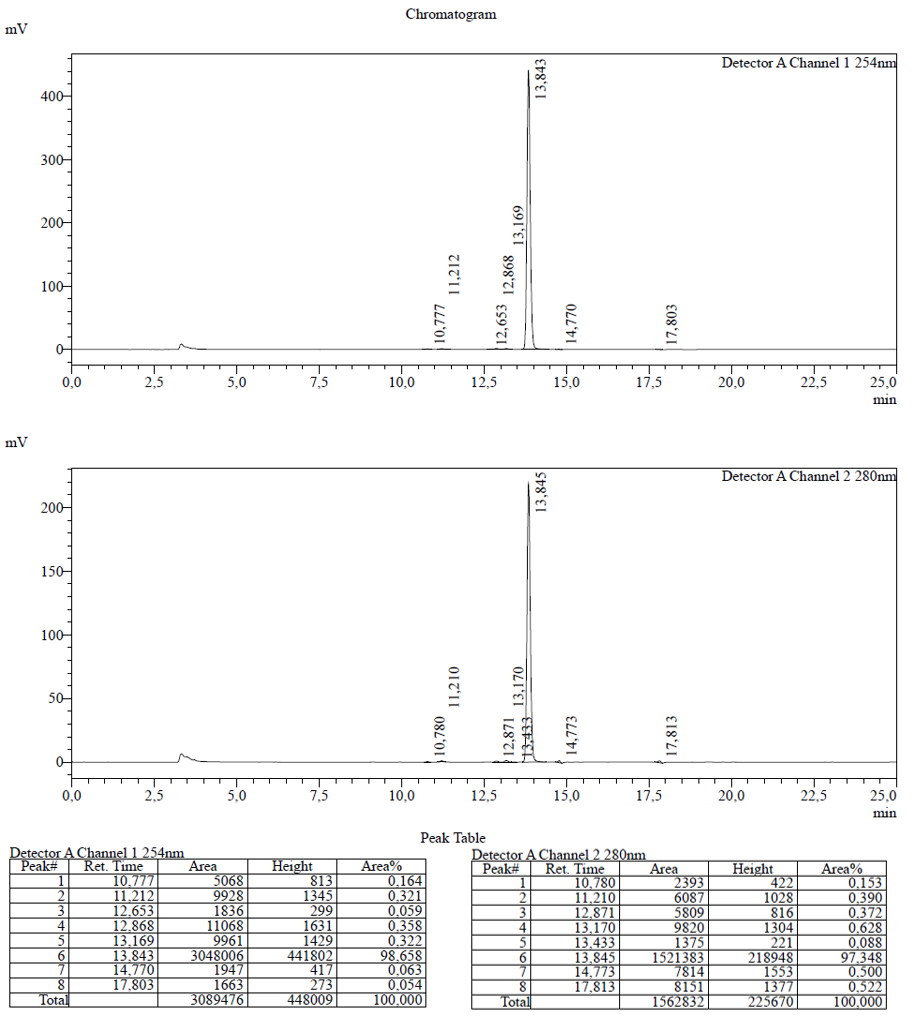
**HPLC:**

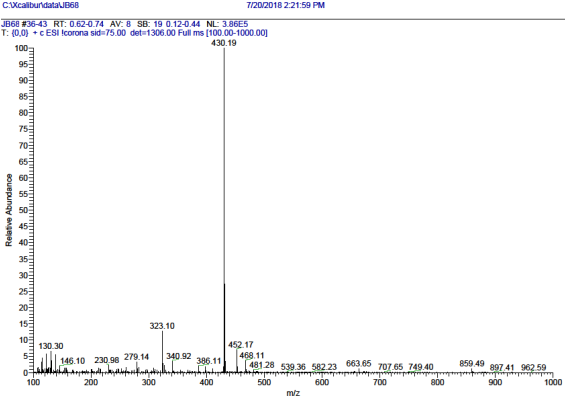
**ESI:**

**HRMS:**

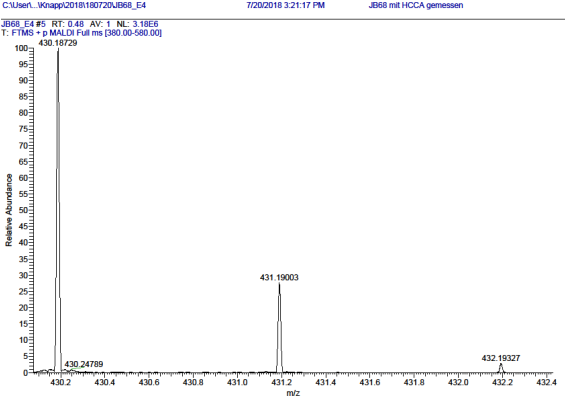

**Compound 19**

**^1^H NMR**

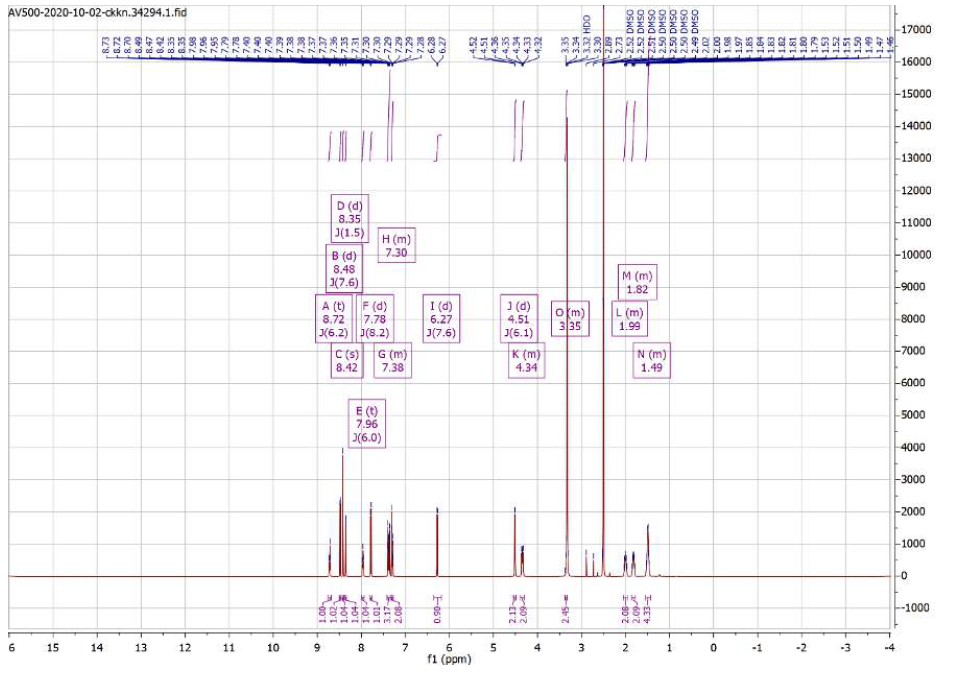

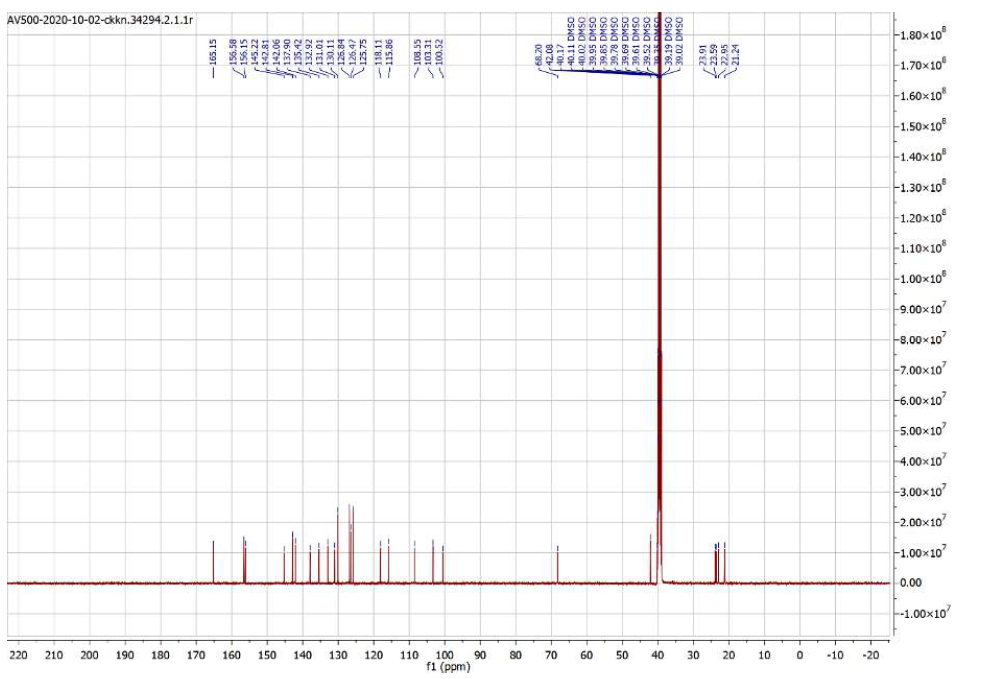
**^13^C NMR**

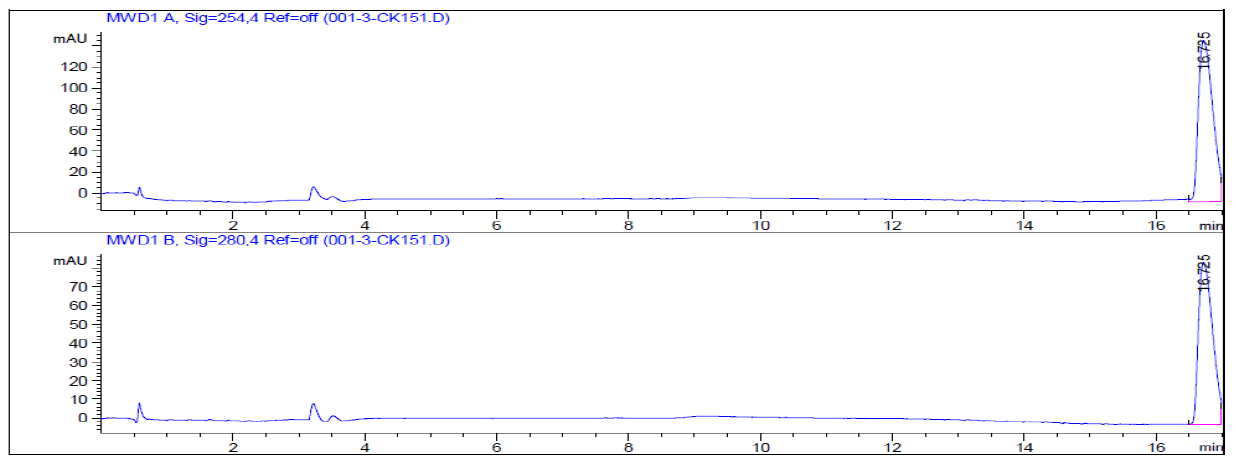
**HPLC:**

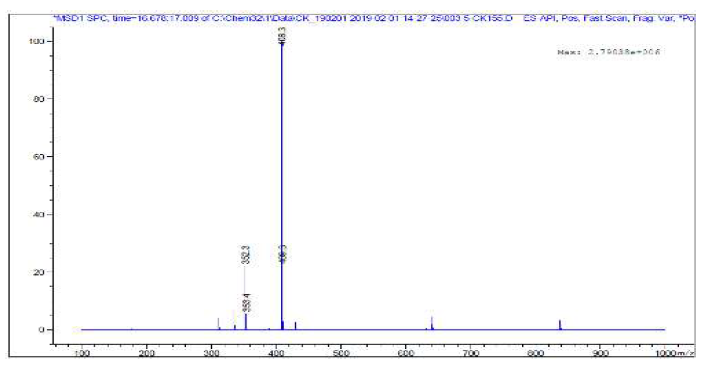

**ESI:**

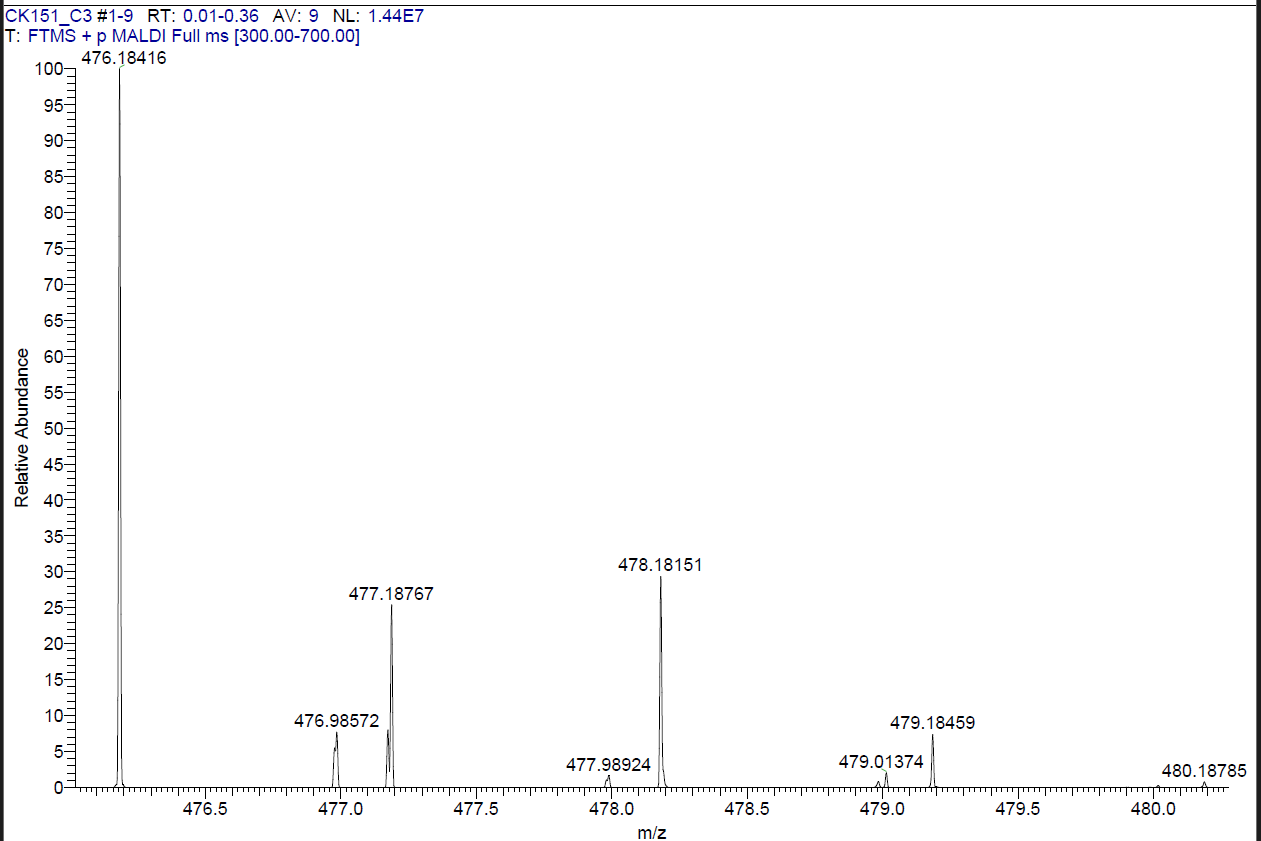
**HRMS:**

**Compound 20**

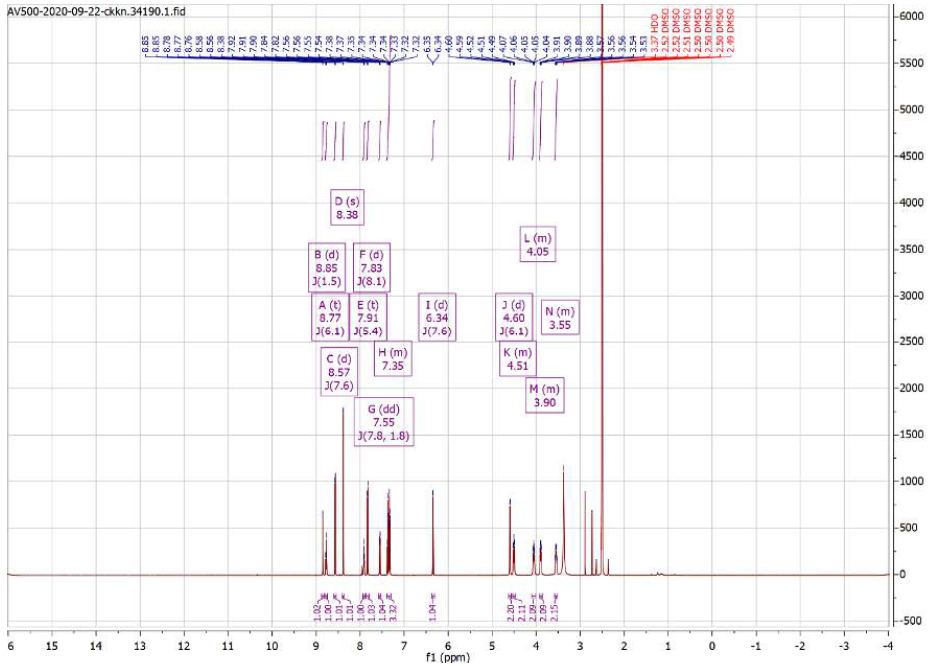
**^1^H NMR**

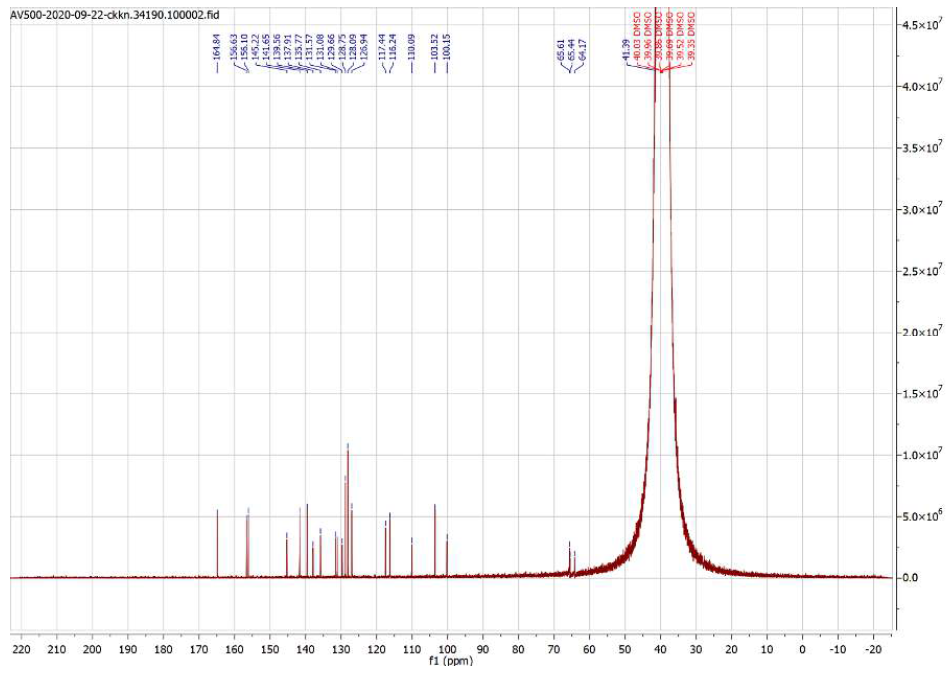
**^13^C NMR**

**HPLC:**

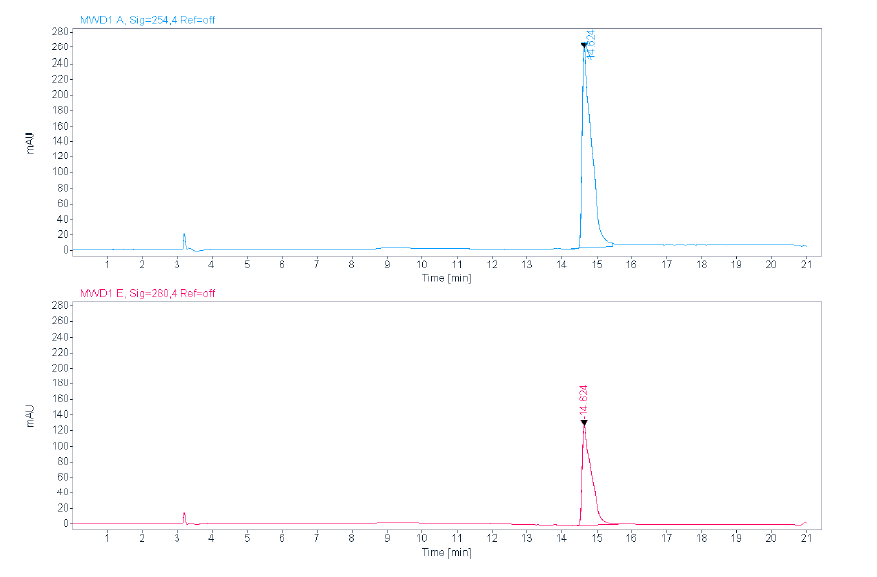

**ESI:**

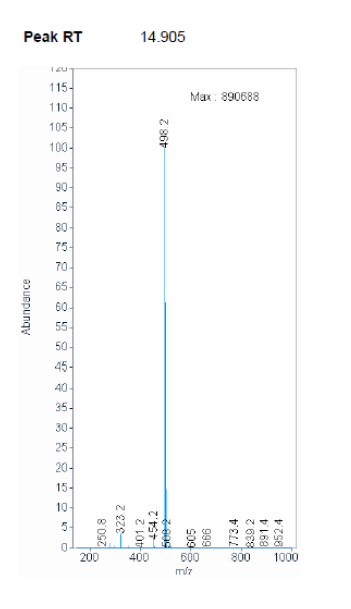

**HRMS:**

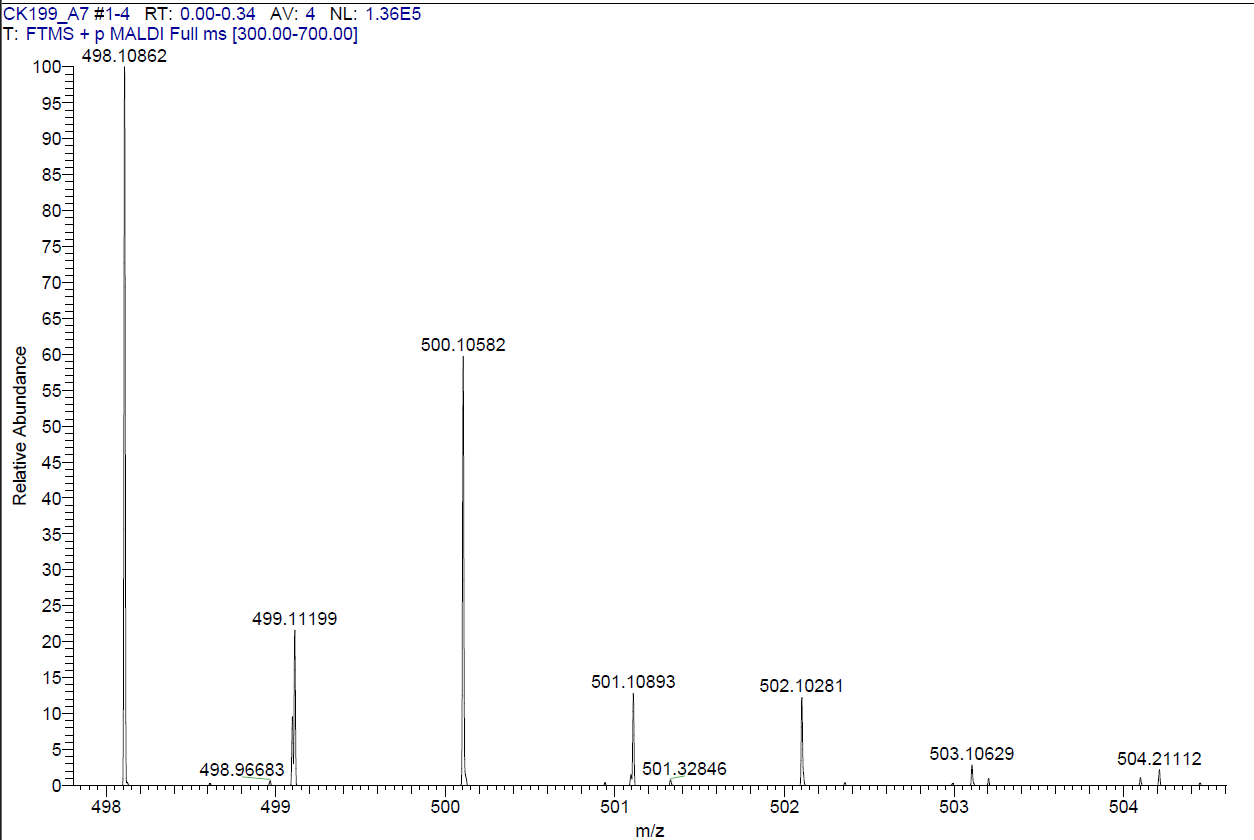

**Compound 21**

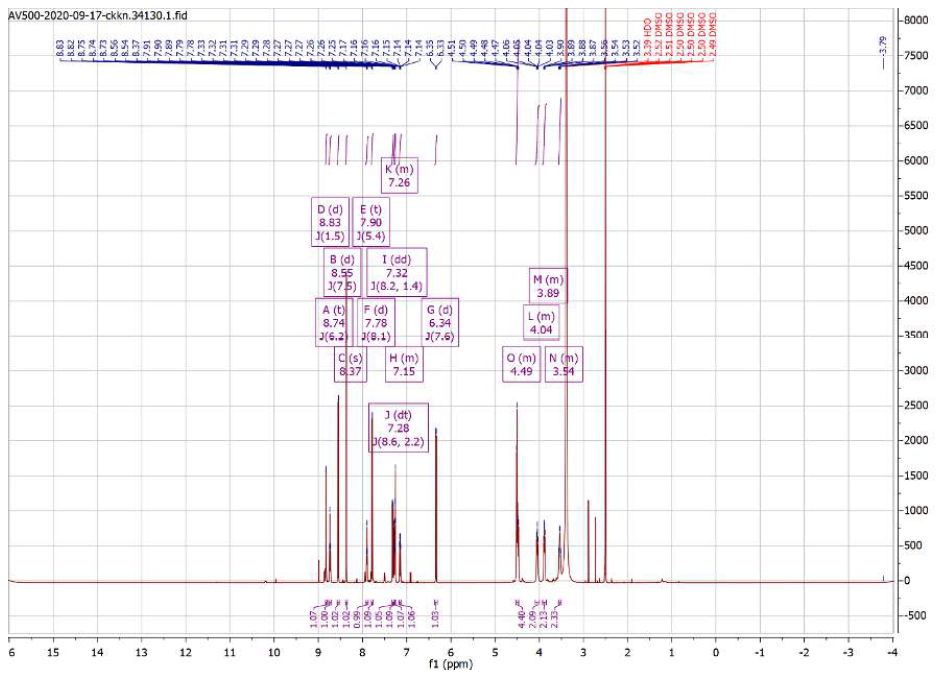
**^1^H NMR**

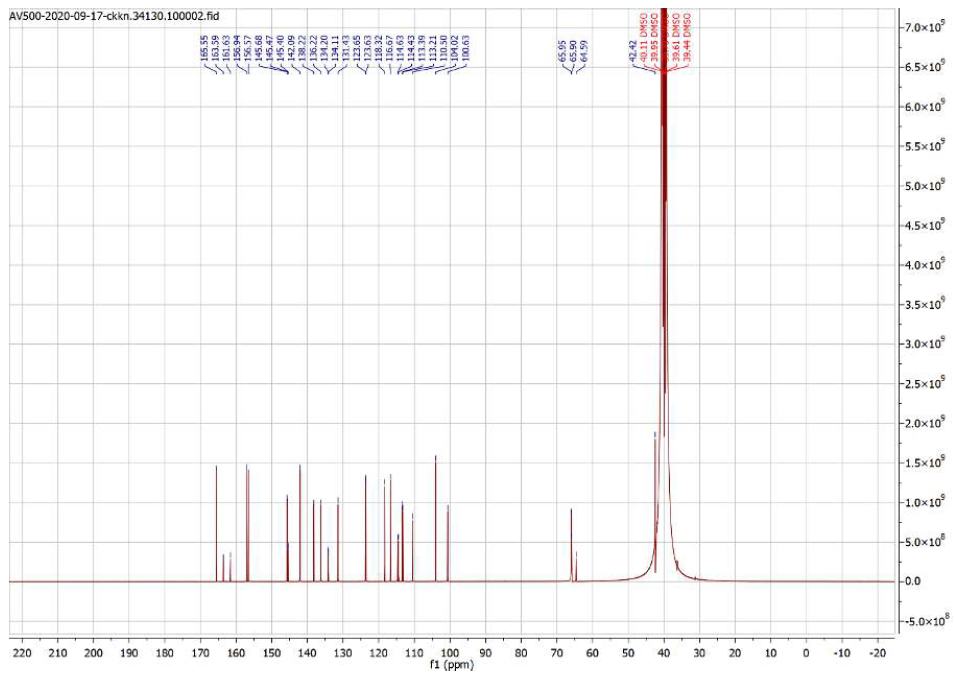
**^13^C NMR**

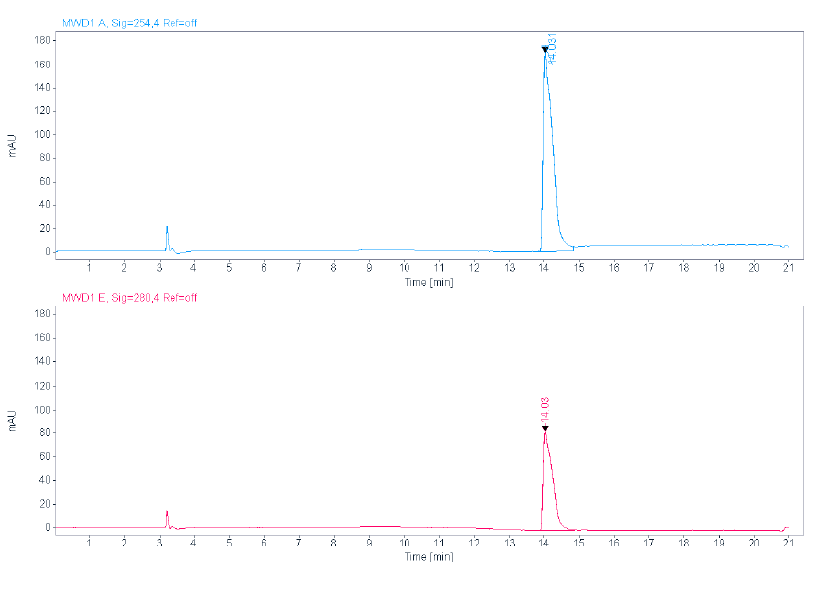
**HPLC:**

**ESI:**

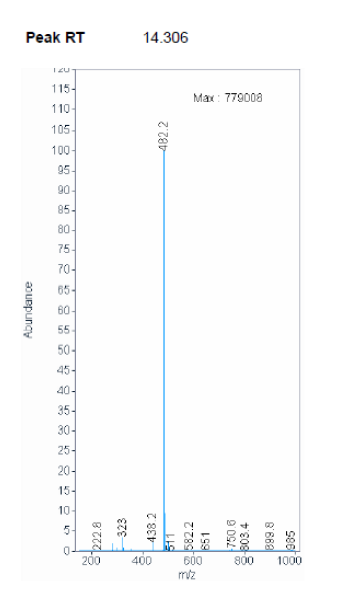

**HRMS:**

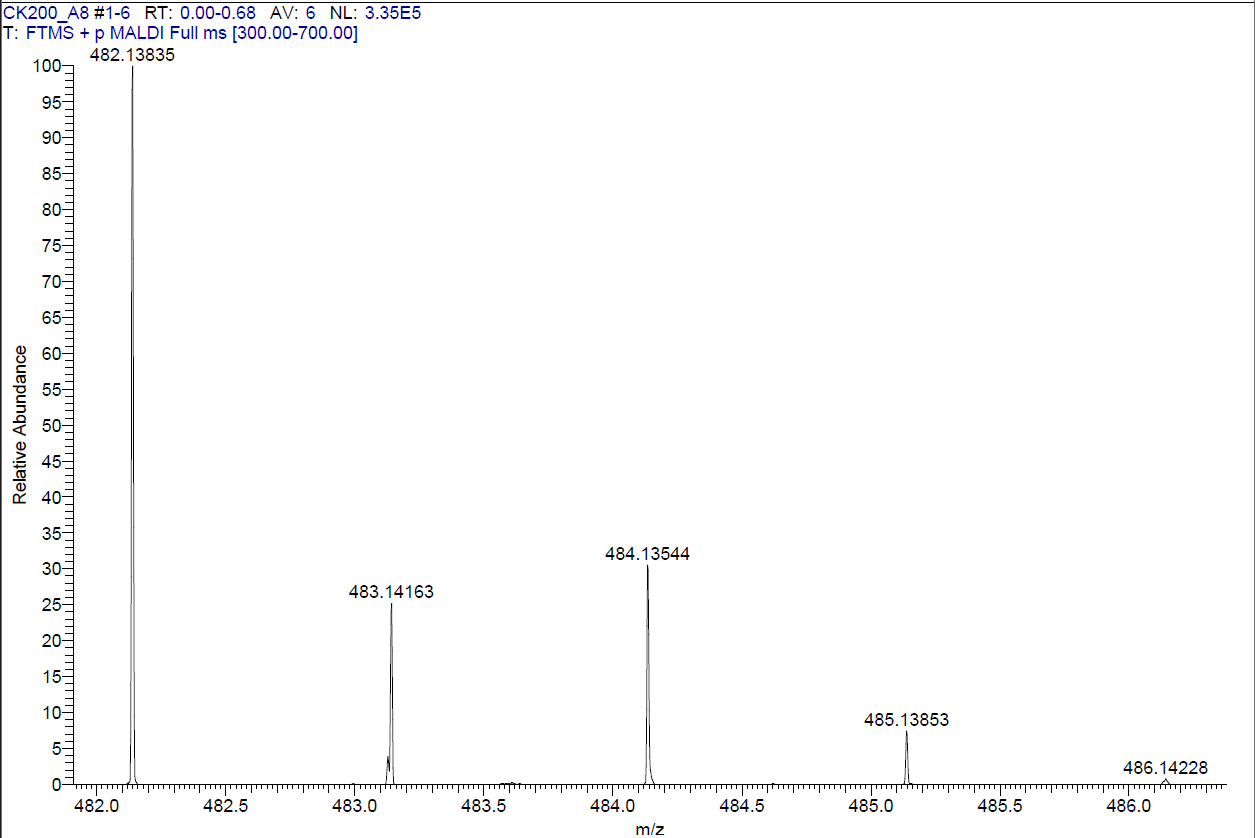

**Compound 22**

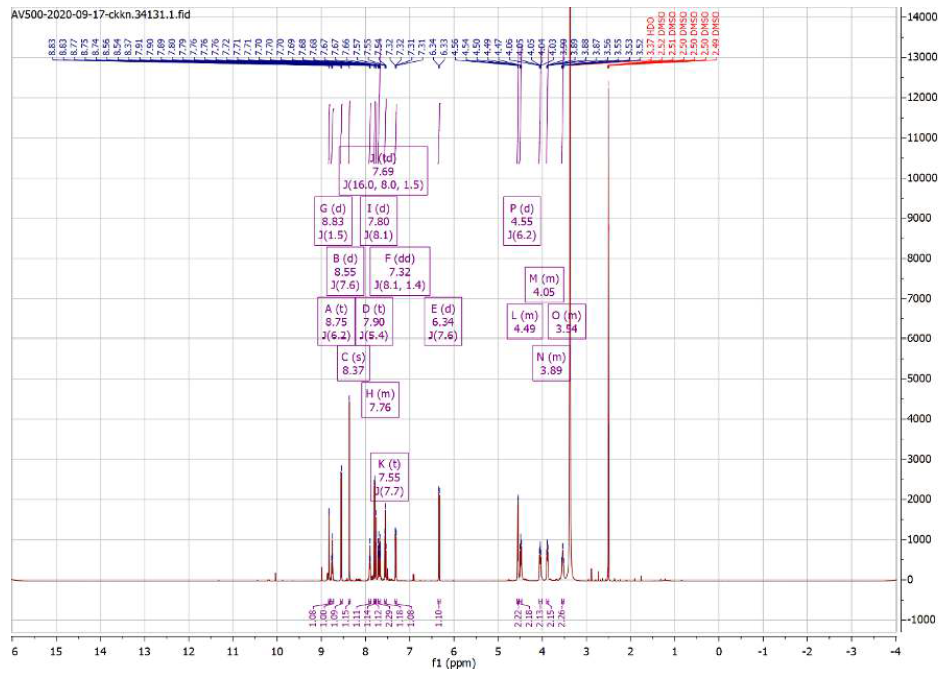
**^1^H NMR**

**^13^C NMR**

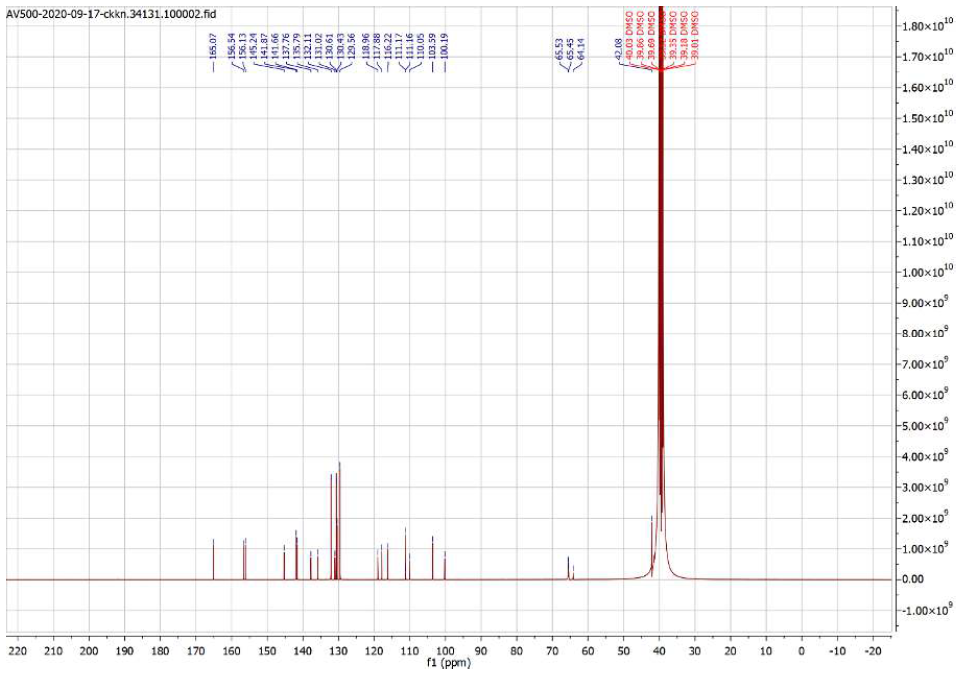

**HPLC:**

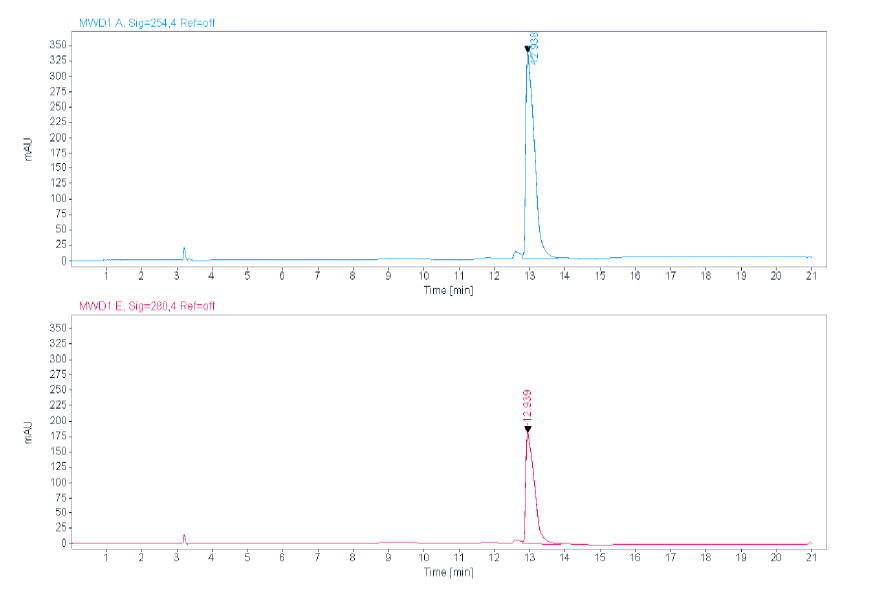

**ESI:**

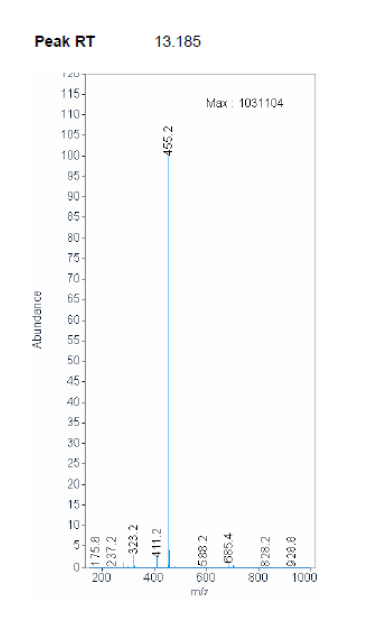

**HRMS:**

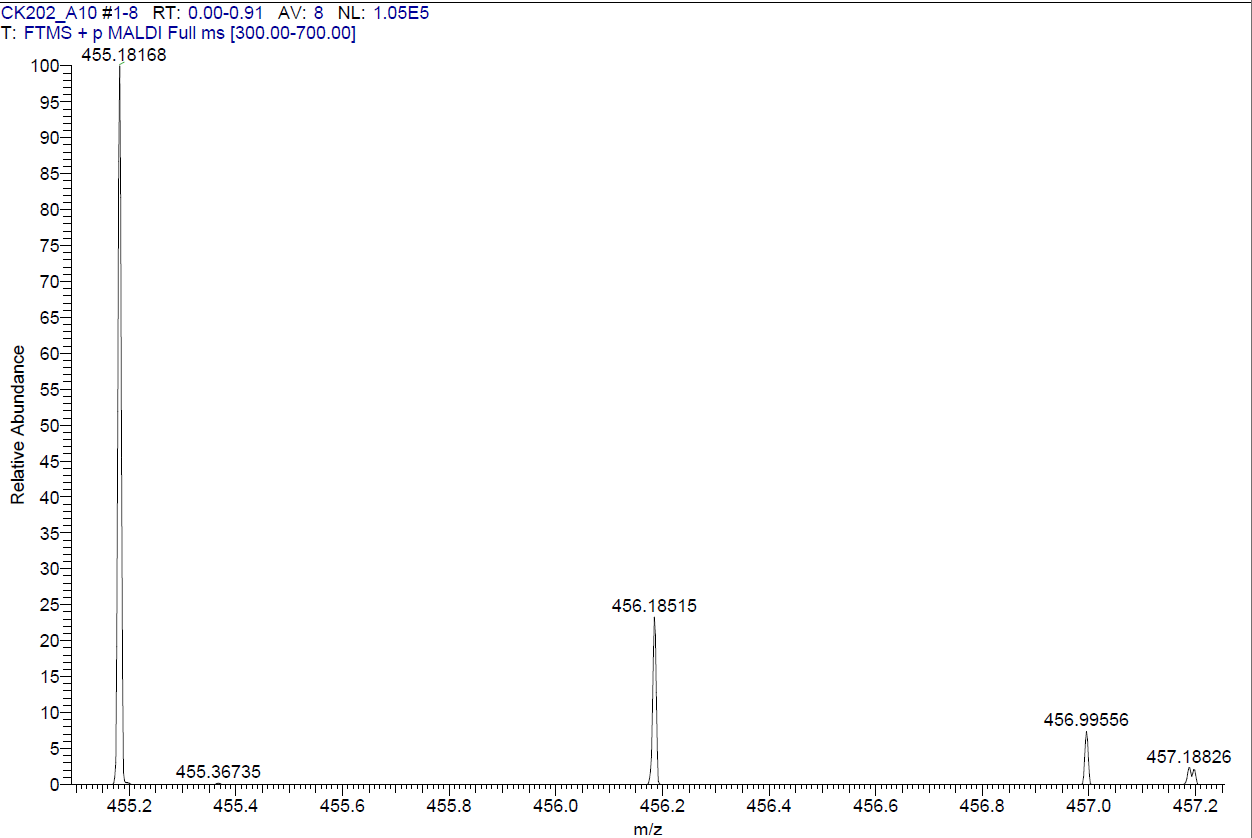

**Compound 23**

**^1^H NMR**

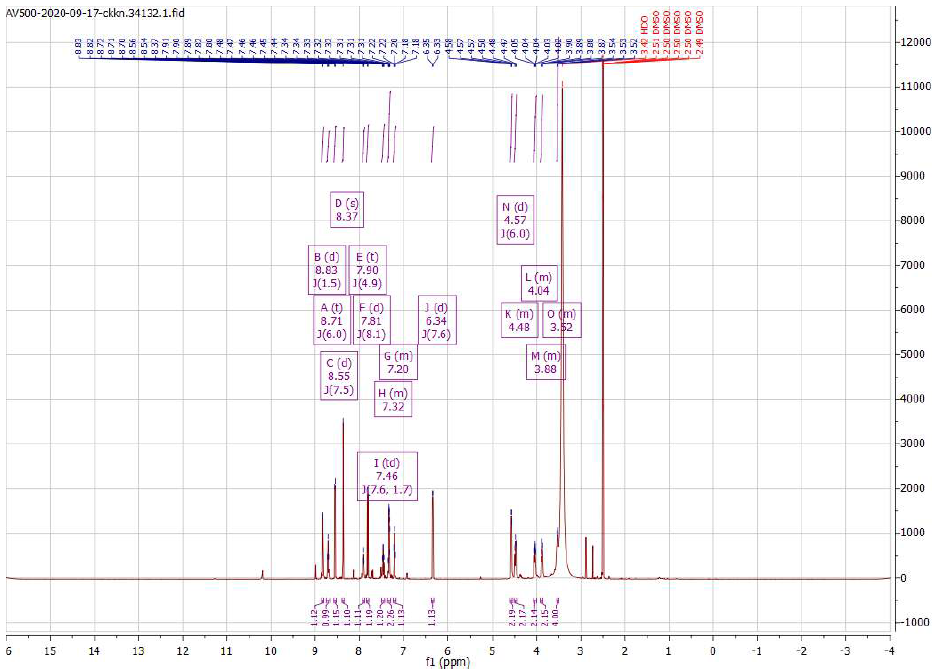

**^13^C NMR**

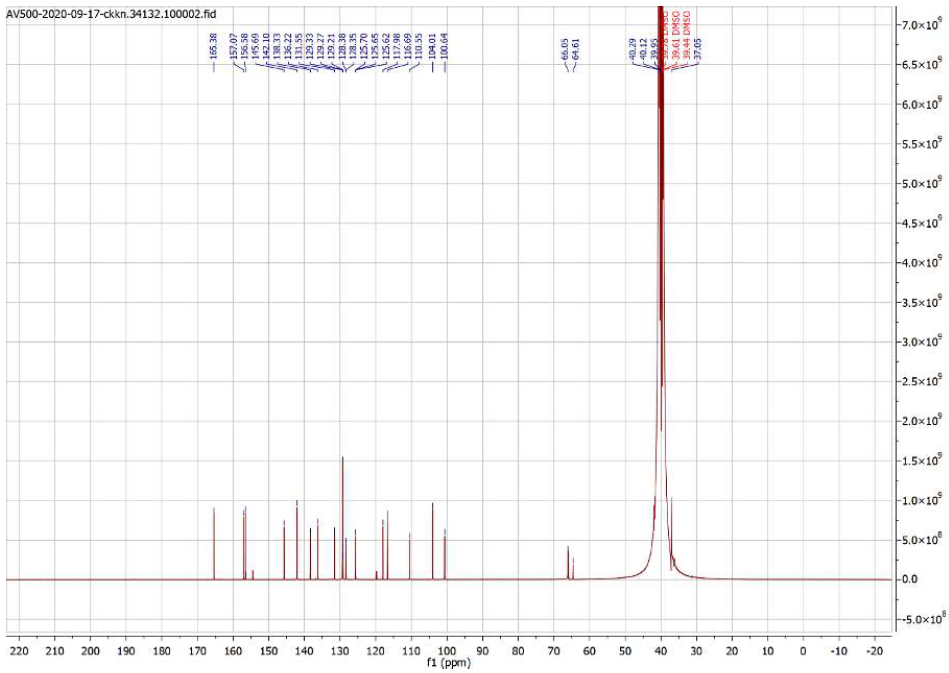

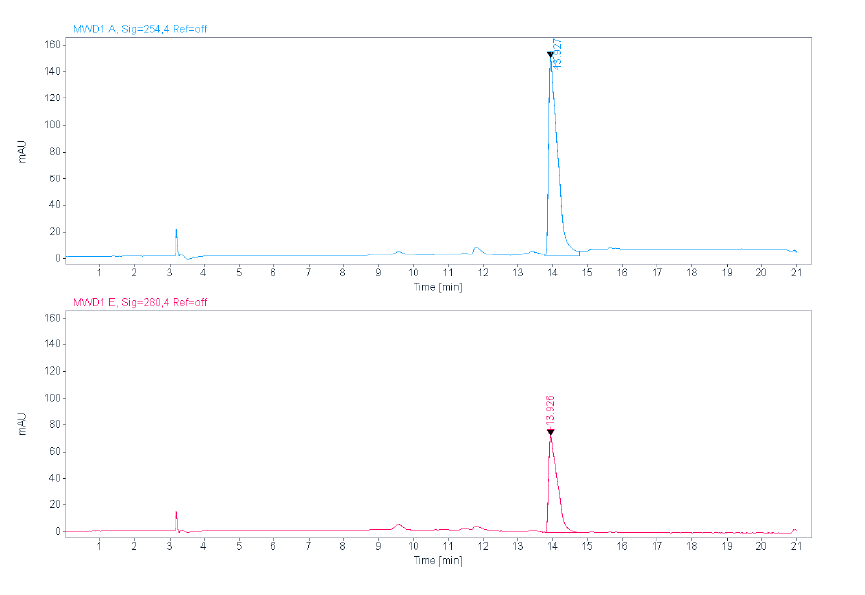
**HPLC:**

**ESI:**

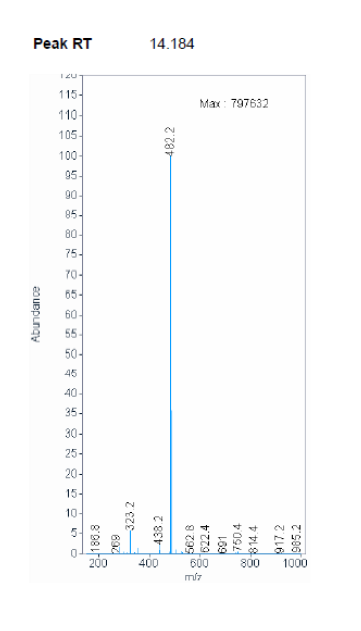

**HRMS:**

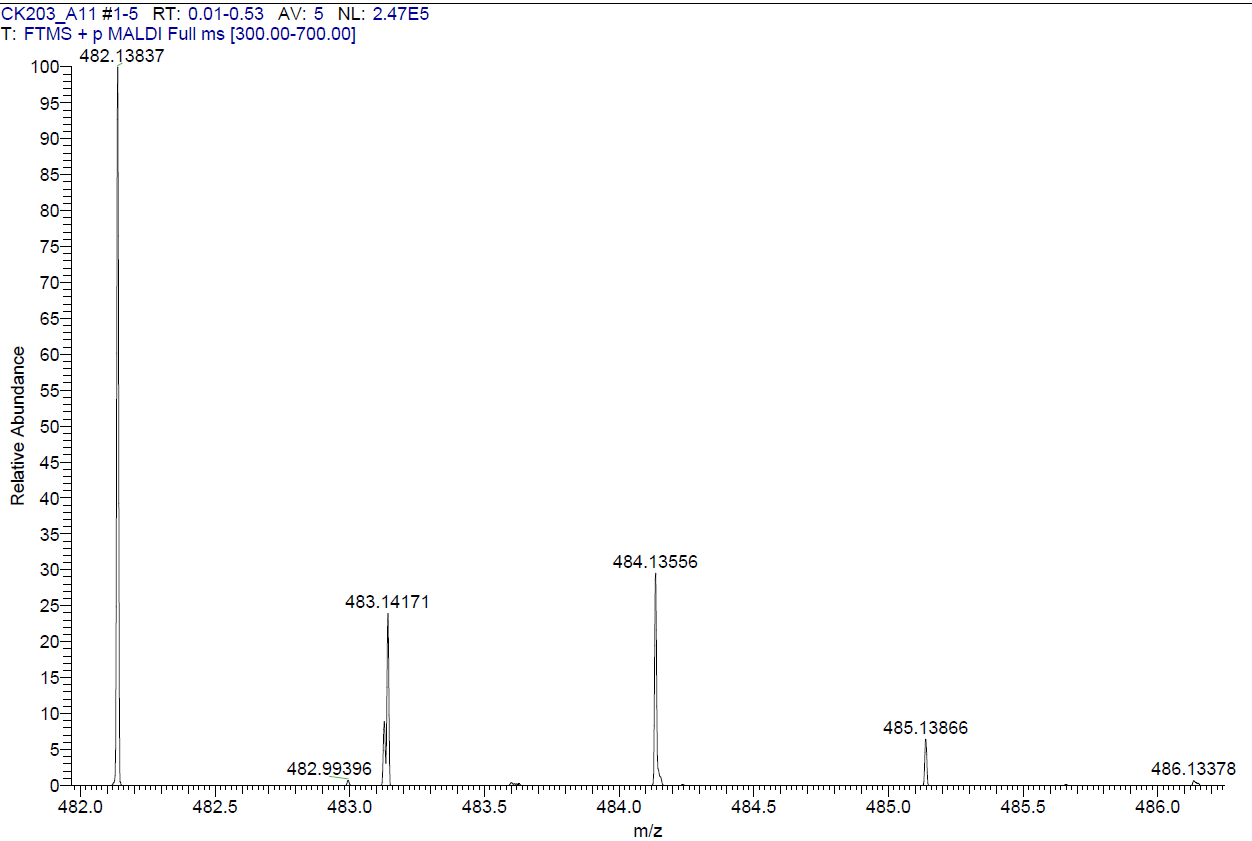

**Compound 24**

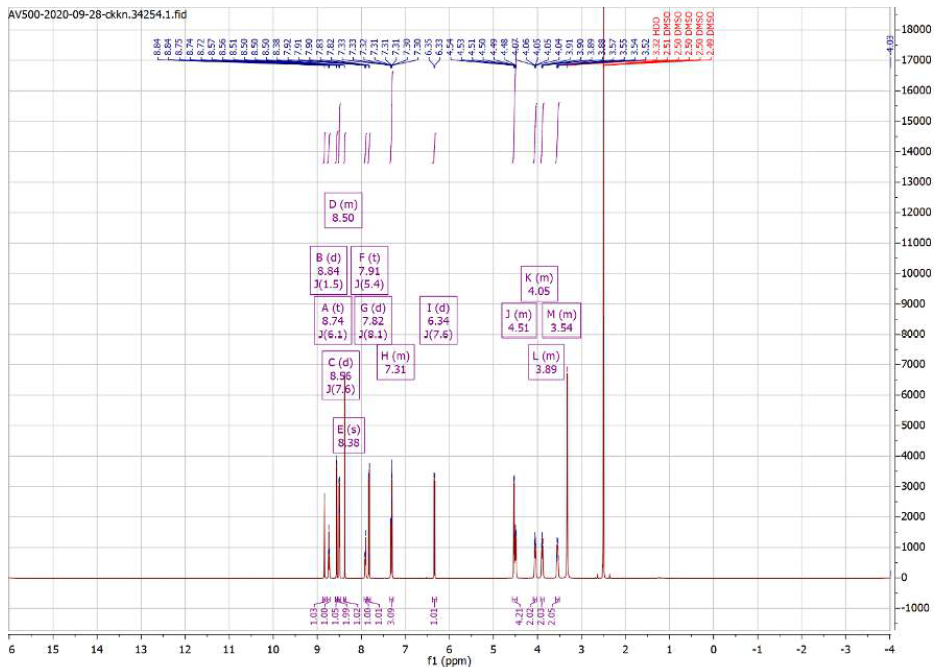
**^1^H NMR**

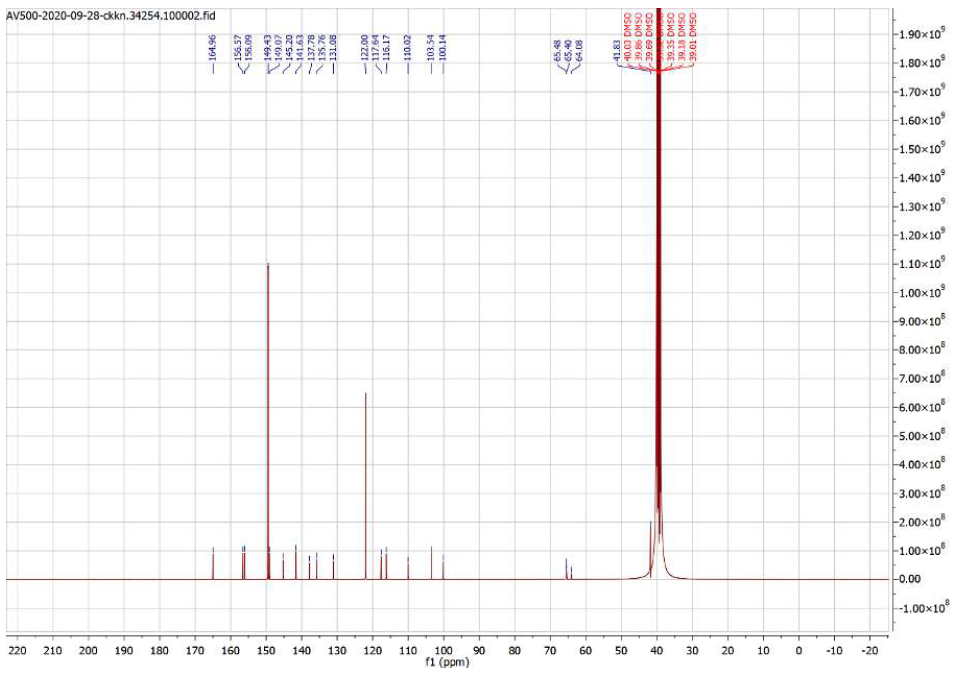
**^13^C NMR**

**HPLC:**

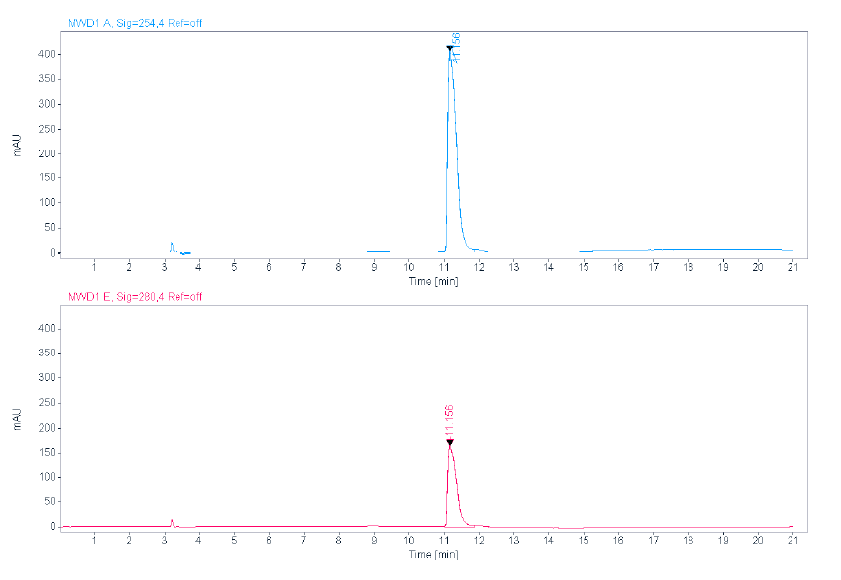

**ESI:**

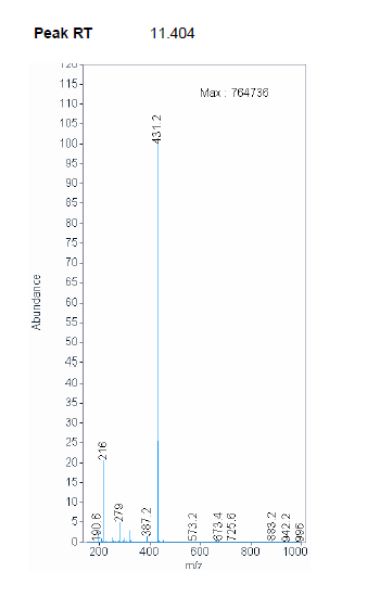

**HRMS:**

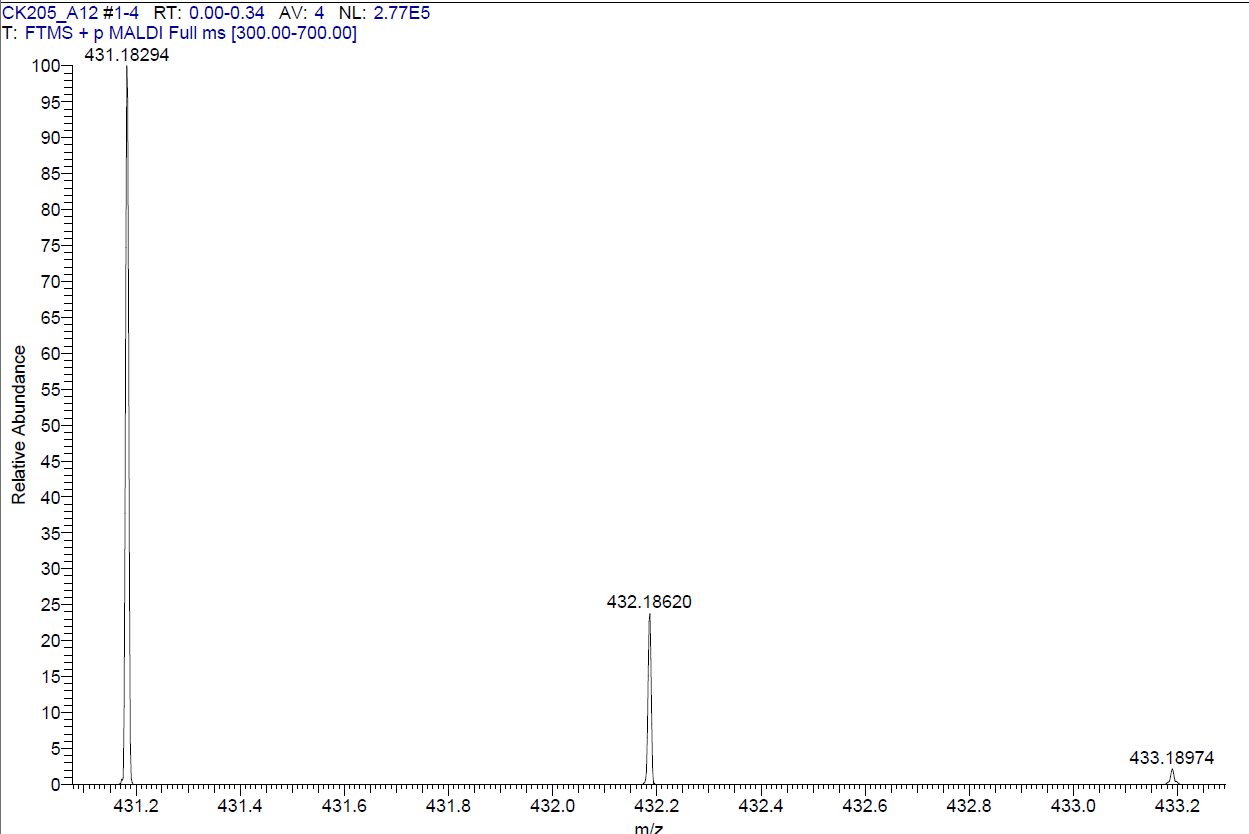

**Compound 25**

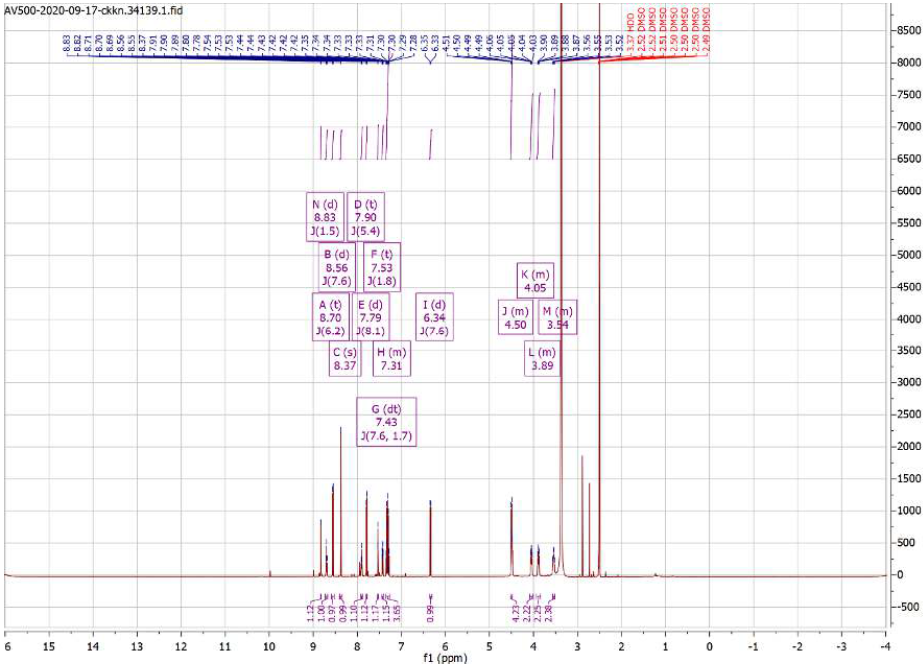
**^1^H NMR**

**^13^C NMR**

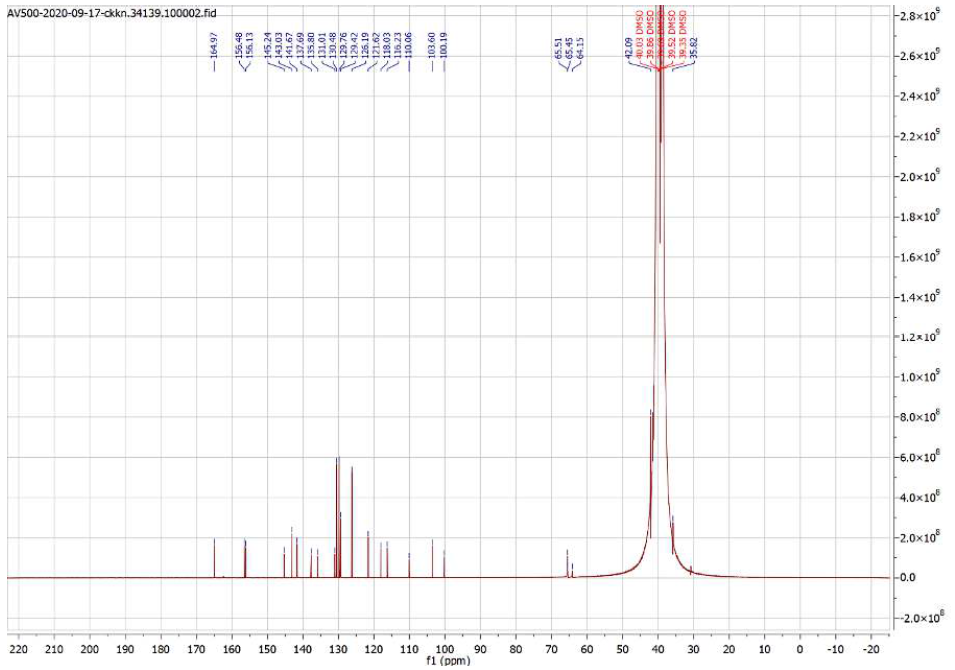

**HPLC:**

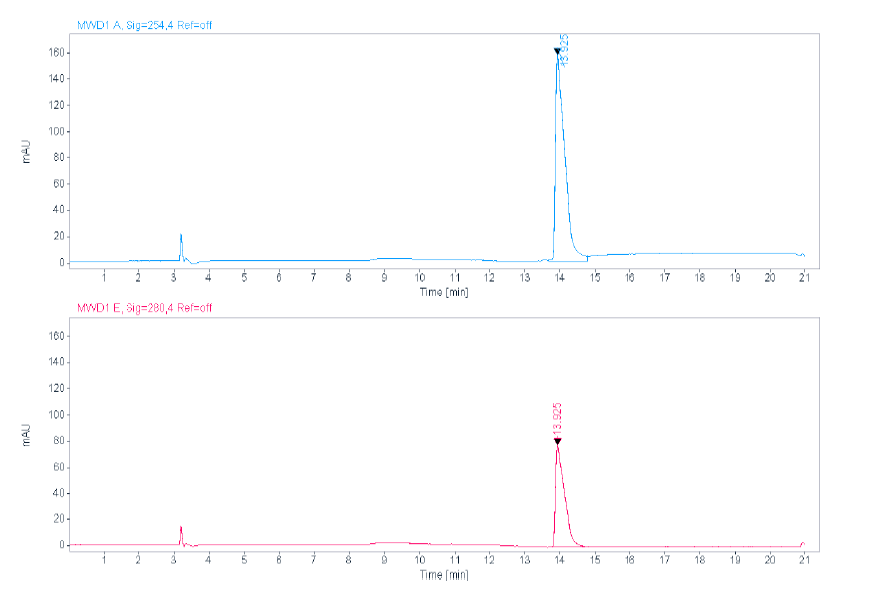

**ESI:**

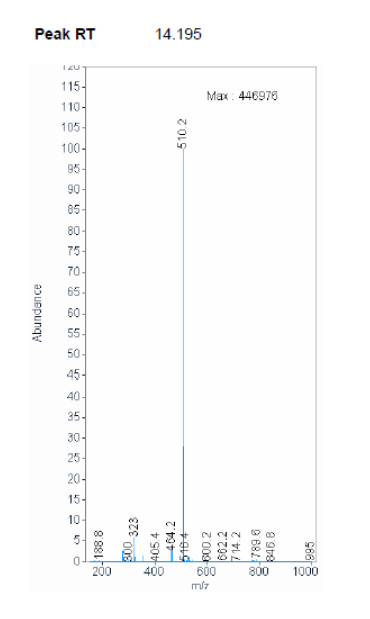

**HRMS:**

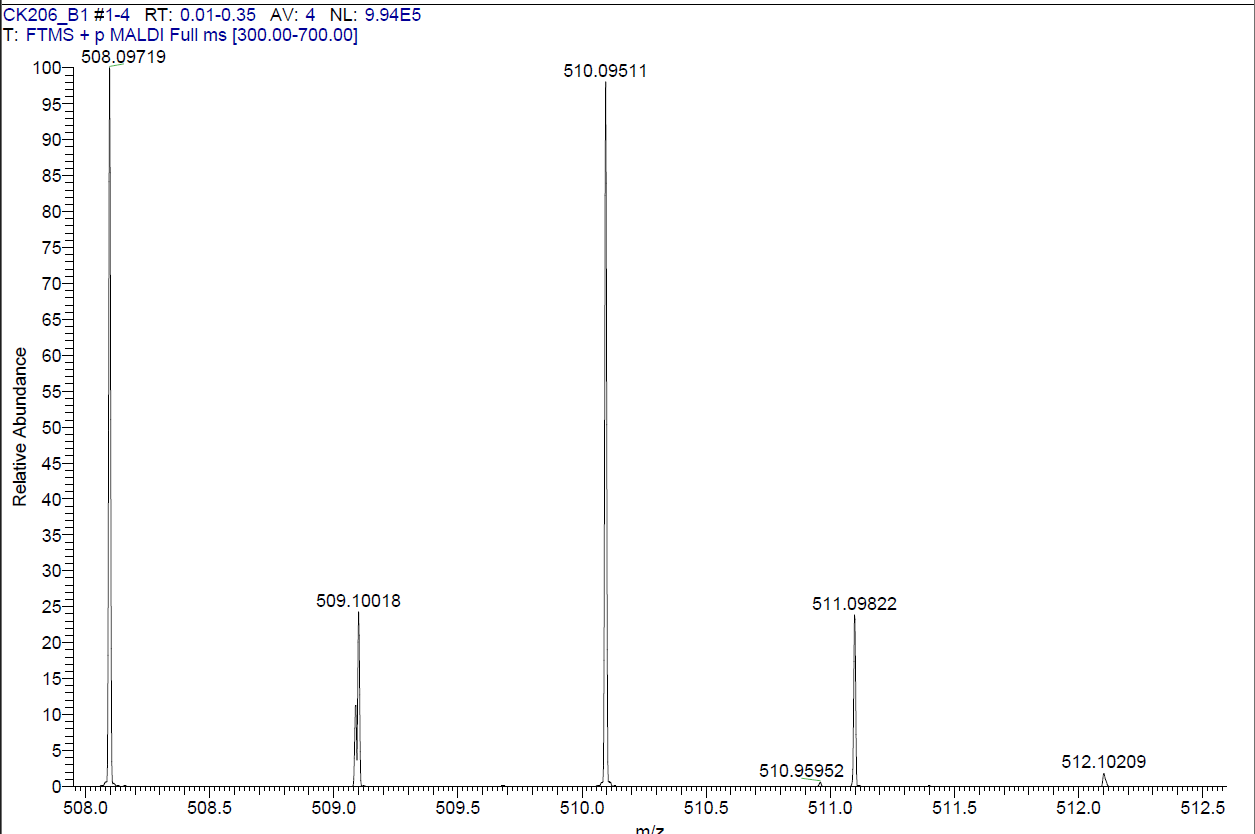

**Compound 26**

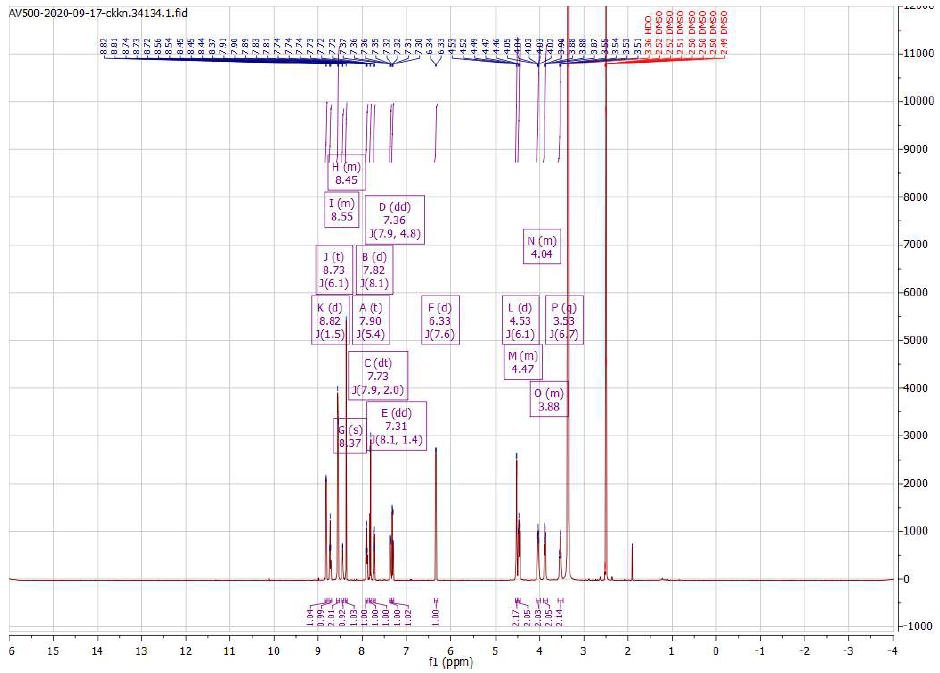
**^1^H NMR**

**^13^C NMR**

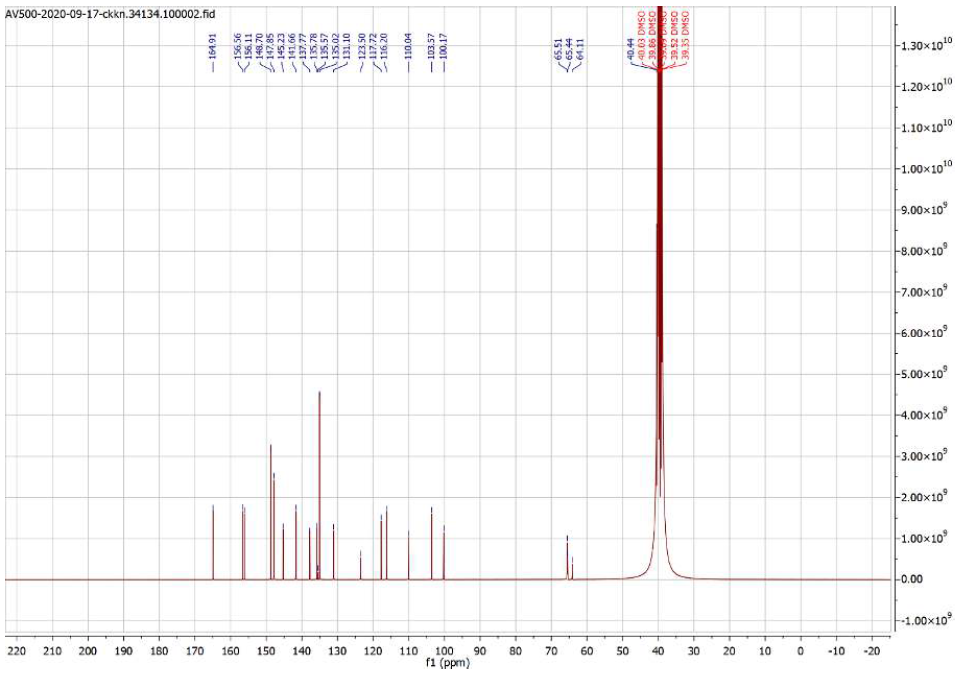

**HPLC:**

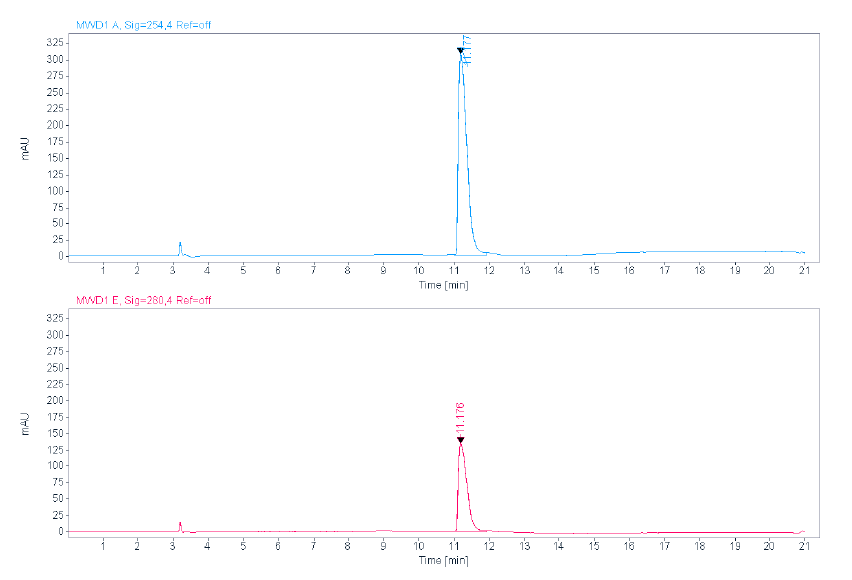

**ESI:**

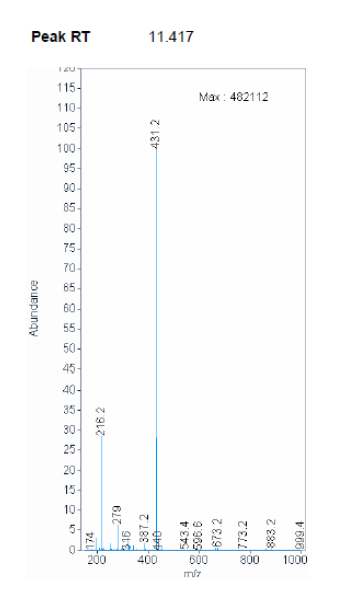

**HRMS:**

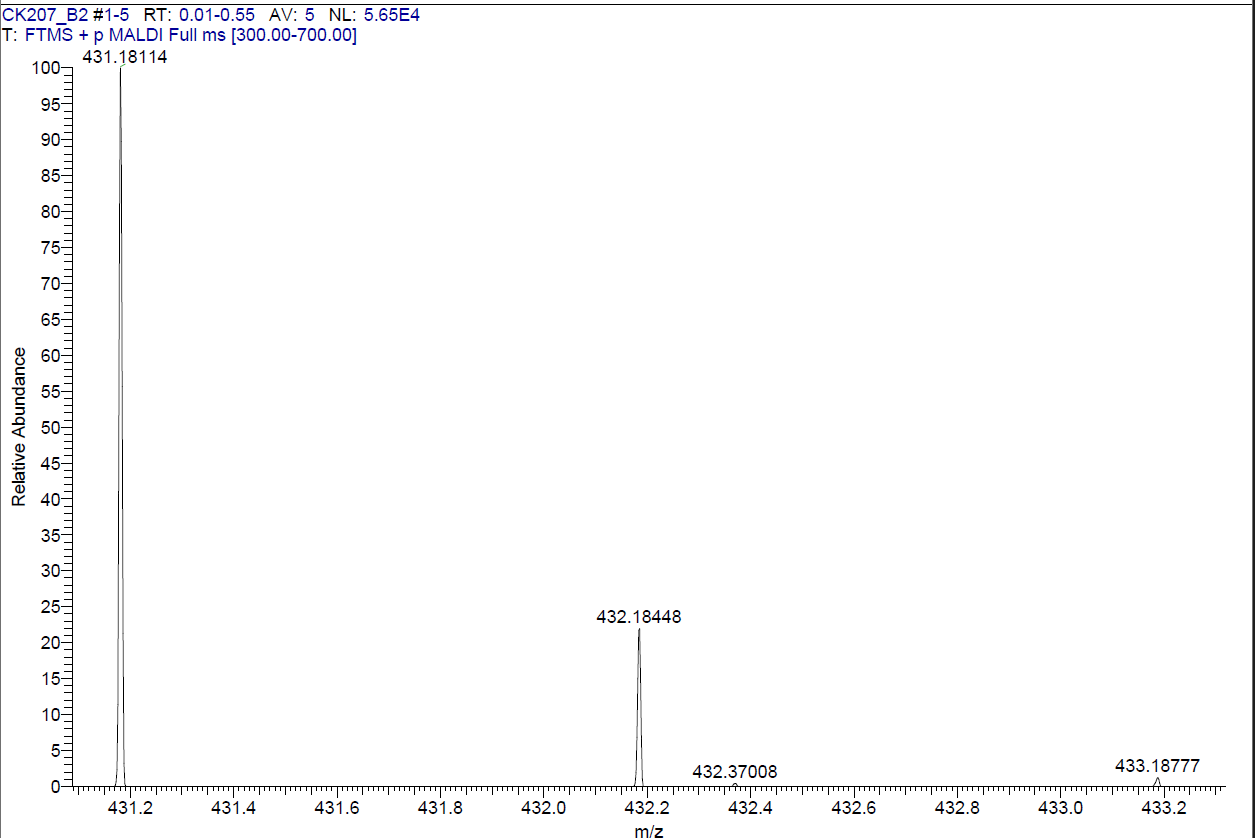

**Compound 27**

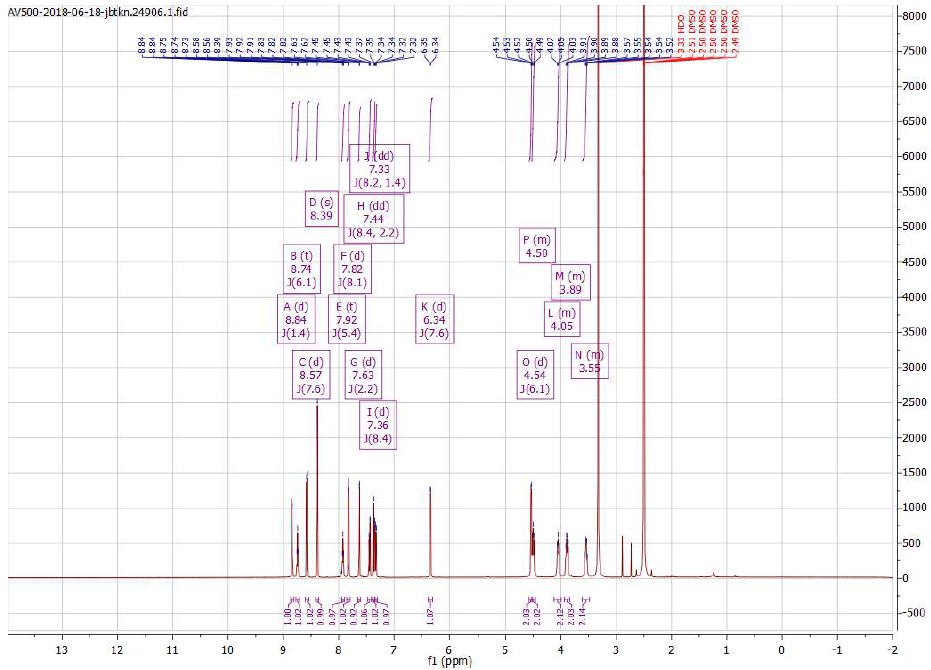
**^1^H NMR**

**^13^C NMR**

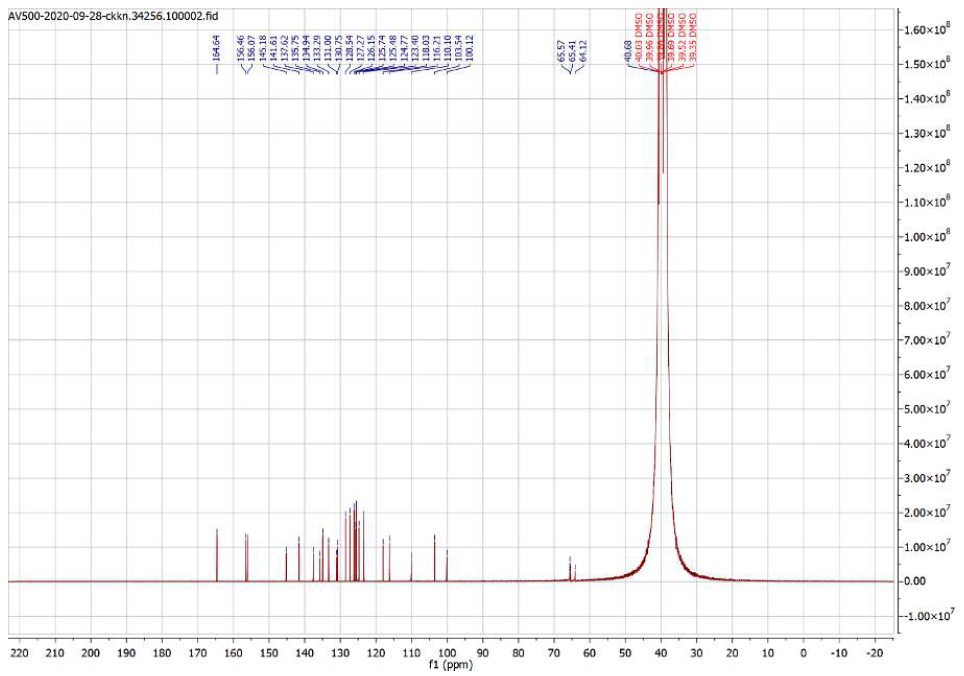

**HPLC:**

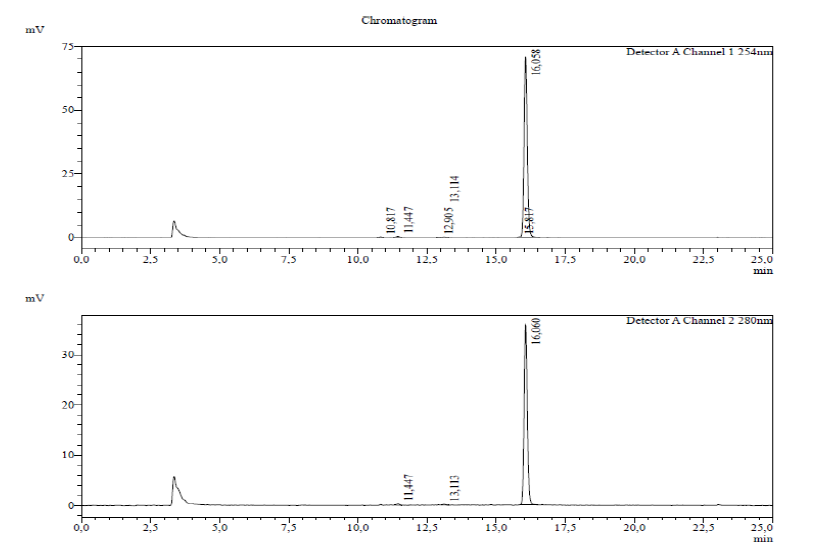

**ESI:**

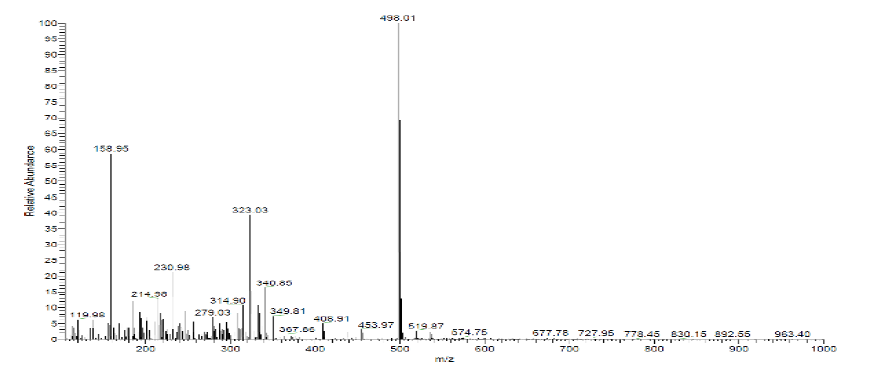

**HRMS:**

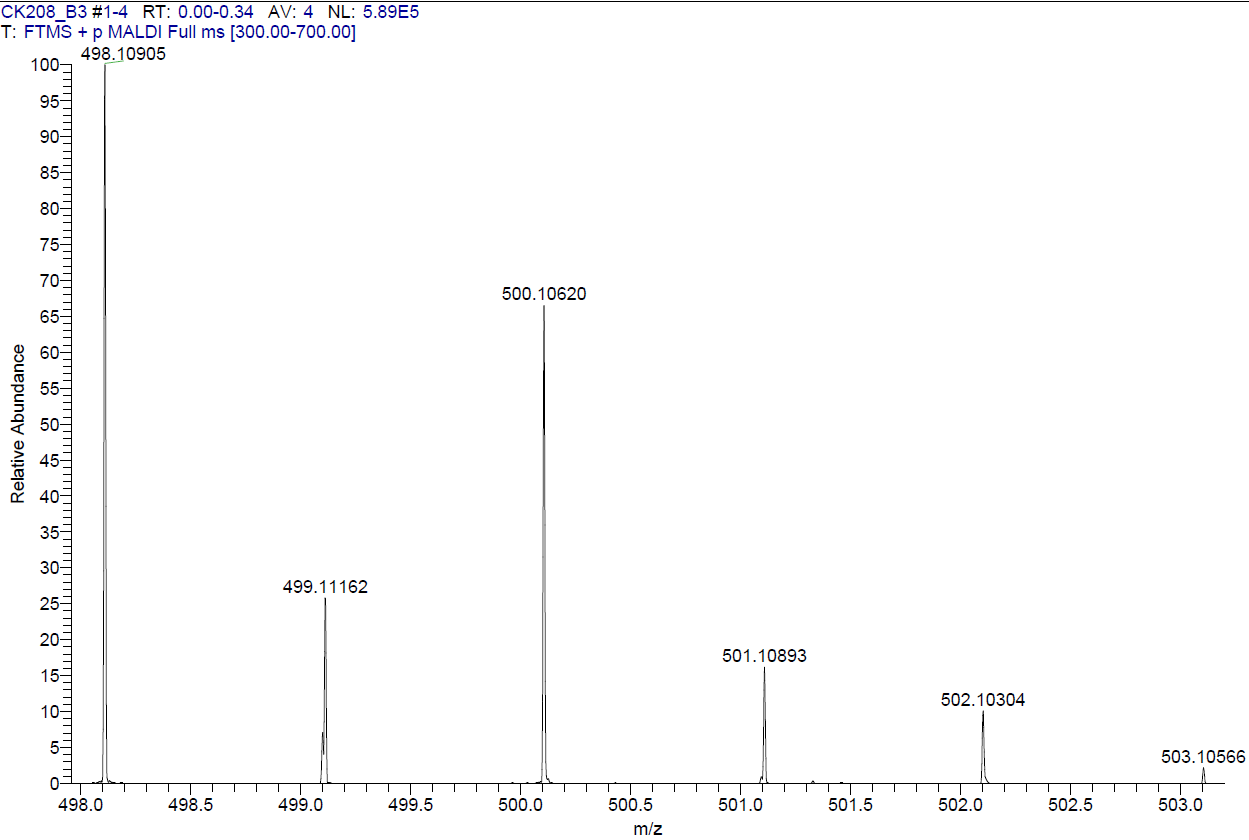

**Compound 28**

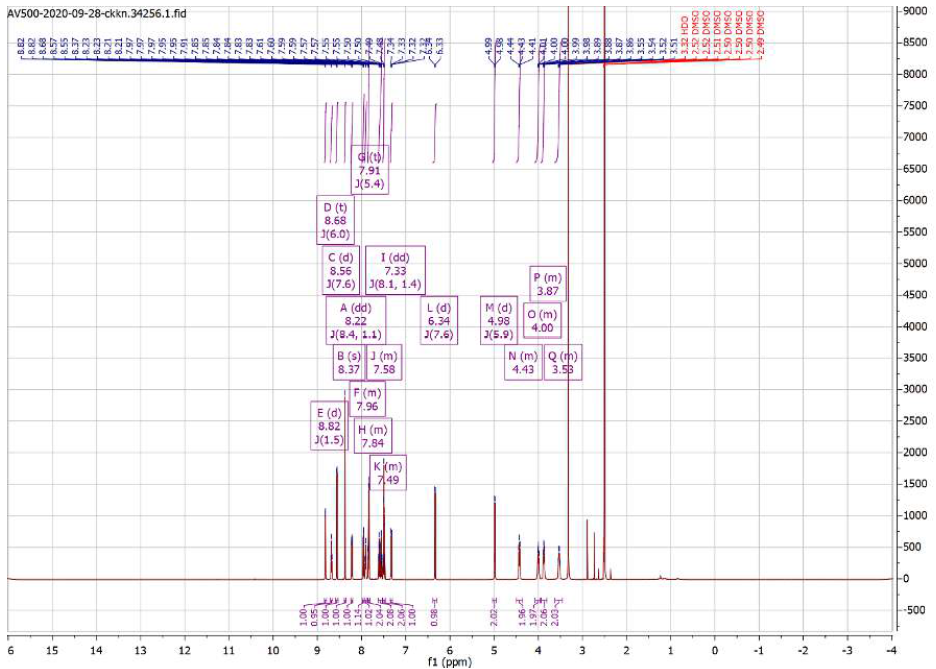
**^1^H NMR**

**^13^C NMR**

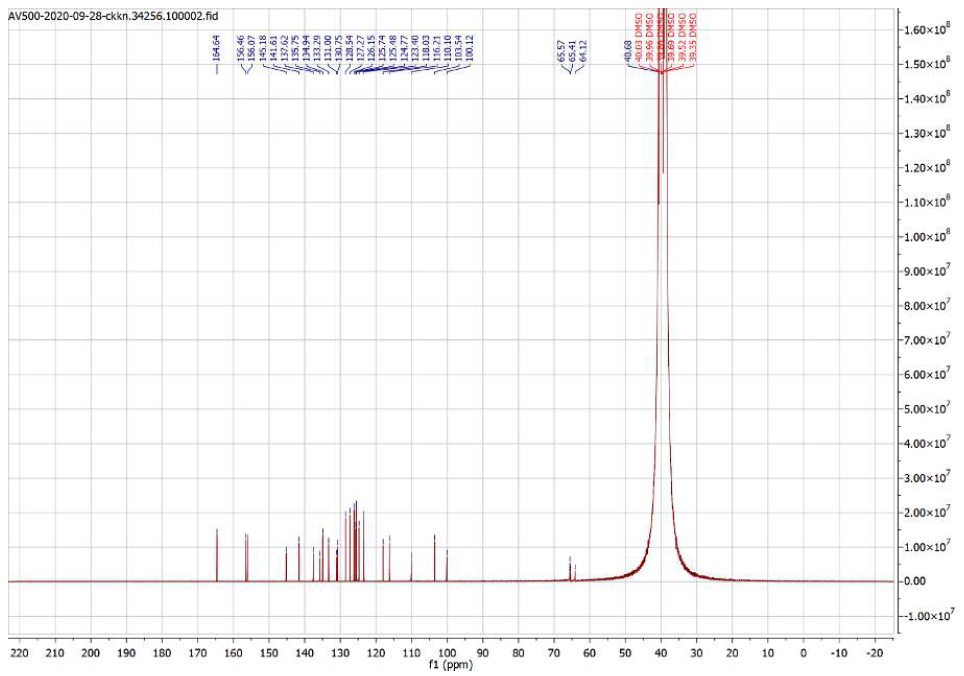

**HPLC:**

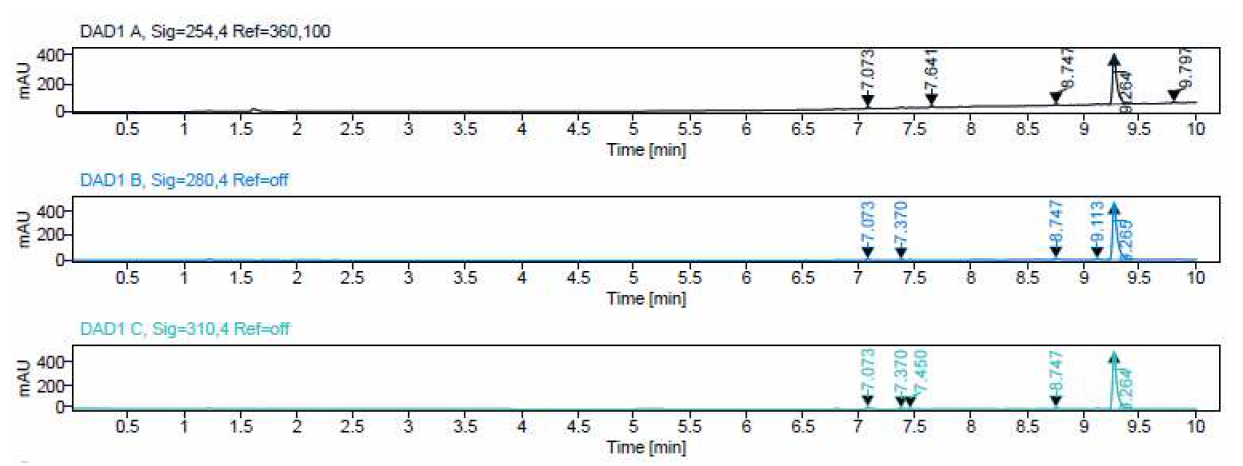

**ESI:**

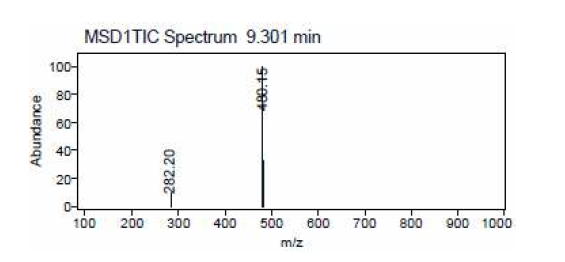

**HRMS:**

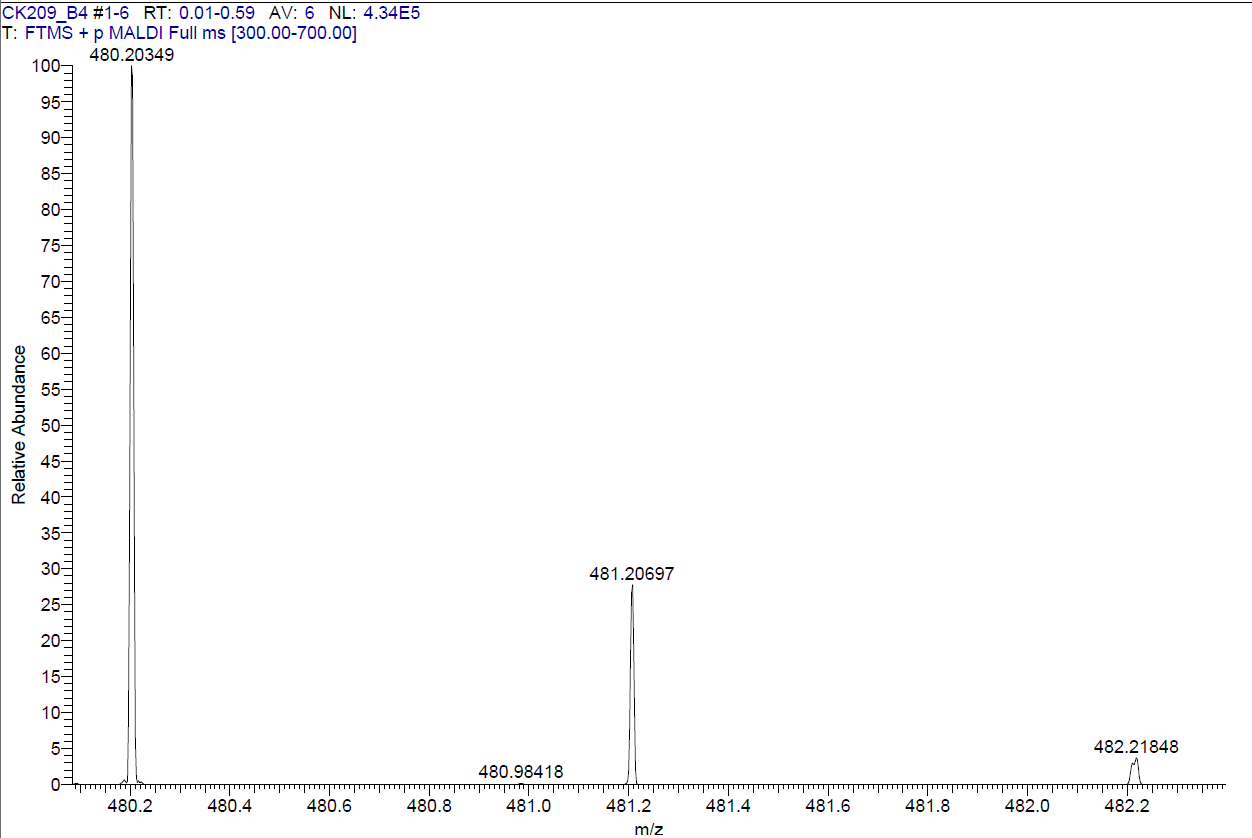

**Compound 29**

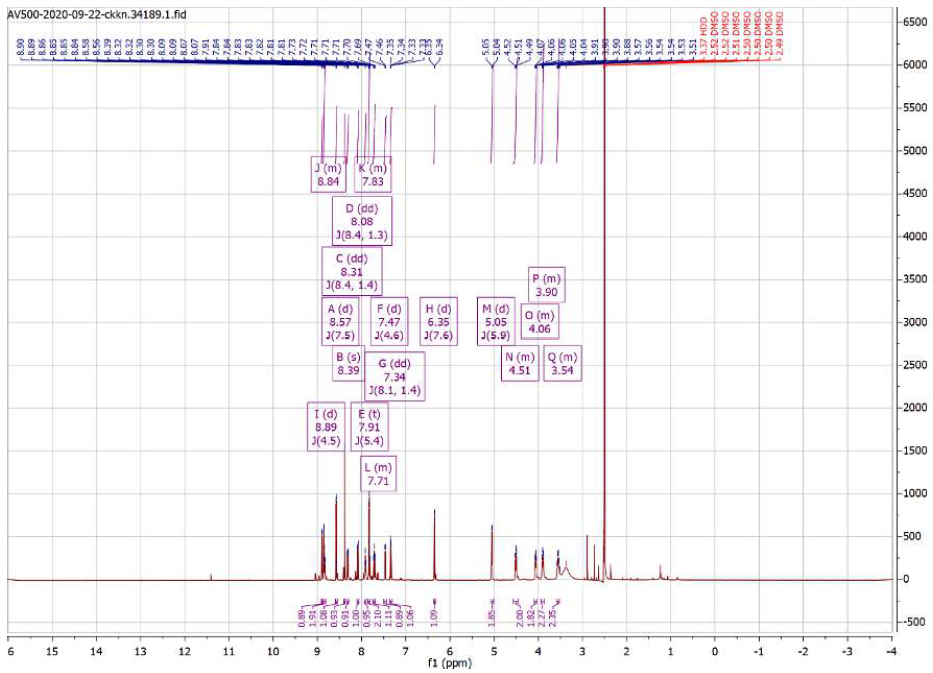
**^1^H NMR**

**^13^C NMR**

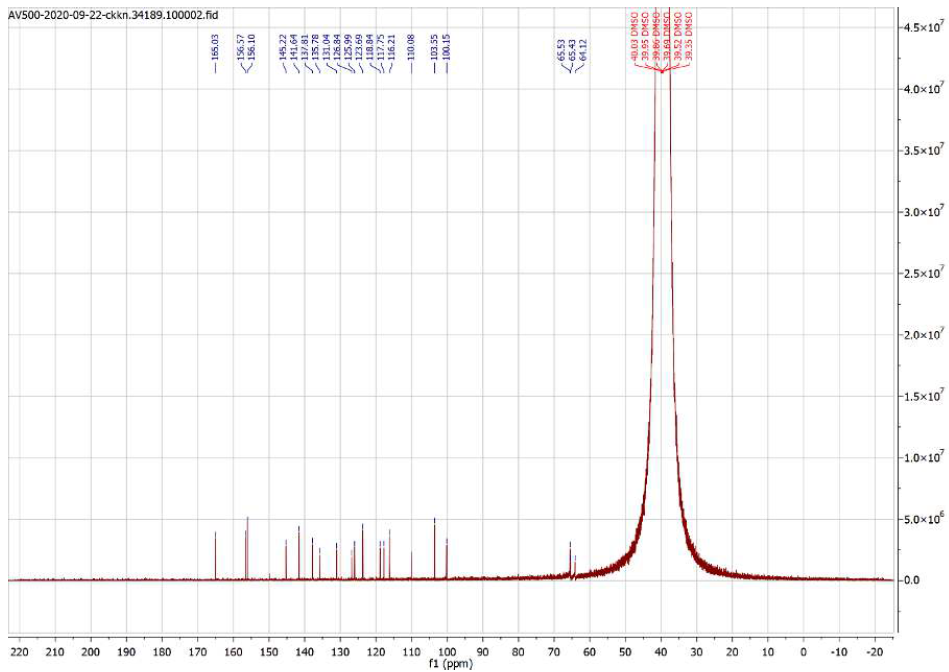

**HPLC:**

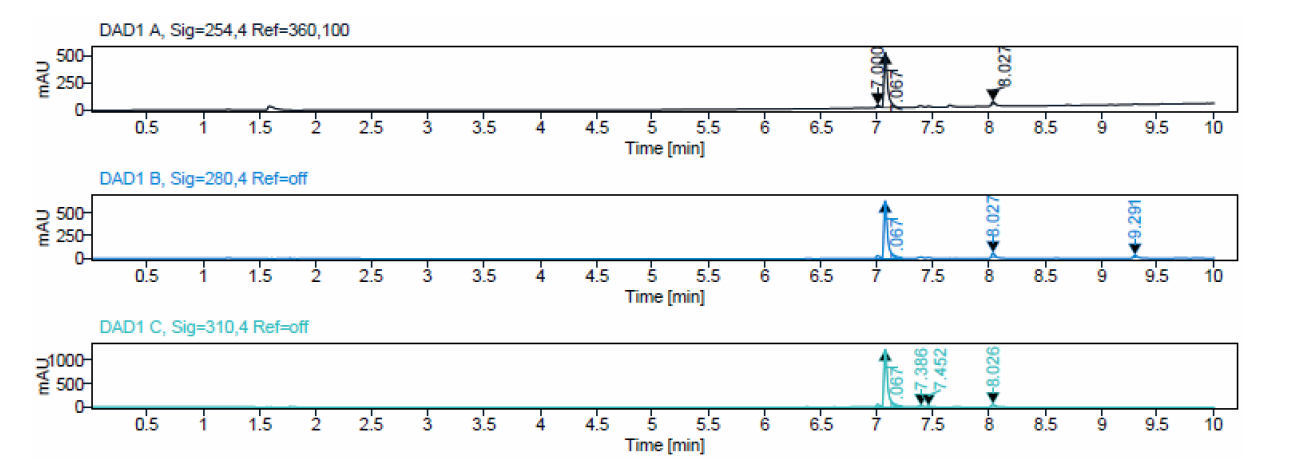

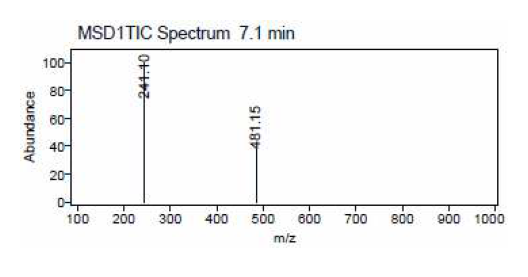
**ESI:**

**HRMS:**

**Compound 30**

**^1^H NMR**

**^13^C NMR**

**HPLC:**

**ESI:**

**HRMS:**
